# The Chlamydomonas Genome Project, version 6: reference assemblies for mating type *plus* and *minus* strains reveal extensive structural mutation in the laboratory

**DOI:** 10.1101/2022.06.16.496473

**Authors:** Rory J. Craig, Sean D. Gallaher, Shengqiang Shu, Patrice Salomé, Jerry W. Jenkins, Crysten E. Blaby-Haas, Samuel O. Purvine, Samuel O’Donnell, Kerrie Barry, Jane Grimwood, Daniela Strenkert, Janette Kropat, Chris Daum, Yuko Yoshinaga, David M. Goodstein, Olivier Vallon, Jeremy Schmutz, Sabeeha S. Merchant

## Abstract

Five versions of the *Chlamydomonas reinhardtii* reference genome have been produced over the last two decades. Here we present version 6, bringing significant advances in assembly quality and structural annotations. PacBio-based chromosome-level assemblies for two laboratory strains, CC-503 and CC-4532, provide resources for the *plus* and *minus* mating type alleles. We corrected major misassemblies in previous versions and validated our assemblies via linkage analyses. Contiguity increased over ten-fold and >80% of filled gaps are within genes. We used Iso-Seq and deep RNA-seq datasets to improve structural annotations, and updated gene symbols and textual annotation of functionally characterized genes via extensive curation. We discovered that the cell wall-less classical reference strain CC-503 exhibits genomic instability potentially caused by deletion of *RECQ3* helicase, with major structural mutations identified that affect >100 genes. We therefore present the CC-4532 assembly as the primary reference, although this strain also carries unique structural mutations and is experiencing rapid proliferation of a *Gypsy* retrotransposon. We expect all laboratory strains to harbor gene-disrupting mutations, which should be considered when interpreting and comparing experimental results across laboratories and over time. Collectively, the resources presented here herald a new era of Chlamydomonas genomics and will provide the foundation for continued research in this important reference.

## INTRODUCTION

The unicellular green alga Chlamydomonas (*Chlamydomonas reinhardtii*) is one of the primary model organisms in plant and cell biology. Chlamydomonas has been instrumental to discoveries in photosynthesis, chloroplast biology, and cilia structure and function, facilitated by its experimental tractability and amenability to classical genetics (Salomé and Merchant 2019). More recently, the species has been used as a powerful model for investigating the eukaryotic cell cycle (Cross and Umen 2015) and conserved mechanisms of sexual reproduction (Ning et al. 2013; Fédry et al. 2017), for discovery of optogenetic tools (Deisseroth and Hegemann 2017), and for in situ structural analyses by cryo-electron microscopy (Engel et al. 2015; Freeman Rosenzweig et al. 2017). Genome-wide mutant libraries form part of a growing suite of tools for exploiting high-throughput functional genomics approaches (Li et al. 2019; Fauser et al. 2022). As the most thoroughly studied green alga, Chlamydomonas also serves as an integral reference for the rapidly expanding fields of algal biology and biotechnology (Crozet et al. 2018; Blaby-Haas and Merchant 2019). The Chlamydomonas Genome Project was initiated two decades ago (Grossman et al. 2003; Merchant et al. 2007), and its continued development has kept the species at the forefront of plant and algal genomics (Blaby et al. 2014). Maintained at Phytozome (Goodstein et al. 2012), the genome assembly and structural annotations are a fundamental resource for contemporary Chlamydomonas research.

The Chlamydomonas genome is ∼111 Mb in length, GC-rich (∼64% genome-wide) and contains 17 chromosomes. Preceded by two preliminary versions (Grossman et al. 2003), the initial draft genome (v3) was assembled from ∼13x coverage of Sanger-sequenced reads (Merchant et al. 2007). Utilizing targeted sequencing of assembly gaps and molecular mapping data (Kathir et al. 2003; Rymarquis et al. 2005), the first chromosome-level assembly (v4) quickly followed in 2008 (Table 1). With the onset of next-generation sequencing, the v5 assembly was released in 2012 and applied both 454 and further Sanger sequencing to target all remaining gaps, successfully filling approximately half of those in v4 (Blaby et al. 2014). At 111.1 Mb, with 1,441 gaps (∼3.7% of the genome) and 37 unplaced scaffolds (∼2.0% of the genome), v5 has been the most long-standing release to date.

**Table 1.** Comparison of assembly metrics between v6 assemblies, previous reference genome versions and the CC-1690 assembly.

| Assembly strain/version | CC-503 v4 | CC-503 v5 | CC-503 v6 | CC-4532 v6 | CC-1690 |
| --- | --- | --- | --- | --- | --- |
| Year | 2008 | 2012 | 2022 | 2022 | 2020 |
| Technology | Sanger | Sanger + 454 | PacBio + Illumina | PacBio + Illumina | Nanopore + Illumina |
| Total length (Mb) | 112.3 | 111.1 | 111.5 | 114.0 | 111.1 |
| Unplaced scaffolds/contigs | 71 | 37 | 42 | 40 | 1 |
| Unplaced length (Mb) | 9.68 | 2.20 | 1.45 | 1.72 | 1.65 |
| Total contigs | 2,739 | 1,495 | 145 | 120 | 21 |
| Contig N50 (Mb) | 0.09 | 0.22 | 2.92 | 2.65 | 3.58 |
| GC (%) | 64.1 | 64.1 | 64.1 | 64.1 | 64.1 |
| Gaps/Ns (%) | 7.54 | 3.65 | 1.66 | 0.81 | <0.01 |
| Transposable elements (%) | 9.84 | 10.61 | 10.80 | 12.42 | 11.24 |
| Microsatellites (%) | 1.32 | 1.43 | 1.72 | 1.76 | 1.65 |
| Satellite DNA (%) | 3.33 | 3.68 | 4.79 | 5.25 | 5.09 |

Unplaced sequence was assembled as scaffolds in v4 and v5, and contigs in all other assemblies. The single unplaced contig in the released version of the CC-1690 assembly was later assembled to the right arm of chromosome 15 (Chaux-Jukic et al. 2021).

Although the assembly metrics of v5 represented a considerable achievement for the Chlamydomonas community, there remained substantial room for improvement relative to the highest quality Sanger-sequenced contemporaries. A decade earlier, near complete assemblies featuring just tens of gaps in the most repetitive regions had been produced for Arabidopsis (*Arabidopsis thaliana*) (Arabidopsis Genome Initiative 2000) and rice (*Oryza sativa*) (Goff et al. 2002). Recently, long-read sequencing technologies have provided a platform to achieve similar contiguity, and even assemble complete telomere-to-telomere assemblies, for far more complex genomes such as maize (*Zea mays*) (Jiao et al. 2017; Liu et al. 2020). Pacific Biosciences (PacBio) sequencing has been applied to close relatives of Chlamydomonas, yielding assemblies more contiguous than v5 for multiple unicellular and multicellular volvocine algae (Hamaji et al. 2018; Craig et al. 2021a; Yamamoto et al. 2021). Although none of these assemblies featured complete chromosomes, a chromosome-level assembly was achieved for the more distantly related alga *Chromochloris zofingiensis* by incorporating optical mapping (Roth et al. 2017). Most recently, O’Donnell et al. (2020) used ultra-long Nanopore sequencing (Liu et al. 2019) to produce an unannotated assembly of Chlamydomonas strain CC-1690 (classically named 21gr) featuring only four gaps, demonstrating that near complete Chlamydomonas assemblies are attainable with current technologies. It is noteworthy that many of the gaps in the v5 assembly are expected to be in genic regions (Tulin and Cross 2016), and improvements to contiguity are therefore expected to advance biological discovery via improved structural and functional annotation.

Perhaps of greater significance than contiguity, recent studies have highlighted inconsistencies between genetic mapping and the v5 assembly, potentially indicating misassemblies. Salomé and Merchant (2019) reported that the phytoene synthase gene (*PSY1*) is presently located on chromosome 2, although its corresponding white mutant *lts1* was mapped to chromosome 11 (McCarthy et al. 2004). Likewise, Ozawa et al. (2020) characterized *MTHI1,* which encodes an octotricopeptide repeat protein and is mutated in the non-photosynthetic strain *ac46*, observing that the gene is located on chromosome 17 despite having been mapped to chromosome 15 (Dutcher et al. 1991). Notably, both inconsistencies were introduced during the transition from v4 to v5, raising the possibility that past assembly improvements may have come at the expense of new errors.

There is also a potential issue with the classical reference strain, the cell wall-less CC-503 (*cw92*), which was chosen to meet the high DNA yield requirements of the early genome project. The *cw* phenotype was induced by mutagenesis of the mating type *plus* (*mt+*) “wild-type” strain 137c+ (later deposited as CC-125) with the methylating agent *N*-methyl-*N*’-nitro-*N*-nitrosoguanidine (MNNG) (Hyams and Davies 1972). MNNG primarily induces G:C to A:T transitions, although it can also induce double-strand breaks (DSBs) and chromosomal aberrations in high doses (Kaina 2004; Wyatt and Pittman 2006). For CC-503, the *cw* phenotype shows aberrant segregation in crosses, suggesting that there may be more than one causal mutation (Davies 1972; Hyams and Davies 1972). However, no causal mutations have been identified, and the potential genome-wide effects of mutagenesis in CC-503 have not been analyzed.

More broadly, a single strain does not represent the genomic diversity present among Chlamydomonas laboratory strains, which are thought to be derived from the haploid progeny of a single diploid zygospore isolated by G. M. Smith in 1945. The genomes of laboratory strains are comprised of two haplotypes, with any two strains carrying different haplotypes over up to ∼25% of their genome, but generally much less (Gallaher et al. 2015). The two haplotypes differ at ∼2% of sites, which is approximately equivalent to the average genetic diversity between any two field isolates from the same location (Craig et al. 2019) and consistent with the haplotypes having been inherited from one parental zygospore. Although many of the between-haplotype variants are expected to be functionally important, the mating type locus (*MT*) located on the left arm of chromosome 6 is the most variable region. The *plus* (*MT+*) and *minus* (*MT–*) alleles feature a small number of mating type-specific genes and several rearrangements that suppress crossover recombination (Ferris et al. 2010; De Hoff et al. 2013). While the CC-503 reference contains the *MT+* sequence, an *MT–* assembly is only available for the divergent field isolate CC-2290 (S1D2) (Ferris et al. 2010). Finally, all previous assembly versions have only included sequence and structural annotations for the nuclear genome, despite the relevance of organelle biology in the Chlamydomonas literature and the long availability of resources for the organelle genomes (Vahrenholz et al. 1993; Maul et al. 2002; Smith and Lee 2009; Gallaher et al. 2018).

Beyond the assembly itself, the structural annotations, which define the genomic coordinates of genes and the proteins they encode, are the foundation of ‘omics analyses, most notably high-throughput transcriptomics and proteomics. The Chlamydomonas structural annotations have also been subject to several rounds of improvement, utilizing advances in both sequencing technology and gene prediction algorithms (see Blaby et al. (2014) and Blaby and Blaby-Haas (2017)). Previous versions incorporated evidence from expressed sequence tags (ESTs) and assembled cDNAs, with protein homology support from *Volvox carteri* genes (Prochnik et al. 2010). The annotations performed for v5 made full use of next-generation sequencing with the incorporation of over one billion RNA-seq reads, resulting in several major changes to gene models (Blaby and Blaby-Haas 2017). The most recent v5 annotation (v5.6) features 17,741 protein-coding genes with 1,785 alternative transcripts. Recent advances in sequencing again provide substantial opportunities to update structural annotations. For example, Gallaher et al. (2021) used PacBio Iso-Seq (i.e. long-read sequencing of cDNA) to discover more than 100 polycistronic loci in Chlamydomonas, although these data have not yet been used to systematically improve structural annotations.

Here we present the first major update to both the Chlamydomonas assembly and annotation in nearly a decade. We present PacBio-based assemblies for the classical *mt+* reference strain CC-503 and for the *mt–* laboratory strain CC-4532, which we initially targeted to obtain an assembly of the *MT–* allele. These assemblies resulted in extensive improvements to both contiguity and annotation quality, which we validated against data from several independent sources. The assemblies resolved several large intra- and inter-chromosomal misassemblies present in v5. Comparative analyses revealed that the CC-503 genome harbors many large structural mutations predicted to affect ∼100 genes, while the genomes of all laboratory strains are likely to harbor a non-negligible and potentially highly variable number of unique transposable element (TE) insertions. We therefore present the CC-4532 assembly as the primary v6 reference genome and discuss the implications of mutation in the laboratory. These updates mark the start of an exciting new era for Chlamydomonas genomics, with developing opportunities to produce high-quality assemblies and annotations for several strains and divergent isolates of the species.

## RESULTS and DISCUSSION

### Version 6: a long-read Chlamydomonas reference assembly

As the first step in updating the reference genome, we produced de novo contig-level assemblies from high coverage (>120x) PacBio Sequel datasets for the *mt+* classical reference strain CC-503 and the *mt–* CC-4532. As presented below, CC-4532 is a putative subclone of CC-621 (NO–) and is partly descended from 137c*+*, although the exact crosses that produced the strain are unknown. In line with the reported inconsistencies with mapping data, we detected multiple contradictions between the prior v5 assembly and the v6 assembled contigs of both CC-503 and CC-4532. We thus reassembled all well-supported contigs to chromosomes without reference to previous versions, which we primarily achieved by mapping the contigs to the near complete Nanopore-based CC-1690 assembly (O’Donnell et al. 2020). This approach not only allowed contigs to be placed on chromosomes in a manner consistent across all three assemblies, but also enabled the estimation of gap lengths between remaining contig breaks in the PacBio assemblies relative to CC-1690. We validated all structural changes by reanalyzing previously published linkage data (Kathir et al. 2003; Liu et al. 2018). In addition, recent knowledge of centromeric (Lin et al. 2018; Craig et al. 2021a) and subtelomeric (Chaux-Jukic et al. 2021) repeats provided extrinsic support for the new chromosome assemblies. While the CC-4532 and CC-1690 assemblies are entirely consistent relative to each other and all supporting evidence, we identified remaining inconsistencies in the CC-503 assembly, indicative of genomic rearrangements unique to this strain. We describe these structural mutations further below, while the following text focuses on CC-4542 as the primary v6 reference assembly.

The final chromosome-level reference assembly, CC-4532 v6, is considerably more contiguous than previous versions (Table 1). The number of contigs decreased by an order of magnitude relative to v5, from 1,495 to 120, with a corresponding increase in the contig-level N50 from 0.22 Mb to 2.65 Mb (i.e. contigs ≥2.65 Mb represent >50% of the assembly length). Although unplaced sequence only fell from 2.20 Mb to 1.65 Mb, the 40 unplaced contigs in CC-4532 v6 are mostly unrelated to the 37 unplaced scaffolds in v5, all but three of which are now at least partially placed on chromosomes. The unplaced contigs from CC-4532 v6 are highly repetitive and generally feature sequences absent from previous versions. With a genome size of 114.0 Mb, CC-4532 v6 is ∼3 Mb larger than v5 and the CC-1690 assembly. This discrepancy can be explained in part by redundancy between the unplaced contigs and the gaps to which they presumably correspond, since gap lengths (represented by unknown bases i.e. Ns) were estimated relative to CC-1690. However, we attribute most of the biological increase in genome size to recent TE activity in the laboratory. In the following sections we present a thorough assessment of the assembly and annotation improvements.

### The version 6 assembly corrects misassemblies of version 5

The CC-4532 v6 assembly has major structural differences relative to v5, affecting the ordering and orientation of sequence both within and between chromosomes. Only six chromosomes (1, 4, 6, 7, 13 and 14) remained consistent with respect to the ordering of scaffolds in v5. The extent of the changes to the remaining 11 chromosomes ranged from minor intra-chromosomal reordering of short contigs to major inter-chromosomal rearrangements affecting megabases of sequence. An overview of the between-chromosome changes is presented in Figure 1A.

**Figure 1.**
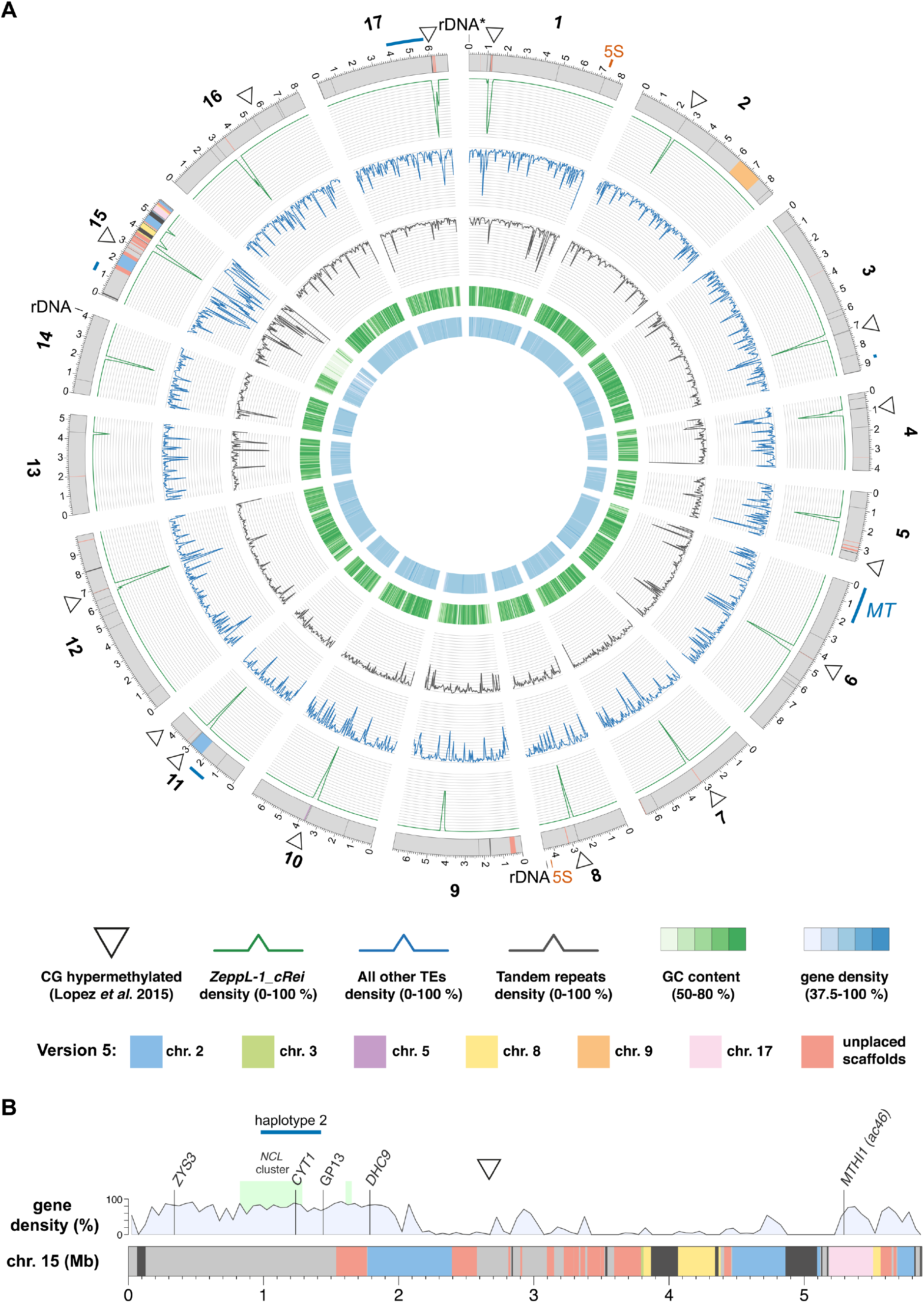
The CC-4532 version 6 assembly. **(A)** Circos plot (Krzywinski et al. 2009) representation of CC-4532 v6. Gray outer bands represent chromosomes, with colors highlighting genomic regions that were assembled on other chromosomes or unplaced scaffolds in v5. Dark gray regions represent gaps between contigs, with any gaps <10 kb increased to 10 kb to aid visualization. Outer lines in dark blue represent haplotype 2 regions, including the mating type locus (*MT*) and flanking regions on chromosome 6. All metrics were calculated for 50 kb windows. Tandem repeats include microsatellite and satellite annotations. CG hypermethylated regions were taken from Lopez et al. (2015) and mapped from v5 to v6 coordinates, with some neighboring regions merged to a single marker in the plot (see Supplemental Figure S3 for all regions). **(B)** Linear representation of chromosome 15. Colors are as in **(A)**, with dark gray representing assembly gaps. Marker genes are from Kathir et al. (2003) and the light green boxes represent the *NCL* gene clusters described by Boulouis et al. (2015). *CYT1* was previously recorded as *CYTC1*.

Many of the changes occurred in proximity to the most repetitive genomic regions, particularly the putative centromeres and the subtelomeres, as well as regions corresponding to unplaced scaffolds in v5. Although approximate centromeric locations were known from molecular mapping (Preuss and Mets 2002), genomic coordinates and sequence characteristics have only recently been reported. Lin et al. (2018) identified 200-800 kb regions tightly linked to the centromeres that featured multiple open reading frames (ORFs) encoding proteins with reverse transcriptase domains. Craig et al. (2021a) linked these ORFs to multiple copies of an *L1* LINE retrotransposon homologous to *Zepp*, the major centromeric component of the trebouxiophyte alga *Coccomyxa subellipsoidea* (Blanc et al. 2012). Termed *Zepp-*like (*ZeppL*) elements in Chlamydomonas, this TE forms highly localized clusters at the putative centromeres, although in v5 chromosomes 2, 3, 5 and 8 featured two clusters, and chromosomes 11 and 15 lacked clusters (Lin et al. 2018; Craig et al. 2021a). Chlamydomonas subtelomeres were recently shown to feature large satellite arrays termed *Sultans*, with other complex repeats present at specific chromosome termini (Chaux-Jukic et al. 2021). Subtelomeres are capped by the telomeric repeat (TTTTAGGG)_n_ (Petracek et al. 1990). Due to their complexity, subtelomeres were previously poorly assembled, and only half of chromosome termini featured a scaffold terminating in telomeric repeats in v5.

Comparisons of chromosomes 5 (Figure 2A) and 11 (Figure 2B) between v5 and v6 illustrate the types of misassemblies that affected these regions. In v5, the left arm of chromosome 5 terminated in a 47 kb contig featuring a *ZeppL* cluster (purple, Figure 2A), which in v6 is assembled within the putative centromere of chromosome 10 (Supplemental Figure 1D). The remaining regions of chromosome 5, consisting of three blocks of ∼0.7, 1.2 and 1.7 Mb (light blue, yellow and orange, respectively), are now rearranged and reorientated. The misassembly of the light blue and yellow regions featured a large gap corresponding to part of scaffold 24 (containing *MUT6*), while the misassembly of the yellow and orange regions featured subtelomeric repeats that are now correctly placed at the left arm terminus in v6. Thus, the reassembled chromosome 5 features a single internal centromere, subtelomeric repeats at both termini, and is congruent with the molecular map (Kathir et al. 2003). On chromosome 11, the movement of an ∼750 kb region (orange) from chromosome 2 simultaneously resolved the absence of a putative centromere on chromosome 11 and the presence of two *ZeppL* clusters on chromosome 2 (Figure 2B). This region includes *PSY1*, which was mapped genetically to chromosome 11 (McCarthy et al. 2004; Salomé and Merchant 2019). Independently, an ∼860 kb region (light blue) was inverted, consistent with the tight linkage of *PETC1* and *DLE2* (Kathir et al. 2003). Misassemblies affecting other chromosomes are shown in Supplemental Figure S1.

**Figure 2.**
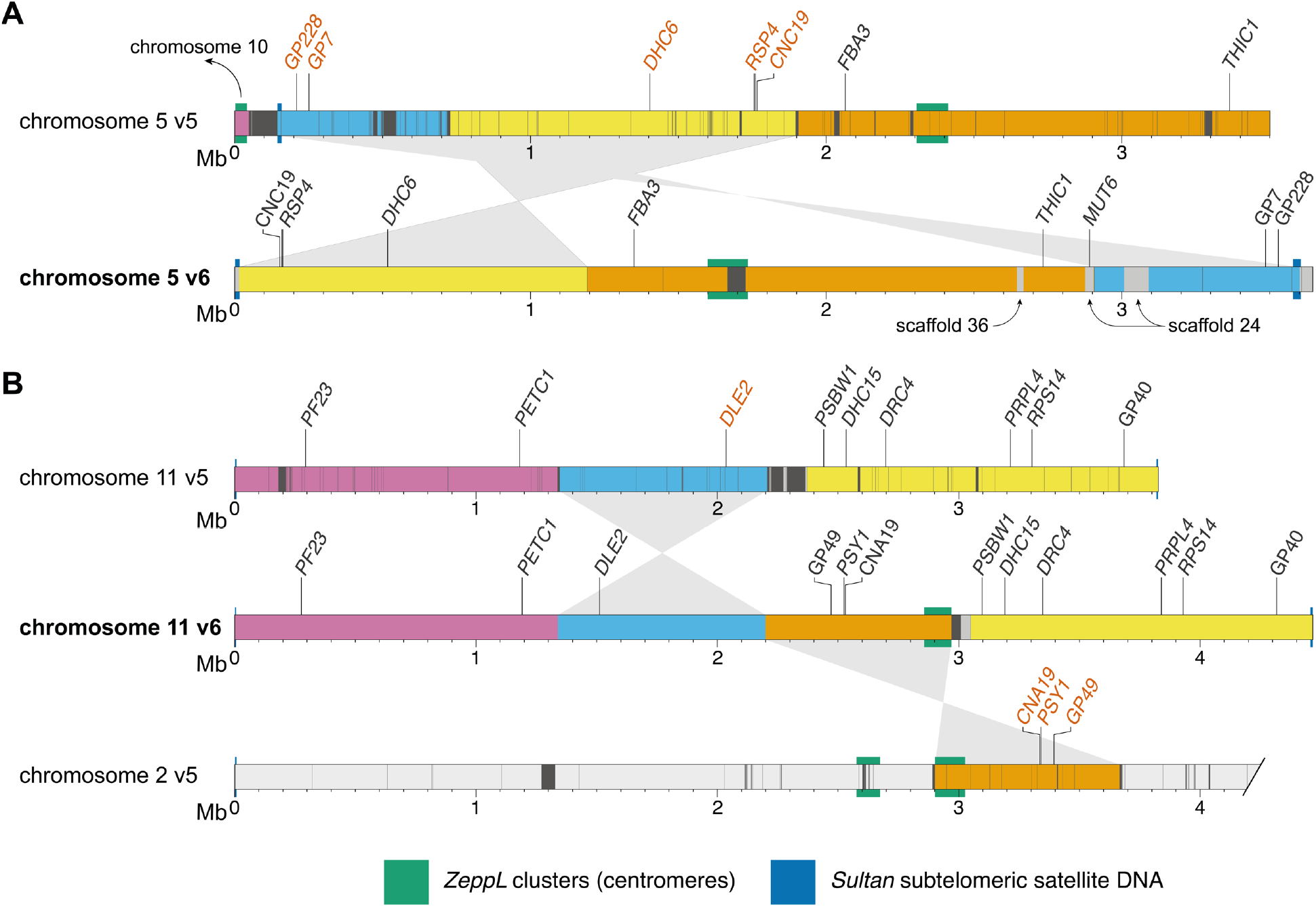
Version 5 misassemblies and their resolution in version 6. Chromosome segments are colored to show the reordering and reorientation of specific regions, and dark gray regions represent assembly gaps. Markers inconsistent with the molecular map of Kathir et al. (2003) are shown in vermillion text. Gene symbols (in italics) were updated where applicable. Note that the plot was made using CC-503 v6 to simplify mapping between versions. CC-503 v6 and CC-4532 v6 are entirely syntenic for chromosomes 5 and 11. **(A)** Reassembly of chromosome 5. The purple region was reassigned to chromosome 10. Light gray regions on the v6 chromosome correspond to sequence not assembled on the v5 chromosome (e.g. the region containing *MUT6* corresponds to part of scaffold 24 in v5). In the original map *RSP4* corresponded to the *pf1* marker (and the neighboring *RSP6* to *pf26*, not shown) (Dutcher 2014). Updated gene symbols: *FBA3* was *ALD*, *THIC1* was *THI8*. **(B)** Reassembly of chromosome 11; only the first 4.2 Mb of chromosome 2 is shown. Genes that originally corresponded to genetic markers are: *PSY1* – *lts1*, *PF23* – *pf23* (Yamamoto et al. 2017), *DRC4 – pf2* (Dutcher 2014), *PRPL4 – ery1, RPS14 – cry1*. Updated gene symbols: *PETC1* was *PETC*, *DLE2* was *VFL2*, *DHC15* was *ODA2*, *PSBW1* was *PSBW*.

By far the most substantial changes affected chromosome 15, which approximately tripled in length from 1.92 Mb in v5 (the shortest chromosome) to 5.87 Mb in CC-4532 v6, acquiring sequence previously assigned to chromosomes 2, 3, 8 and 17, as well as 17 unplaced scaffolds (Figure 1B). The sequence reassembled from chromosomes 2 (∼1.2 Mb) and 17 (∼0.3 Mb) each featured a marker gene previously mapped to chromosome 15: *DHC9* (Porter et al. 1996; Kathir et al. 2003) and the aforementioned *MTHI1* (Dutcher et al. 1991; Ozawa et al. 2020), respectively. Some of the sequence reassembled from chromosome 8 (∼0.4 Mb) and unplaced scaffolds (∼1.1 Mb total) featured *ZeppL* elements, explaining the absence of centromeric repeats on chromosome 15 in v5. We attribute the degree of past misassembly to the unique sequence characteristics of chromosome 15. Its repeat content (47.2%) is substantially higher, and its gene density lower (36.7%), than the remaining 16 chromosomes (mean 17.7% and 79.0%, respectively) (Supplemental Table S1). Furthermore, this pattern is not uniform: the gene density of the chromosome arms (67.1%, ∼2.1 Mb left and ∼0.6 Mb right) approaches that of other chromosomes, while the internal region is massively repetitive (66.7%) and gene-poor (10.9%). As a result, chromosome 15 remains the most fragmented in CC-4532 v6, featuring 10 gaps spanning 9.2% of the chromosome length, relative to a mean of three gaps and 0.4% for the remaining chromosomes. We expect that many of the unplaced contigs may in fact belong on chromosome 15, although their extreme repeat content (69.8%) hinders efforts to place them without longer reads.

The unusual features of chromosome 15 raise questions about its evolutionary origins, gene content and chromosomal environment. Except for *MTHI1*, all marker genes (*ZYS3*, *CYT1* and *DHC9*) are located within the relatively gene-rich left arm of the chromosome. This region is also notable for containing almost all the *NCL* (*NUCLEAR CONTROL OF CHLOROPLAST GENE EXPRESSION-LIKE*) genes, encoding a family of RNA binding proteins that are experiencing ongoing diversification in Chlamydomonas (Boulouis et al. 2015). All but one of the 49 *NCL* genes are on chromosome 15, with 43 present in a cluster spanning ∼460 kb, and three forming a shorter upstream cluster that was assembled on scaffold 19 in v5 (Figure 1B). The mutation responsible for the *yellow-in-the-dark* mutant *y1* was also mapped to the left arm of chromosome 15 and is linked to *DHC9* (Porter et al. 1996). The unknown *Y1* gene might thus have been assigned to either chromosome 2 or an unplaced scaffold in v5, and it may be worth revisiting attempts to clone the gene. The remainder of chromosome 15 contains only 145 genes, 80 of which are in the highly repetitive internal region. Although most of these genes are not functionally annotated, we expect at least some to be essential (e.g. the plastid 50s ribosomal protein gene *PRPL3*). It would be interesting to determine if much of chromosome 15 is heterochromatic, and if so, whether genes are expressed from heterochromatic environments (e.g. as is the case for many genes on the repeat-rich dot chromosome in *Drosophila melanogaster* (Riddle and Elgin 2018)). Similarly, it would be interesting to explore whether the high repeat content results in an atypical recombination landscape on chromosome 15.

### Linkage data validates the CC-1690 and version 6 assemblies

As presented above, many of the misassembly corrections are supported by prior genetic and molecular mapping. To systematically validate the improvements between v5 and v6, we turned to two independent genetic recombination datasets to determine linkage between sets of markers. We primarily compared v5 to the CC-1690 assembly, since CC-1690 was used as a reference to scaffold the v6 assemblies, and the CC-1690 and CC-4532 v6 assemblies are entirely syntenic. We repeated these analyses using CC-503 v6 following the discovery of outstanding inconsistencies in this assembly. We first identified the v5 chromosomal coordinates of 239 molecular markers described by Kathir et al. (2003). We then ordered the genotype data used to generate the genetic map based on the v5 coordinates before estimating a new genetic map with the R/QTL package (Broman et al. 2003). To assess the concordance between assigned and true genomic positions, we visualized recombination frequencies between marker pairs: two unlinked markers should exhibit random segregation and appear as dark blue squares (low log of the odds [LOD] score), whereas linked markers should appear in yellow. While most markers agreed with their v5 chromosomal locations, we identified 10 misplaced markers, one each on chromosomes 4 and 14, with the remaining eight mapping to chromosomes 2 or 9 (Figure 3A, Supplemental Figure S2). Markers CNA19 and GP49 were located on chromosome 2 in v5, but showed strong linkage with chromosome 11. Satisfyingly, both markers relocated to chromosome 11 in CC-1690 (Figure 3B) and subsequently in both v6 assemblies (Figure 2B). We also resolved the genomic location of most other mismapped markers when using CC-1690 coordinates. Yet, we noted that using CC-503 v6 coordinates did not fix issues with the remaining misplaced markers located on chromosomes 2 or 9 (Figure 3C). As detailed below, this mismapping stems from a putative chromosomal rearrangement unique to CC-503. Two markers remained apparently wrongly assigned when using CC-1690 or CC-503 v6 coordinates: GP332 and *ODA16*, which were assigned to the top of chromosome 14 and 4, respectively, in both assemblies. The genetic mapping data of Kathir et al. (2003) indicated strong linkage between GP332 and chromosome 7 markers (CNC43, *CHL27A* and *GLTR1*), and between *ODA16* and chromosome 5 markers (*DHC6*, CNC19 and *RSP6* – see Figure 2A). In both cases, the chance of the regions corresponding to these sequences being misassembled in the exact same location on independent contigs in CC-1690, CC-4532 v6 and CC-503 v6 is negligible, and their previous mapping locations or associated sequences are presumably incorrect.

**Figure 3.**
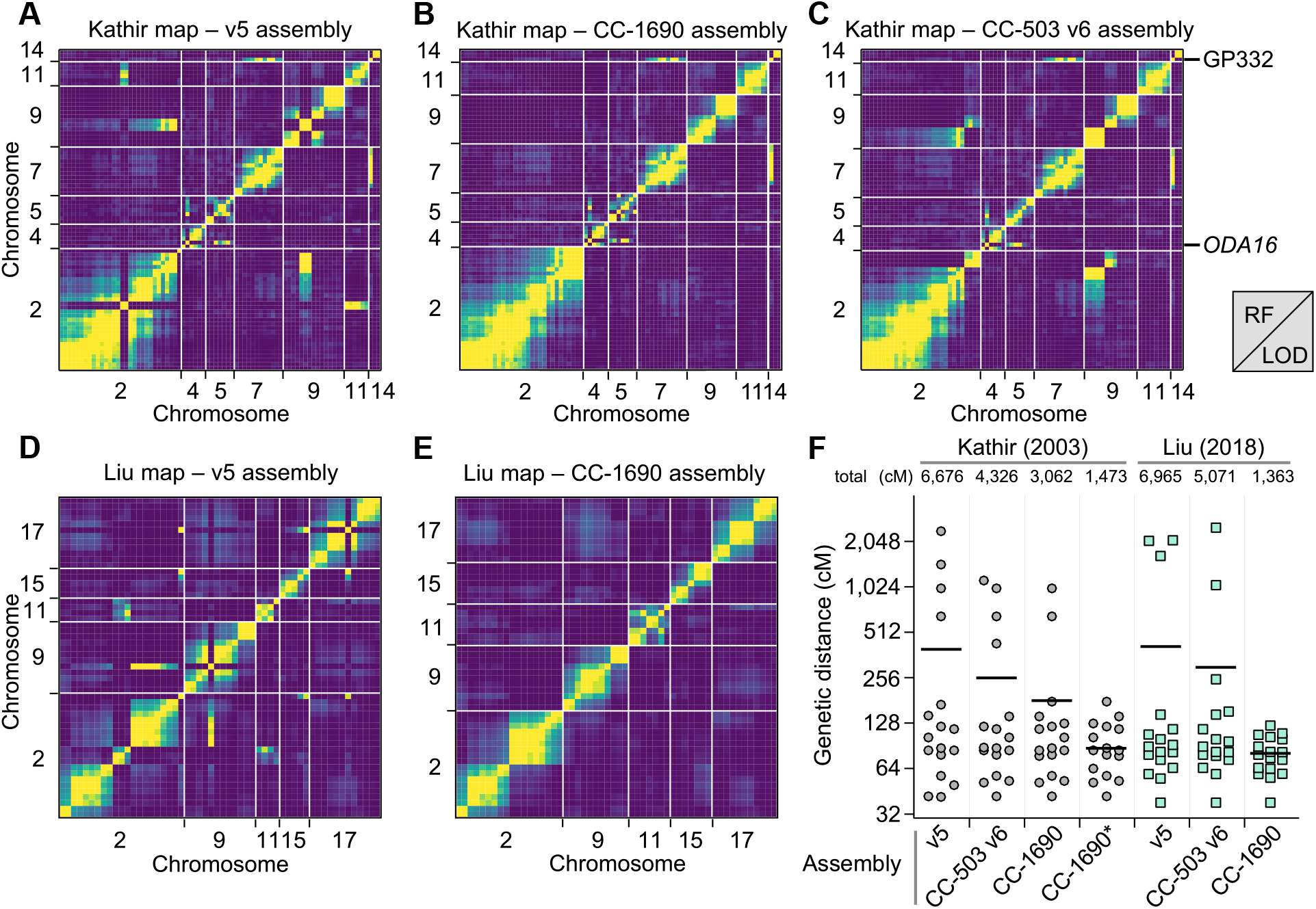
Validation of the Chlamydomonas genome reassemblies by recombination maps. **(A)** Partial plot of recombination frequencies between molecular markers from Kathir et al. (2003). Strong linkage is indicated by a yellow color; absence of linkage is shown as dark blue. **(B, C)** Partial recombination frequency plots between the same molecular markers with updated genomic coordinates according to the CC-1690 (B) or CC-503 v6 (C) assembly. Note that the markers GP332 and *ODA16* are consistently mismapped. (**D, E**) Partial recombination frequency plots between informative SNPs extracted from Liu et al. (2018), when using the genomic coordinates from the v5 **(D)** or CC-1690 **(E)** assemblies. RF: recombination fraction. LOD: logarithm of the odds. **(F)** Gradual improvement of the estimation of genetic map length, from v5, to CC-503 v6, to CC-1690. Chromosome lengths are plotted in cM for each increment of the genetic maps. CC-1690* denotes the use of CC-1690 genomic coordinates with the removal or the GP332 and *ODA16* molecular markers from the analysis. Total map length, in cM, is listed above each dot plot. Horizontal bar, mean.

We followed the same steps to generate a genetic map from whole-genome re-sequencing data of tetrads derived from crosses between two Quebec field isolates (Liu et al. 2018). We reduced the data to keep only single nucleotide polymorphisms (SNPs) that were informative of haplotype transitions (164 SNPs). Again, the deduced recombination map largely agreed with v5 chromosomal positions, except for 14 SNPs, eight of which had been wrongly assigned to chromosome 2 (Figure 3D). The CC-1690 genomic coordinates corrected all mismapping (Figure 3E) and greatly reduced the overall length of the genetic map, from over 6,000 cM using v5 coordinates to ∼1,400 with CC-1690 coordinates (Figure 3F). As with the molecular markers, any discordance between CC-1690 and CC-503 v6 mapped to the putative rearrangement affecting chromosomes 2 and 9. We therefore conclude that CC-1690, and thus the v6 assemblies, receives strong recombination support from two independent mapping datasets, which were derived from both a laboratory strain (CC-1690) and diverse field isolates (CC-1952 in one case, CC-2935 and CC-2936 in the other). These results validate the improvements from v5, and it is now expected that the order and orientation of chromosomal sequence in the CC-1690, CC-4532 v6 and CC-503 v6 assemblies represents the biological reality for these strains.

### The sequence context of assembly gaps

Building from the contiguity improvements described above, we analyzed the filled and remaining assembly gaps relative to the repeat landscape of the Chlamydomonas genome. We annotated almost 1,000 filled v5 gaps based on their sequence context in CC-4532 v6, either as “TE” (∼8% of the gaps), “microsatellite” (16%) or “satellite” (12%) if the novel sequence featured >50% of the corresponding repeat class, “repetitive” (15% gaps) if the sequence had >25% repeat content but not >50% for one of the above classes, and “other” (26%) for less repetitive sequences (Figure 4A). Microsatellites and satellites are tandem repeats that differ in the length of repeated monomers, here either <10 bp (microsatellites) or ≥10 bp (satellites). We further classified gaps based on their overlap with the CC-4532 v6 structural annotations (see below) as either entirely intergenic (∼19% of the gaps), entirely intronic (34%) or at least partially exonic (47%) (i.e. the filled sequence contained at least some novel exonic sequence annotated de novo in CC-4532 v6). Tandem repeats were associated with nearly four times as many gaps as TEs, despite covering almost half as much of the genome (Table 1). Furthermore, while 81% of TE-associated gaps were intergenic, 84% of gaps associated with tandem repeats were within genes (Figure 4A). These results are consistent with the previously reported underrepresentation of TEs (Philippsen et al. 2016) and overrepresentation of tandem repeats (Zhao et al. 2014) in introns, and are consistent with our own annotation of repeats by site class (Supplemental Table S2). The high proportion of genic gaps supports the study of Tulin and Cross (2016), which identified more than 100 “hidden” exons by comparing a de novo-assembled transcriptome to the v5 assembly. Although many gaps containing exonic sequence were not particularly repetitive overall (class “other”), they often featured repeats at their flanks (see Figure 5A example). Overall, our results suggest that prior targeted gap filling was largely successful in assembling intergenic TEs, while the higher density of intronic tandem repeats precluded the more complete assembly of genic regions by Sanger and short-read technologies. Finally, we noticed that 23% of gaps were not filled in v6 but instead lost redundant sequence from one or both flanks (class “redundant”, Figure 4A). Approximately half of these cases resulted in the removal of redundant exonic sequence, providing further potential to improve structural annotation (see Figure 5B example).

The CC-4532 v6 chromosomes still contain 63 gaps that generally coincide with the most repetitive genomic regions. Approximately one third fall within the putative centromeres and subtelomeres, with another third accounted for by tandem repeats, especially large satellites (Figure 4B). Despite the complexity of the *Sultan* satellites present at subtelomeres, 26 of the 34 chromosome termini are capped with telomeric repeats. Among the incomplete termini are the two ribosomal DNA (rDNA) arrays on the right arms of chromosome 8 and 14 (Figure 1A; note that the chromosome 1 rDNA array is truncated and likely non-functional in laboratory strains, but potentially not so in field isolates (Chaux-Jukic et al. 2021)). One gap corresponds to the 5S rDNA array on chromosome 1, while the second 5S rDNA array on chromosome 8 is putatively complete (Figure 1A). Although approximately half of the microsatellite-associated gaps are intronic, almost all the remaining repeat-associated gaps are intergenic. Unfortunately, 12 gaps contain exonic sequence, putatively affecting 18 genes based on comparison to CC-503 v6 (Supplemental Table S3). Most of these gaps are not obviously repetitive (“other” class, Figure 4B) and will be prime targets for future manual finishing.

**Figure 4.**
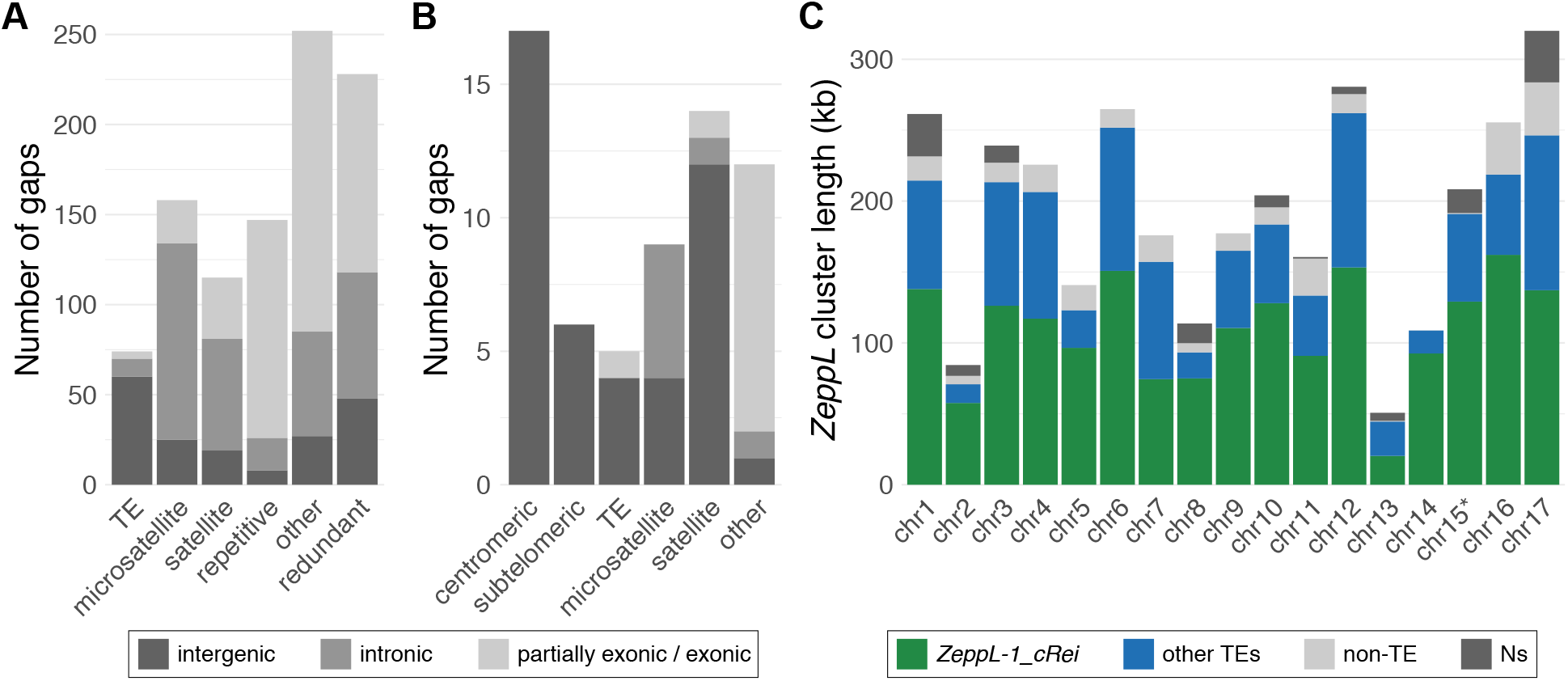
Filled gaps and the remaining assembly challenges in CC-4532 version 6. (**A**) Repeat classification of v5 gaps filled in CC-4532 v6, bars are split into entirely intergenic gaps, entirely intronic gaps and gaps with at least partial exonic overlap. See main text for details of gap definitions by repeat class. **(B)** Classification of the remaining gaps in CC-4532 v6, shading follows **(A)**. “Other” gaps were associated with other repeat types (e.g. large duplications) or were not clearly associated with repeats. **(C)** Summary of the length of putative centromeric *ZeppL* clusters. Colors represent the number of bases annotated as *ZeppL-1_cRei* (the only *ZeppL* family in Chlamydomonas), any other TE, non-TE sequence, and assembly gaps (Ns). Note that chromosome 15 contains two short *ZeppL* clusters downstream of the main cluster (Supplemental Table S4), which are not shown.

Following the misassembly corrections, each v6 chromosome features a single localized cluster of *ZeppL* elements (Figure 1A), except for chromosome 15, where we identified two minor clusters (∼30 kb and 9 kb) downstream of the major cluster. Although most putative centromeres feature at least one gap, they are not particularly long; by comparison to the CC-1690 assembly, we estimate that more than 95% of putatively centromeric sequence is assembled in CC-4532 v6 (Figure 4C, Supplemental Table S4). Estimating their length based on the span of *ZeppL* elements, the putative centromeres range from 51 – 320 kb, with a mean of 192 kb. Approximately 60% of the sequence is composed of the *ZeppL* element itself, with most of the remaining sequence contributed by other TEs (Figure 1A and 4C, see also Supplemental Figure S3 for CC-1690), as reported previously (Craig et al. 2021a). Satellite DNA does not appear to be a major component of the clusters (except chromosome 16, Supplemental Table S4), although we observed satellites immediately flanking the clusters on some chromosomes (e.g. 4 and 5, Supplemental Figure S3) that may form part of the centromere. The exact repeat structure of these regions warrants further study, as does the localization of centromeric histone H3, which may be encoded by two paralogous genes in Chlamydomonas (Cui et al. 2015).

Finally, we revisited the genomic landscape of CG methylation (C^5^-methylcytosine, 5mC) in Chlamydomonas. Lopez et al. (2015) identified 23 hypermethylated loci relative to a genomic background of very low methylation (<1%). We determined that 19 of the hypermethylated regions coincide with the putative centromeres on 11 chromosomes, with a further two localizing to subtelomeres (Figure 1A, Supplemental Figure S3). Chaux-Jukic et al. (2021) called CG methylation directly from Nanopore reads, which facilitates mapping to highly repetitive regions, revealing ubiquitous hypermethylation of subtelomeres. Using the same Nanopore dataset (Liu et al. 2019) and approach, we extended this analysis to the entire CC-1690 assembly and established that all the putative centromeres are hypermethylated (Supplemental Figure S3). Alongside subtelomeres, a few other highly repetitive regions were hypermethylated (e.g. a ∼200 kb region on the left arm of chromosome 12), while we frequently observed more localized methylation peaks of smaller magnitude. Presumably, these regions were previously overlooked due to the limitations of mapping short-read bisulfite sequencing data to repeats and the incompleteness of some of the most repetitive regions in v5. Strenkert et al. (2021) reported an atypical chromatin architecture for the previously identified hypermethylated regions, and it is possible that the hypermethylated centromeres, subtelomeres, and potentially some other repeat-rich islands, constitute heterochromatin in Chlamydomonas.

### Version 6 structural annotations

We annotated both the CC-4532 v6 and CC-503 v6 assemblies de novo, incorporating Iso-Seq data, more than 500 Gb of RNA-seq, and protein homology from the growing number of green algal structural annotations. Notably, more than 1.6 billion strand-specific 150 bp RNA-seq read pairs were introduced from the JGI Gene Atlas (https://phytozome-next.jgi.doe.gov/geneatlas/), which assessed gene expression under 25 conditions. We predicted gene models using several annotation tools, with the model receiving the best support from transcriptomic and protein homology evidence retained in cases of redundancy. Focusing on CC-4532 v6, we then made several further improvements (see below) to the de novo gene models to arrive at the final CC-4532 v6 annotation, named CC-4532 v6.1, featuring 16,801 protein-coding genes (Table 2). The number of predicted alternative transcripts also increased more than eight-fold relative to v5.6. Dedicated analyses will be required to validate these new isoforms (see Labadorf et al. (2010); Raj-Kumar et al. (2017)). One highlight of the annotations was that the longest transcripts overlap for 29% of adjacent genes, 64% of which are on opposite strands (see examples in Figure 5). While the longest transcripts may not always be the most abundant, this result nevertheless speaks to the compactness of the genome. Overlapping models were essentially absent from v5.6 (1% of neighboring genes) and were largely made possible by Iso-Seq support, and even the present count may be an underestimate since these data do not cover all genes.

**Table 2.** Comparison of structural annotations between reference genome versions.

| Annotation | CC-503 v4.3 | CC-503 v5.6 | CC-503 v6.1 | CC-4532 v6.1 |
| --- | --- | --- | --- | --- |
| Nuclear genes | 17,114 | 17,741 | 16,795 | 16,801* |
| Alternative transcripts | / | 1,789 | 14,874 | 14,979 |
| Transposable element genes | / | / | 647 | 810 |
| Low coding potential genes | / | / | 1,435 | 1,417 |
| Plastome genes | / | / | 74** | 74** |
| Mitogenome genes | / | / | 8 | 8 |
| BUSCO (chlorophyta_odb10, N=1,519) | C:96.7%<br>[S:96.0%,D:0.7%]<br>F:1.3%,M:2.0% | C:98.9%<br>[S:98.2%,D:0.7%]<br>F:0.3%,M:0.8% | C:100.0%<br>[S:99.3%,D:0.7%]<br>F:0.1%,M:0.0% | C:99.8%<br>[S:98.8%,D:1.0%]<br>F:0.1%,M:0.1% |

C, complete; S, single-copy; D, duplicated; F, fragmented; M, missing.
\*CC-4532 v6.1 contains 16 *MT+* specific genes (see below).
\*\*the three trans-spliced exons of *psaA* are here counted as individual models.

Since so many of the v5 assembly gaps were within genes, the assembly improvements provided considerable potential to improve gene models. First highlighted by Tulin and Cross (2016) as an example of a gene featuring “hidden exons”, *PARALYZED FLAGELLA 20* (*PF20*) encodes a 606 amino acid (aa) protein important for cilia function (Smith and Lefebvre 1997). The filling of a v5 assembly gap in *PF20* resulted in the correction of the gene model in CC-4532 v6.1, adding three new exons (exons 9, 10 and 11 in CC-4532 v6.1) and shifting the 3’ splice site of exon 8 (Figure 5A). A second example is the putative metal ion transporter *NATURAL RESISTANCE-ASSOCIATED MACROPHAGE PROTEIN 2* (*NRAMP2*), which featured two gaps in v5 that were both classified as “redundant” in our prior analysis. While one “gap” duplicated only 26 bp of intronic sequence, the second duplicated exons 10 and 11, resulting in the erroneous repetition of 63 aa in the v5 protein (Figure 5B). Although this exonic duplication fortuitously maintained the reading frame, this was not always the case for other affected genes. Finally, while *PF20* and *NRAMP2* were annotated as single genes in v5, some genes were incorrectly split into separate models by assembly gaps (Supplemental Figure S4). We chose these examples from hundreds of affected genes, demonstrating the scale of improvement made possible by assembly improvements.

**Figure 5.**
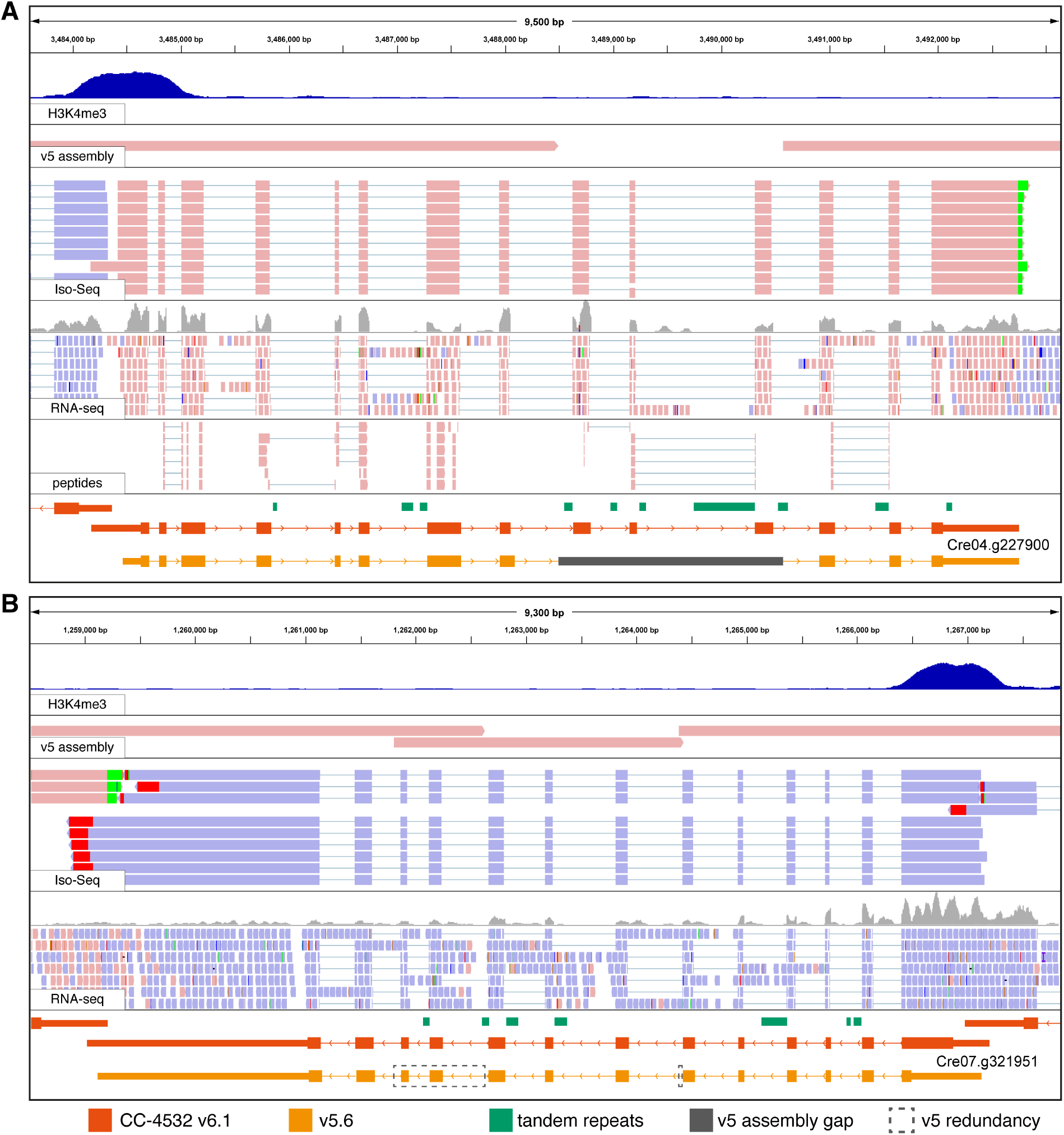
Browser views of example gene models improved between v5.6 and CC-4532 v6.1. H3K4me3 ChIP-seq (dark blue peaks) marks promoters. The v5 assembly track shows an alignment of v5 contigs to CC-4532 v6, with assembly gaps appearing as unmapped regions and redundant sequence as overlapping regions. Peptides are from mass spectrometry analysis of the proteome. Coordinates for v5.6 gene models (orange) were converted to CC-4532 v6. Thick blocks represent coding sequence, thin blocks UTRs, and conjoining lines introns. Forward strand mappings are shown in pink and reverse in blue. Red and green mismatches at the end of Iso-Seq reads correspond to poly(A) tails. **(A)** *PF20*, CC-4532 v6 coordinates: chromosome 4, 3,483,590 – 3,493,250. **(B)** *NRAMP2*, CC-4532 v6 coordinates: chromosome 7, 1,258,513 – 1,267,855. Note that the redundant sequences (boxed) are not included in the gene model converted from v5.6, since these duplicated sequences do not exist in CC-4532 v6. No peptides were identified.

We further focused on several specific issues that have been previously highlighted. Cross (2015) showed that more than 4,000 v5 gene models have in-frame upstream ORFs, many of which likely correspond to genuine N-terminal protein extensions based on comparison to *V. carteri* orthologs. To address this issue, we generally annotated the first in-frame start codon for each predicted mRNA as the start codon in the v6 annotations. *NRAMP2* also serves as an example of this change, with the CC-4532 v6.1 protein extended by 126 aa at its N terminus (Figure 5B). Second, two studies (Blaby and Blaby-Haas 2017; Craig et al. 2021a) reported more than 100 strongly supported gene models that are absent from the v5 annotations. Many of these genes were present in the v4 annotations (e.g. *PSBW1*), and 25 are part of polycistronic transcripts (Gallaher et al. 2021). We attempted to transfer any strongly supported gene model from the v4.3, v5.6 or preliminary CC-503 v6 annotations to CC-4532 v6.1 if they were absent in the preliminary de novo annotation. Third, we manually curated a modest number of genes of interest, including 12 encoding selenoproteins (Novoselov et al. 2002) that were all previously misannotated due to their use of the canonical stop codon “TGA” to encode selenocysteine.

Finally, two changes based on our previous analyses (Craig et al. 2021a) caused the nuclear gene count to fall by 940 between v5.6 and CC-4532 v6.1. First, more than 1,000 v5.6 genes are likely part of TEs, and these gene models have been moved to their own category in the v6 annotations (“transposable element genes”, Table 2). The motivations for this category are detailed below. Second, several hundred v5.6 genes were reported to have low coding potential and are unlikely to represent protein-coding genes. This designation was reached by combining evidence from functional annotation, comparative genomics, population genetics, and intrinsic features of Chlamydomonas genes and coding sequence (codon usage bias and the strength of Kozak-like sequences). We repeated these analyses on the de novo v6 annotations, conservatively calling 1,417 “low coding potential” gene models in CC-4532 v6.1 (Table 2, Supplemental Figures S5 and S6). Validating these analyses, we found no peptide support for these models in our proteomics analysis (see below). We did not include these models in the main annotations, but they can be accessed from Phytozome by interested users. Many of these loci with transcriptomic support may be long noncoding RNA (lncRNA) genes that contain spurious short ORFs, or simply short ORFs located within the untranslated regions (UTRs) of other genes.

### Organelle genomes and structural annotations

The genomes of the plastid and mitochondria, the plastome and mitogenome, respectively, have not been included in any previous reference versions. They encode abundant cellular proteins (see below) and contribute disproportionately to the transcriptome: 46% of the cell’s mRNA is transcribed from the plastome, and just eight mitochondrial genes contribute 1.4% to the total mRNA pool (Gallaher et al. 2018). We recently produced high-quality assemblies and annotations of the plastome and mitogenome (Gallaher et al. 2018), which are now included in the v6 releases (Table 2). Importantly, there are no genetic variants to distinguish the organelle genomes of CC-4532 and CC-503, since the laboratory strains are putatively descended from the same zygote and the multicopy organelle genomes are inherited uniparentally.

The circular plastome is 205.6 kb, features two inverted repeat regions and two single copy regions, and is present in 83 copies on average per cell (Gallaher et al. 2018). It carries 72 protein-coding genes, with two (*psbA* and *I-CreI*) duplicated in the two repeat regions. Many of the genes are expressed from polycistronic transcripts. Cavaiuolo et al. (2017) used small RNA profiling to accurately map the plastid genes, and we incorporated their improvements to the v6.1 annotations. The *psaA* gene, which encodes photosystem I chlorophyll *a* binding apoprotein A1, is expressed as three separate transcripts that are trans-spliced to generate the mature mRNA molecule (Kück et al. 1987). The three separate genes that contribute to the mature transcript are out of order and in different orientations, a situation that many computational tools are not equipped to handle. As such, we assigned three separate, but sequential, gene IDs (CreCp.g802280, CreCp.g802281, and CreCp.g802282) to the three *psaA* exons.

The 15.8 kb mitogenome is a linear molecule, as found in several other closely related species (Hamaji et al. 2017). Each cell carries 183 copies of the mitogenome, on average, distributed among its mitochondria. It carries only eight protein-coding genes, which are expressed from a single bidirectional promoter. Seven of these genes encode components of the respiratory complex, while the eighth, reverse transcriptase-like (*rtl*), is likely required for mitogenome replication (Smith and Craig 2021). We incorporated the more accurate annotations of Salinas-Giegé et al. (2017), who demonstrated that the 5’ end of each mature mitochondrial transcript begins immediately at the start codon (i.e. there are no 5’ UTRs).

### Gene model validation

We successfully mapped existing “Cre” IDs from v5.6 (see below) to 15,224 nuclear genes in the CC-4532 v6.1 annotation (90.6%, Supplemental Dataset S1). The remaining gene models were either novel or had changed considerably relative to their v5.6 counterparts (due to gene model mergers, splits, etc.). We validated the CC-4532 v6.1 annotation using several independent approaches. We first queried all predicted proteins against the BUSCO (Benchmarking Universal Single Copy Orthologs) chlorophyte dataset (Manni et al. 2021), with the number of fragmented and missing genes dropping from five and eleven, respectively, in v5.6, to only one and two in CC-4532 v6.1 (Table 2). Notably, CC-503 v6.1 had no missing genes, and upon inspection, the two missing genes in CC-4532 v6 were found within the small number of remaining genic gaps (see above). Nevertheless, we consider the CC-4532 v6.1 annotation to be superior to that of CC-503: as detailed below, many more genes are affected by major loss-of-function mutations in CC-503, although none are genes in the BUSCO dataset (many of which may be essential).

We next turned to chromatin-immunoprecipitation followed by deep-sequencing (ChIP-seq) data of trimethylated histone H3 lysine 4 (i.e. H3K4me3), which reliably mark promoter regions (Ngan et al. 2015; Strenkert et al. 2021) (see Figure 5). We queried 1,224 H3K4me3 peaks that had been called as intergenic relative to the v5 genome and v5.6 annotation, assigning 244 peaks to gene transcription start sites (TSSs) in CC-4532 v6.1. Approximately 30% of the genes newly associated with H3K4me3 peaks did not have gene IDs mapped forward from v5.6, suggesting that the improvements can be attributed to both the inclusion of new genes and changes to the TSSs of existing genes. It is not surprising that almost 1,000 H3K4me3 peaks remain unannotated, since they are expected to be associated with features other than protein-coding genes, such as lncRNAs (Strenkert et al. 2021). Furthermore, ∼40% of the remaining peaks coincided with TEs, which may be an underappreciated source of active promoters in Chlamydomonas.

Finally, we queried the v5.6 and CC-4532 v6.1 predicted proteins against a pool of proteomics data. We identified at least one unambiguously assigned peptide for 14,339 v5.6 proteins and 14,841 v6.1 proteins, an increase of 3.5% (Supplemental Table S5). The v6.1 total included 14,770 proteins encoded by the nuclear genome (including TE proteins), 65 from the plastome, and 6 from the mitogenome. We noted a 7.2% increase in the total number of unique peptides assigned to CC-4532 v6.1 relative to v5.6, and a 7.0% increase in the total number of peptides. These increases can be attributed to several improvements in v6.1, including the incorporation of entirely new nuclear genes, the inclusion of new exons within previous assembly gaps, and the N-terminal ORF extensions. For example, we identified three unique peptides assigned to the previously “hidden” exons of *PF20* (Figure 5A). The addition of the organelle annotations also contributed substantially. This was especially true for the total number of peptides, since the 65 plastome-encoded proteins with identified peptides accounted for 5.0% of all peptides assigned to CC-4532 v6.1, and the 6 mitogenome-encoded proteins accounted for 0.018%. Notably, these estimates are far lower than the total mRNA contribution from the organelles to the cell (see above).

### Gene IDs

Starting with the v4 annotations and becoming standard for all genes in the v5.5 release, Chlamydomonas locus IDs have taken the form CreYY.gNNNNNN, where YY is the chromosome number and NNNNNN is a unique number that nominally increases along the chromosome (Blaby et al. 2014). These IDs are generally reported in the literature together with gene symbols when available, and maintaining this format is critical for continuity. We aggressively mapped the v5.6 IDs forward onto the CC-4532 v6.1 models, and manually curated the transfer of IDs for v5.6 genes with expert annotation (gene symbols or descriptions) where automation failed. For the 2,277 CC-4532 v6.1 genes with no v5 equivalent (including TE genes and organelle genes), new NNNNNN numbers were introduced, ranging from 800000 to 802251 and increasing with genomic coordinates. All plastome and mitogenome genes were assigned new locus identifiers, from CreCp.g802263 to CreCp.g802335, and from CreMt.g802337 to CreMt.g802344. Since we also annotated the CC-503 v6 assembly (and many more genomes may follow), it was necessary to distinguish between orthologous gene models annotated in each assembly. To achieve this, we included a four-digit strain-specific suffix to the IDs: CreYY.gNNNNNN_4532 for CC-4532 v6.1 and CreYY.gNNNNNN_0503 for CC-503 v6.1. With CC-4532 becoming the reference, gene models in other assemblies (including CC-503 v6) will be attributed IDs based on their mapping to this annotation.

It is also imperative to note that the misassembly corrections reported above and the CC-503 structural rearrangements (see below) resulted in many genes having mapped forward IDs in CC-4532 v6 that now refer to the wrong chromosome (i.e. YY number). Similarly, the NNNNNN numbers may no longer be contiguous with their new neighbors. In fact, this was already an issue for some IDs in v5 due to assembly changes relative to v4. Unfortunately, both the YY and NNNNNN numbers are now meaningless (and in fact potentially misleading) for many genes, and users are cautioned that no spatial information should be extracted from the IDs. To counter any confusion that could arise from this issue, we devised a new spatially correct “associated locus ID” for each gene. They follow the format XXXX_YY_NNNNN, where XXXX is the strain identifier from the Chlamydomonas Resource Center, YY is the chromosome number, and NNNNN is a unique gene number that increases along the chromosome, with odd numbers for forward strand genes and even numbers for reverse strand genes. Successive IDs feature NNNNN numbers separated by 3 or 4 unused numbers depending on relative strandedness (rising to 53 or 54 for genes on either side of an assembly gap), serving as placeholders for possible new gene models. As an example, in CC-4532 v6.1 *PSY1* has the primary ID Cre02.g095092_4532 and the associated locus ID 4532_11_52343, with the latter providing the correct chromosomal location (Figure 2B). These IDs also carry additional information as optional suffixes e.g. all transposable element genes feature the suffix “_TE”, making them instantly recognizable. The associated locus IDs have a one-to-one relationship with the existing “Cre” IDs (Supplemental Dataset S1) and we envision that they will be used in parallel (e.g. to simultaneously assess spatial information).

### Expert annotation and gene symbols

Over decades of research, Chlamydomonas genes have been assigned a gene symbol, designed to uniquely identify and succinctly characterize a given locus. In v5.6, 5,130 genes (28.9%) were annotated with a gene symbol (Supplemental Table S6). Some of these symbols were assigned to connect a genetic locus to the protein it encodes e.g. the gene that encodes the plastocyanin protein, Cre03.g182551, was named *PCY1*. Others were based on experimental characterization e.g. Cre09.g390023 was named *CRR1* due to its role as a <u>C</u>u responsive <u>r</u>egulator of gene expression (Sommer et al. 2010). Many more were assigned based on a phenotype identified for mutants at that locus. For example, the *NIT1* locus is so named because the *nit1* mutations in this gene prevent utilization of nitrate as a N source. Some genes were assigned symbols in lists resulting from large scale gene/protein identification experiments or phylogenomic comparisons, as in the “GreenCut” for genes conserved in the green lineage, or the “CiliaCut” for genes conserved in ciliated organisms (Merchant et al. 2007), exemplified by *CGL75* for Cre07.g337516 and *MOT7* for Cre01.g038750. Many gene symbols were assigned based on orthology to genes named in other model organisms; especially streptophytes like Arabidopsis for plant-specific genes (Cre05.g246800 is *GUN4* for orthology to a gene with that name in Arabidopsis), *Saccharomyces cerevisiae* for common eukaryotic genes (Cre02.g073650 is *PRP19* based on its orthology to the yeast *PRP19* gene), and (cyano)bacteria (Cre07.g329300 is *MSC1* based on its relationship to *mscS* in *Escherichia coli*). Lastly, there have been systematic attempts to name groups of functionally related proteins, such as the chloroplast protein translocation complexes, TIC and TOC (Kalanon and McFadden 2008).

These gene symbols are an important and powerful tool for interpreting, analyzing, and communicating research in Chlamydomonas, especially for large-scale and systems biology research. There is constant pressure to assign more gene symbols to the 71% of genes that currently lack them, and this has driven the proliferation of uninformative, and even erroneous, gene symbols. For example, the root “*ANK*” was used to assign gene symbols to 20 genes in v5.6 due solely to the presence of a predicted ankyrin repeat domain. Similarly, there are 51 *HEL* genes (encoding proteins with a DEAD/DEAH box helicase domain) and 35 *DNJ* genes (encoding proteins with a DnaJ domain) in v5.6. These symbols do not provide more information than their automated PFAM domain annotations. The presence of a gene symbol may imply that the gene has been at least partially characterized and perhaps has a validated function corresponding to the name, but none of this applies for the 51 *HEL* genes or the 34 *DNJ* genes beyond the presence of a certain domain. Furthermore, some symbols rely on erroneous predictions. For example, Cre14.g629650 was named *NIK1* because a simple homology search suggested that this gene encodes a Ni transporter. However, Chlamydomonas has no known Ni requiring genes and no nutritional requirement for Ni, so this functional annotation is probably incorrect (Blaby-Haas et al. 2016).

Ambiguous, uninformative, and erroneous gene symbols can complicate and confuse the analysis of systems biology data and can even inhibit efforts to identify the correct function of a gene. The Chlamydomonas annotations are frequently used to help guide the annotation of newly sequenced Chlorophyte genomes (Roth et al. 2017), which propagates the low information or misinformation throughout the Chlorophyte lineage. Therefore, we sought to improve and update the gene symbols, which consisted of three phases: 1) the addition of new gene symbols wherever those annotations were based on expert analysis or empirical data, 2) transfer of a primary gene symbol to “previous identifiers” for uninformative and misleading gene symbols, and 3) reformatting or changing existing gene symbols to conform to a uniform style.

We added 610 new gene symbols to the CC-4532 v6.1 annotation. The majority of these were assigned in collaboration with the authors of individual chapters in the forthcoming 3rd edition of the Chlamydomonas Sourcebook. Still others were based on recent publications. We reclassified 1,332 v5.6 gene symbols as “previous identifiers”, preserving connections to historical research that may have used those symbols (Supplemental Dataset S2). Resultingly, there are now 4,408 out of 16,801 (26.2%) genes with a gene symbol in v6.1 (excluding TE genes, none of which presently have symbols). An additional 549 genes had their gene symbol replaced, altered, or reformatted to improve clarity, highlight orthologies and unify formatting. This effort was guided by several rules, updated and expanded from our previous work (Blaby et al. 2014). These rules can be found as an appendix to this text, and we recommend that they be applied for the naming of all Chlamydomonas genes going forward.

While important, the gene symbols are only one part of communicating functional information about a given locus. Many genes, including those that have not yet been assigned a symbol, have a defline and associated comments. These may include a description of the gene’s function, relevant expression data, paralogy and orthology information, and links to related peer-reviewed literature. This last feature, in the form of PMID accession numbers, has also been expanded and updated from 1,852 genes supported by one or more PMIDs (2,626 total PMIDs) in v5.6, to 3,042 genes (4,697 total PMIDs) in CC-4532 v6.1 (Supplemental Table S6).

Finally, the rate at which genes are expertly annotated in the literature outpaces that of updates to the Chlamydomonas genome and structural annotations. We have therefore created a dedicated email account, to receive and store user updates. We encourage users to send curated annotation updates. This may include gene symbol suggestions, textual annotation, PMIDs, expression data, functional validation, among other information. We also welcome manually curated gene models (preferably in GFF3 format), either for entirely new genes or for evidence-based corrections to existing models. We are committed to collating this information so that future updates are both efficient and representative of recent advances in Chlamydomonas research.

### Haplotype 2 and the mating type locus

The genomes of Chlamydomonas laboratory strains are collectively comprised of two divergent parental haplotypes (Gallaher et al. 2015). Haplotype 1 was arbitrarily defined relative to CC-503, with any region with the other haplotype in any other strain referred to as haplotype 2. CC-4532 contains five haplotype 2 regions spanning 4.6% of the genome (Figure 1A, Supplemental Figure S7) and featuring 818 genes (Supplemental Table S7). Many variants of functional significance are likely to segregate between the haplotypes, including SNPs, small insertions and deletions (indels) and larger structural variants (Gallaher et al. 2015). The latter category may also include presence/absence and copy number variants, which segregate among Chlamydomonas field isolates (Flowers et al. 2015). The rapidly evolving *NCL* gene family, forming a major cluster of 32 genes on chromosome 15 in CC-503 (i.e. haplotype 1) (Boulouis et al. 2015), is one clear example of copy number variation among laboratory strains. The major *NCL* cluster partly intersects a haplotype 2 region in CC-4532 (Figure 1A), and we annotated 43 *NCL* genes as part of the cluster in CC-4532 v6.1. Although this example demonstrates that large variants exist among laboratory strains, we did not perform a systematic analysis of structural variation present between the two haplotypes; the CC-503 – CC-4532 comparison captures less than a fifth of the total haplotype variation among laboratory strains (which can affect up to ∼25% of the genome), and this question would be best addressed by assembling and comparing genomes of additional strains. We did however revise the coordinates of the haplotype 2 blocks reported by Gallaher et al. (2015) relative to CC-4532 v6 (Supplemental Table S8), since some were affected by assembly corrections. The distribution of haplotype blocks among many of the most widely used laboratory strains is shown in Supplemental Figure 8.

The mating type locus (*MT*) on the left arm of chromosome 6 is naturally within a region where strains carry one or the other haplotype, *mt+* strains haplotype 1, and *mt–* strains haplotype 2. Except for genes unique to either allele, *MT* genes have gametologs present on both alleles, although those within the rearranged (R) domain are generally not syntenic between *MT+* and *MT–*. Since CC-503 is *mt+*, past assembly versions have lacked the two *MT–* specific genes, *MINUS DOMINANCE 1* (*MID1*) and *MATING TYPE REGION D-1* (*MTD1*). With the reference now based on the *mt–* CC-4532 the situation is reversed, however this is a greater issue since there are at least 16 *MT+* specific genes in five *MT+* specific regions, three of which originated from autosomal insertion (*MTP0428*, the MTA region and the SRL region) (De Hoff et al. 2013). To address this, we appended a 375 kb *MT+* R domain contig extracted from CC-503 v6 to the reference CC-4532 v6 assembly. To avoid potential mismapping of ‘omics data we hardmasked (i.e. replaced with Ns) any gametologous regions on the appended contig, so that only sequences corresponding to *MT+* specific regions and genes were included. Finally, we manually curated all R domain gene models and appended *MT+* specific genes to the CC-4532 v6.1 annotation. CC-4532 v6 should thus be suitable for analyses of data from both *mt+* and *mt–* strains, and we expect that the availability of highly contiguous and well-annotated assemblies of both alleles will be a major resource for the Chlamydomonas community.

We compared our resources for CC-503 v6 and CC-4532 v6 to the existing curated *MT+* (CC-503 v4) and *MT–* (CC-2290) annotations of De Hoff et al. (2013) (Figure 6). The gapless CC-4532 R domain (∼211 kb) was entirely syntenic with that of CC-2290 (∼218 kb), although intergenic regions were often unalignable due to variation in the repetitive sequences present. The only major change in both *MT–* and *MT+* affected *OTUBAIN PROTEIN 2* (*OTU2*), which was extended to incorporate the genes *155027* and *MT0618* into a single gene model (i.e. the correct *OTU2* was split across three gene models in CC-2290 and CC-503 v4). The *MT+* allele of *OTU2* was recently shown to function in the uniparental inheritance of the plastome (Joo et al. 2022). In *MT+, OTU2* is located immediately upstream of an *MT+* specific region termed the “16 kb repeats”, consisting of a 17.2 kb tandemly repeated region containing multiple copies of *EARLY ZYGOTE 2* (*EZY2*), *INTEGRASE 1* (*INT1*) and what was previously annotated as *OTU2* (i.e. the repeats contain duplicates of only a 3’ fragment of the full *OTU2* gene, which may be pseudogenized). *INT1* shares strong sequence similarity to the proteins of DIRS retrotransposons from Chlamydomonas (e.g. *TOC3* (Goodwin and Poulter 2004)) and is likely derived from a TE family that is no longer present elsewhere in the genome. Although the reverse transcriptase domain is missing, *INT1* does contain sequence encoding the RNAse H and methyltransferase domains of a DIRS element in addition to the “integrase” (actually a tyrosine recombinase). Assuming *INT1* has not been co-opted, the multiple copies of *EZY2*, which produce zygote-specific transcripts (Ferris et al. 2002), may be the only functional genes in the repeat. The *MT+* specific regions are collectively responsible for the larger size of the *MT+* allele. However, the assembly of the 16 kb repeats remains incomplete in CC-503 v6, with two gaps relative to CC-1690 (which is also *mt+*). We detected no structural variants indicative of mutations between CC-503 v6 and CC-1690 in the R domain, suggesting that CC-503 v6 provides a typical representation of all *mt+* laboratory strains across this region. Notably, there were two full-length copies of *OTU2* annotated in v5 (Joo et al. 2022), however we found no evidence for this state in either CC-503 v6 or CC-1690, and this was likely a misassembly of the regions flanking the 16 kb repeats.

**Figure 6.**
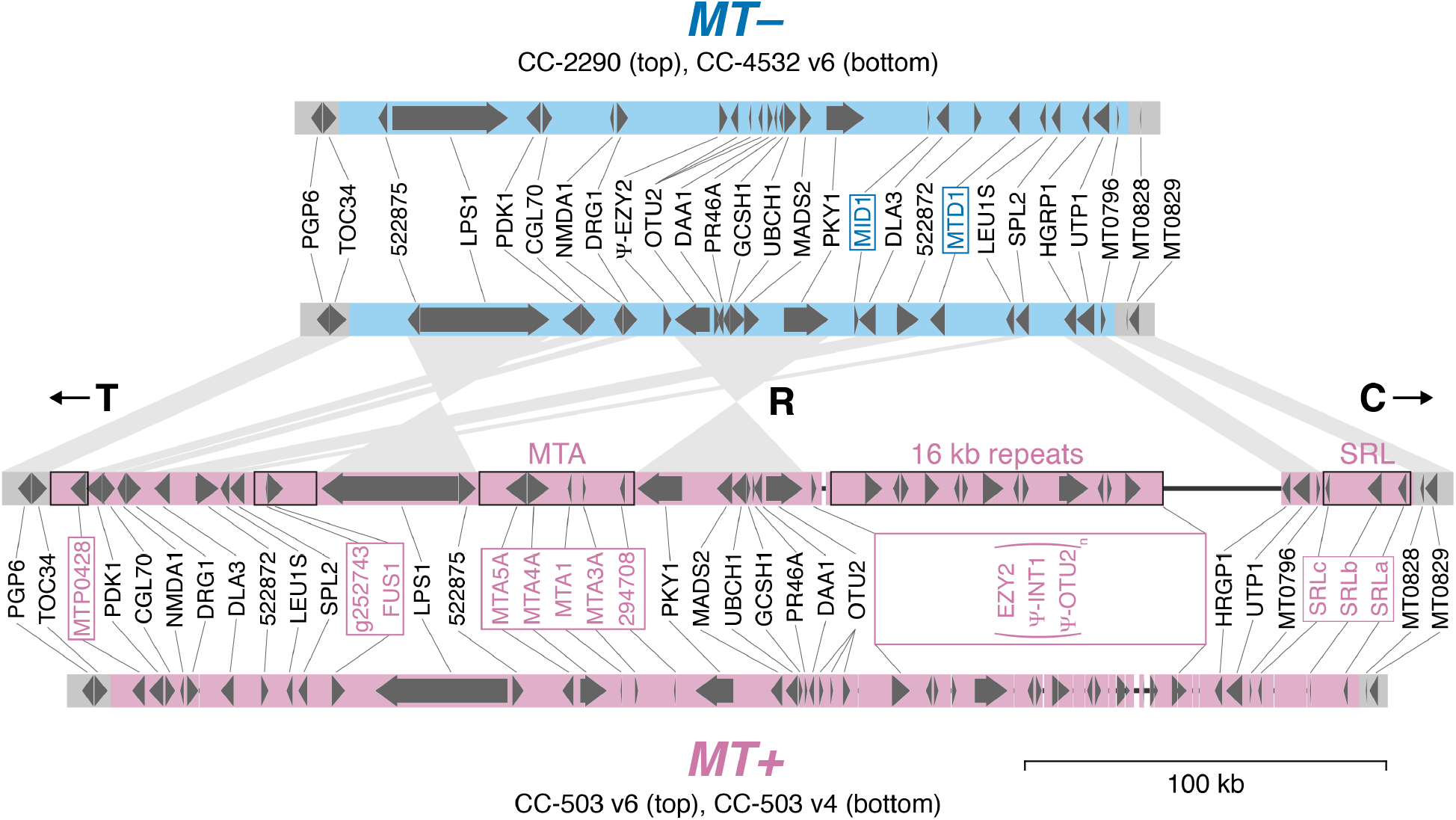
Assembly and annotation comparisons of the *plus* (*MT+*) and *minus* (*MT–*) alleles of the mating type locus rearranged (R) domain. Block arrows represent protein-coding genes. Mating type-specific gene symbols are boxed. CC-503 v6 *MT+* specific regions that were not hardmasked in the *MT+* contig appended to CC-4532 v6 are outlined in black. Synteny between CC-4532 v6 *MT–* and CC-503 v6 *MT+* genes is represented by wedges. T and C refer to the telomere-proximal and centromere-proximal domains, respectively. Only copies of *EZY2* within the 16 kb repeats are included in the *MT+* gene annotation. Copies of *OTU2* within the 16 kb repeats are truncated and are marked as putative pseudogenes (as are *INT1* copies, see main text). Thin black lines represent assembly gaps. The CC-2290 and CC-503 v4 annotations are from De Hoff et al. (2013). Gene symbols are from De Hoff et al. (2013), except for symbols updated herein (Supplemental Table S9).

### The CC-503 genome is unstable and harbors major structural mutations

As outlined above, most differences between the two laboratory strain haplotypes represent genetic variation between two parental genomes. Conversely, any variants segregating between copies of the same haplotype in different strains may be attributed to mutations arising in the laboratory, since all copies of the same haplotype are expected to have been inherited from the same ancestor. We thus performed a systematic analysis of within-haplotype structural variation by comparing haplotype 1 regions in CC-4532 v6 and CC-503 v6, polarizing mutations relative to CC-1690 (i.e. a mutation was called in a given assembly if the other assembly was consistent with CC-1690).

The most conspicuous mutation, large enough to be observed from the linkage analysis (Figure 3), affected chromosomes 2 and 9 in CC-503 v6. Indeed, these chromosomes were misassembled in all past versions, and changes that occurred between v4 and v5 were noted previously (Lin et al. 2013). In v5, the aberration was misassembled as a complex translocation that would have involved at least five DSBs (Figure 7A), presumably due to conflicting evidence from read-based contig assembly and linkage-based scaffolding. Via manual inspection of the CC-503 v6 contigs, we inferred that chromosomes 2 and 9 have instead experienced a putative reciprocal translocation, with an inversion affecting part of the fragment translocated from chromosome 2 to 9 (Figure 7B). This model implies three DSBs, one on chromosome 9 (DSB2 between purple and vermilion, Figure 7B) and two on chromosome 2 (DSB1 between blue and green, and DSB3 green and orange). The 0.9 Mb inversion appears to share DSB1 with the translocation event, implying that all three DSBs occurred, and were subsequently misrepaired, simultaneously. Notably, all DSBs and their repair events were associated with indels, ranging from a few bp to 1,950 bp, and all were predicted to disrupt coding sequence relative to CC-4532 v6.1 (Supplemental Figure S9). As an example, the deletion at DSB2 entirely removed the second exon of a gene (Cre09.g390100) encoding a 318 aa protein with an *S*-adenosylmethioine-dependent methyltransferase domain, with the remaining (and presumably pseudogenized) exons now split between the derived chromosomes 2 and 9 in CC-503 v6 (Supplemental Figure S10). To reiterate, we are confident that CC-503 represents the mutant state, since CC-4532 v6 and CC-1690 are entirely consistent both with each other and the linkage evidence, which was partly derived from field isolates. Furthermore, re-sequencing data from CC-125 (the progenitor of CC-503) mapped across the deletions at each DSB (Supplemental Figure S11), implying that the mutation is unique to CC-503.

**Figure 7.**
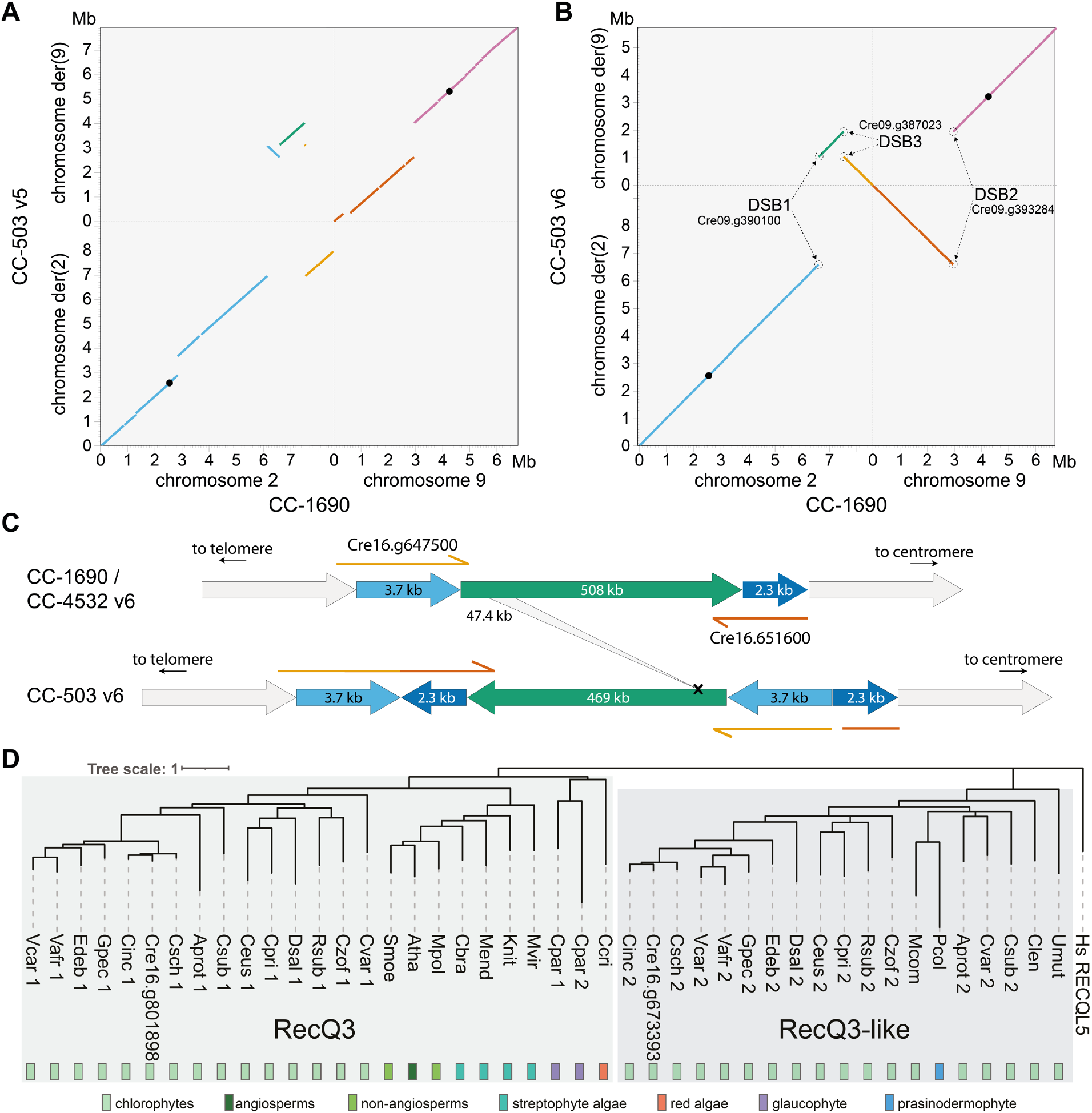
Structural mutations in the CC-503 version 6 genome. (**A, B**) Dotplot representation of chromosomes 2 and 9 between v5 and CC-1690 **(A)**, and CC-503 v6 and CC-1690 (**B**). Colors link fragments between panels **(A)** and **(B)**. Black circles represent putative centromeres. CC-503 chromosomes are named as derivatives (der) based on their centromeres. Genes disrupted by DSBs are labelled. **(C)** Schematic of the dupINVdup and deletion double mutation. The duplicated flanks (light and dark blue) are shown 50x the scale of the main inverted fragment (green). Disrupted and partly duplicated genes are labelled. The left flank is predicted to have formed a gene fusion in CC-503 v6.1, although this is entirely based on ab initio prediction. The 47.4 kb internal deletion is represented by the gray ribbon. **(D)** Protein-based phylogeny of the RecQ3 and RecQ3-like subfamilies of RecQ helicases in Archaeplastida. Branches with bootstrap values <50% were removed. Full species names and protein IDs can be found in Supplemental Dataset 3.

Remarkably, we identified 71 additional structural mutations present in CC-503 v6 and absent in CC-4532 v6 and CC-1690, putatively affecting 103 genes (Supplemental Table S10). This number excludes TEs, which are presented separately below. In full, we called 63 deletions (cumulatively 302.1 kb and including events >10 kb), six duplications, one insertion and one inversion. Many of the mutations were complex, for example the duplications were often associated with indels and inversions. One of the most striking mutations was a ∼508 kb inversion between 0.81 and 1.32 Mb on chromosome 16 (Figure 7C). Inspection of the two DSBs and their subsequent repair revealed that this event is an unusual dupINVdup (duplication-inversion-duplication) mutation (Brand et al. 2015), in which both flanks (3.7 kb to the left and 2.3 kb to the right) of the unique inverted sequence are duplicated and themselves inverted. At both flanks genic sequence was disrupted and partially duplicated (Figure 7C). Surprisingly, the inverted region itself harbored a 47 kb deletion that partially or fully deleted 10 genes (Supplemental Table S10).

Although it is tempting to directly attribute the exceptional number of structural mutations in CC-503 to its past mutagenesis with MNNG (Hyams and Davies 1972), we unexpectedly observed that 46 of the 72 structural mutations were not present in past assembly versions (Supplemental Table S10), including the chromosome 16 dupINVdup/deletion described above. Previous assemblies were primarily based on Sanger sequencing from the initial genome project, while the v6 PacBio sequencing was performed on DNA extracted from a CC-503 culture obtained from the Chlamydomonas Resource Center by Gallaher et al. (2015). Given that many of the mutations were shared between past versions and CC-503 v6, some of which were very large and distinctive (e.g. the reciprocal translocation described above), the more recently acquired culture undoubtedly shares a clonal common ancestor with that used in the original genome project. It therefore appears that the CC-503 genome is unstable, and that approximately two thirds of the mutations may have occurred over the past two decades.

Given the complexity of many of the mutations (e.g. large indels at repair points), we hypothesized that the genomic instability of CC-503 may be the result of a deficiency in DSB repair, potentially induced by mutagenesis and contemporary to the loss of the cell wall. We examined each gene affected by a mutation that was common to CC-503 v6 and all past assembly versions, under the assumption that these mutations could have originated during mutagenesis. We identified a RecQ helicase gene (Cre16.g801898) as a possible candidate, which was fully deleted in CC-503 as part of a 48 kb deletion on chromosome 16 that partially or fully deleted at least six genes (note that this is unrelated to the chromosome 16 deletion described above, see Supplemental Table S10).

RecQ helicases have been referred to as “guardians of the genome” and play key roles in genome maintenance and all DSB repair pathways in human (Croteau et al. 2014; Lu and Davis 2021). Many eukaryotes possess multiple RecQ helicase genes that belong to ancient gene families, with five genes in human and seven in Arabidopsis (Dorn and Puchta 2019). We performed a phylogenetic analysis including the deleted Cre16.g801898 and homologous proteins in algae and plants, which demonstrated that Cre16.g801898 is a putative ortholog of the plant RecQ3 subfamily (Figure 7D), which is homologous to human *RECQ-LIKE HELICASE 5* (*RECQL5*) (Wiedemann et al. 2018). Furthermore, this demonstrated that the RecQ3 subfamily is present across Archaeplastida (the green lineage plus red algae and glaucophytes). Interestingly, our analysis also revealed a green algal-specific subfamily, RecQ3-like, which formed a clade with the canonical RecQ3 subfamily (Figure 7D). All analyzed species from the Chlorophyceae and Trebouxiophyceae had both RecQ3 and RecQ3-like subfamily genes, indicating strong conservation. However, the RecQ3 subfamily appeared to be absent in prasinophytes (e.g. *Micromonas* spp.) and ulvophytes (*Caulerpa lentillifera* and *Ulva mutabilis*). Such a deep evolutionary division between the RecQ3 and RecQ3-like subfamilies is roughly analogous to the plant-specific RecQsim subfamily, which forms a clade with the eukaryotic RecQ6/WRN group (Wiedemann et al. 2018).

The specific functions of RecQ helicases have not been studied in green algae and it is difficult to draw parallels with other species, since the evolution of RecQ helicases is dynamic in many lineages. Certain plants have lost specific subfamilies and duplicated others e.g. the moss *Physcomitrella patens* has no RecQ1 or RecQ3 genes but two RecQsim paralogs, and Arabidopsis lacks a RecQ6 gene but has two RecQ4 paralogs. All the subfamilies appear to be represented in Chlamydomonas, although only a mutant of the RecQ5 subfamily gene (Cre15.g634701; homologous to human *RECQL4*), which is unable to undergo cell division, has been described (Tulin and Cross 2014). These findings suggest that neo- and subfunctionalization may occur in RecQ helicase evolution and that orthologous proteins may not have identical functions in different species. In human, RECQL5 downregulation results in genomic instability and chromosomal rearrangements, and *recql5* mutants are associated with tumorigenesis (Lu and Davis 2021). However, Arabidopsis *recq3* mutants were viable and had no growth abnormalities, although this does not rule out longer term genomic instability (Röhrig et al. 2018). It remains to be seen if the deletion of *RECQ3* in Chlamydomonas can explain the genomic instability of CC-503, and it will likely never be known if this specific deletion was caused by mutagenesis or arose later in culture.

Finally, we also identified a candidate for the cell wall-less phenotype. A 6.0 kb deletion on chromosome 1 almost entirely removed a putative prolyl 4-hydroxylase (*P4H*) gene (Cre01.g800047; Supplemental Figure S12). P4Hs catalyze the formation of 4-hydroxyproline (Gorres and Raines 2010), a major post-translational modification of the hydroxyproline-rich glycoproteins (HRGPs) that comprise the Chlamydomonas cell wall (Woessner and Goodenough 1994; Sumper and Hallmann 1998). The Chlamydomonas genome encodes more than 20 putative P4Hs, and although their specific roles are generally unknown, they have different patterns of expression and are unlikely to be redundant. Keskiaho et al. (2007) showed that the knockdown of one, *P4H-1* (now annotated as *PFH12*; Cre03.g160200), was sufficient to induce abnormal cell wall assembly. Notably, the deleted gene in CC-503 has one paralog, *PFH5* (Cre01.g014650; 76% aa identity), immediately downstream that appears to be intact, although its regulation may be affected by the deletion. It is therefore unclear whether the loss of Cre01.g800047 can be responsible for the *cw* phenotype alone. Indeed, as introduced, more than one mutation may underlie the loss of the cell wall (Davies 1972; Hyams and Davies 1972).

### CC-4532 also carries structural mutations

We also identified 10 non-TE structural mutations unique to CC-4532 v6, predicted to disrupt eight genes (Supplemental Table S11, Supplemental Figure S7). The largest mutations were both duplications, of 24.5 kb on chromosome 3 and 89.1 kb on chromosome 12, which together caused the duplication of 17 complete genes. Using a coverage-based approach, Flowers et al. (2015) inferred the presence of several large duplications among various laboratory strains, and it may be the case that duplications are an important source of laboratory mutation. Interestingly, three gene-disrupting insertions in CC-4532 v6 consisted entirely of a satellite, *MSAT-11_cRei*, ranging from ∼8 kb to >19 kb (two caused assembly gaps and their full length is unknown). For example, one insertion interrupted the first exon of a gene possibly encoding nicotinate phosphoribosyltransferase (Cre03.g188800), the first step of the nicotinamide adenine dinucleotide (NAD) salvage pathway. *MSAT-11_cRei* arrays consist of a 1.9 kb tandemly repeated monomer and are present on chromosomes 7 and 12 in all three available genomes, with two additional unique insertions in CC-1690 (not shown). Similarly, *MSAT-11_cRei* de novo insertions have recently been observed in experimental lines of the field isolate CC-2931 (López-Cortegano et al. 2022). There are very few observations of de novo satellite dissemination and its mechanisms are generally unclear (Ruiz-Ruano et al. 2016), although rolling circle replication and reinsertion via extrachromosomal circular DNA intermediates has been proposed (Navrátilová et al. 2008). Collectively, these results imply that all laboratory strains may harbor at least a small number of gene-disrupting structural mutations relative to the ancestral wild type.

### Transposable element proliferation and the strain history of 137c

The most remarkable feature of CC-4532 v6 was its length of 114.0 Mb, 2.9 Mb longer than the CC-1690 assembly (Table 1). While this is partly an artefact caused by redundancy between gaps and unplaced contigs (with gaps cumulatively adding 0.9 Mb to CC-4532 v6), we attribute most of the biological increase in genome size to TE insertions in the laboratory. We identified 26 TE insertions unique to CC-503 v6 (nine of which were absent in v5, suggesting recent activity; Supplemental Table S12) and 109 insertions unique to CC-4532 v6 (Supplemental Table S13, Supplemental Figure S7), which collectively involved 14 different TE families. Remarkably, 86 of the 109 CC-4532 v6 insertions were of the same 15.4 kb *Gypsy* long terminal repeat (LTR) retrotransposon (*Gypsy-7a_cRei*, Figure 8A), adding ∼1.3 Mb of unique sequence (all TE insertions ∼1.4 Mb). This family has not been reported as active among laboratory strains, and we identified no insertions of *Gypsy-7a_cRei* in CC-503 v6, where the element is present as only one partial and two full-length ancestral copies. Only 10 of the 86 insertions were predicted to disrupt coding sequence (in some cases breaking the annotated gene model, Supplemental Table S13), and we observed intergenic insertions 2.6 times more frequently than expected by chance. *Gypsy-7a_cRei* may have a mechanism of targeted insertion, or alternatively individuals carrying genic insertions may have been selected against in the laboratory. The *Gypsy-7a_cRei* Gag-Pol polyprotein contains a plant homeodomain (PHD) finger, an accessory domain found in several Chlamydomonas TEs (Perez-Alegre et al. 2005; Craig 2021) that may be involved in chromatin remodeling to minimize deleterious insertions (Kapitonov and Jurka 2003). Nonetheless, intergenic insertions may still affect gene expression, and we also observed 10 insertions into introns and 25 into UTRs (including the 3’ UTR of *TUB2*, the gene encoding beta-tubulin). While a small number of insertions were unusual (either containing less or more than one full copy), we did not observe any solo LTRs, which are formed by recombination between the LTR subparts.

We next used whole-genome re-sequencing data (Gallaher et al. 2015) to test whether *Gypsy-7a_cRei* is active in any other laboratory strains. We analyzed 14 laboratory strains, including the oldest extant strains (CC-124, CC-125, CC-1009, CC-1010, CC-1690, and CC-1691) that are parental to all typical laboratory strains. Insertions were identified by extracting read pairs where one read mapped uniquely to a non-repetitive genomic region and the other mapped to *Gypsy-7a_cRei*. These reads formed distinct coverage peaks flanking insertions (see Supplemental Table S14 for insertion coordinates). This approach retrieved 68 of the 86 *Gypsy-7a_cRei* insertions in CC-4532 v6, the difference being attributable to insertions occurring in the ∼8 years between the Illumina and PacBio sequencing, or the inability to call insertions in repetitive regions with short reads (e.g. centromeres, see Supplemental Figure S7). All strains carry at least two to four *Gypsy-7a_cRei* copies, depending on their proportions of haplotype 1 and 2, which are presumably ancestral (collectively three copies in haplotype 1 and one in haplotype 2). Six of the fourteen strains (CC-124, CC-503, CC-620, CC-1690, CC-1009, CC-1010) had only these ancestral loci, despite being propagated for over seven decades, suggesting that *Gypsy-7a_cRei* is largely quiescent. However, in a few strains, particularly those descended from 137c+, we observed massive expansions of *Gypsy-7a_cRei*, like that in CC-4532. Indeed, CC-125, the linear descendant of 137c+, had the most novel insertions of any strain (83, Figure 8B). This was unexpected, since there are no new insertions in CC-503, which was derived from 137c+ by mutagenesis. Furthermore, we found no insertions in CC-620, another direct descendent of 137c+. CC-4532 shared 19 of its 68 laboratory insertions with CC-621 (Figure 8B), which corroborates our understanding that CC-4532 and CC-621 are both subclones of NO– from Ursula Goodenough that have been separated by at least three decades. Strains CC-4286 and CC-4287 also had some shared and some unique insertions relative to CC-4532 and CC-621, indicating shared ancestry. Anecdotally, we also observed two examples where *Gypsy-7a_cRei* inserted independently in two different strains at loci that were only ∼200 bp apart, potentially supporting targeted insertion.

**Figure 8.**
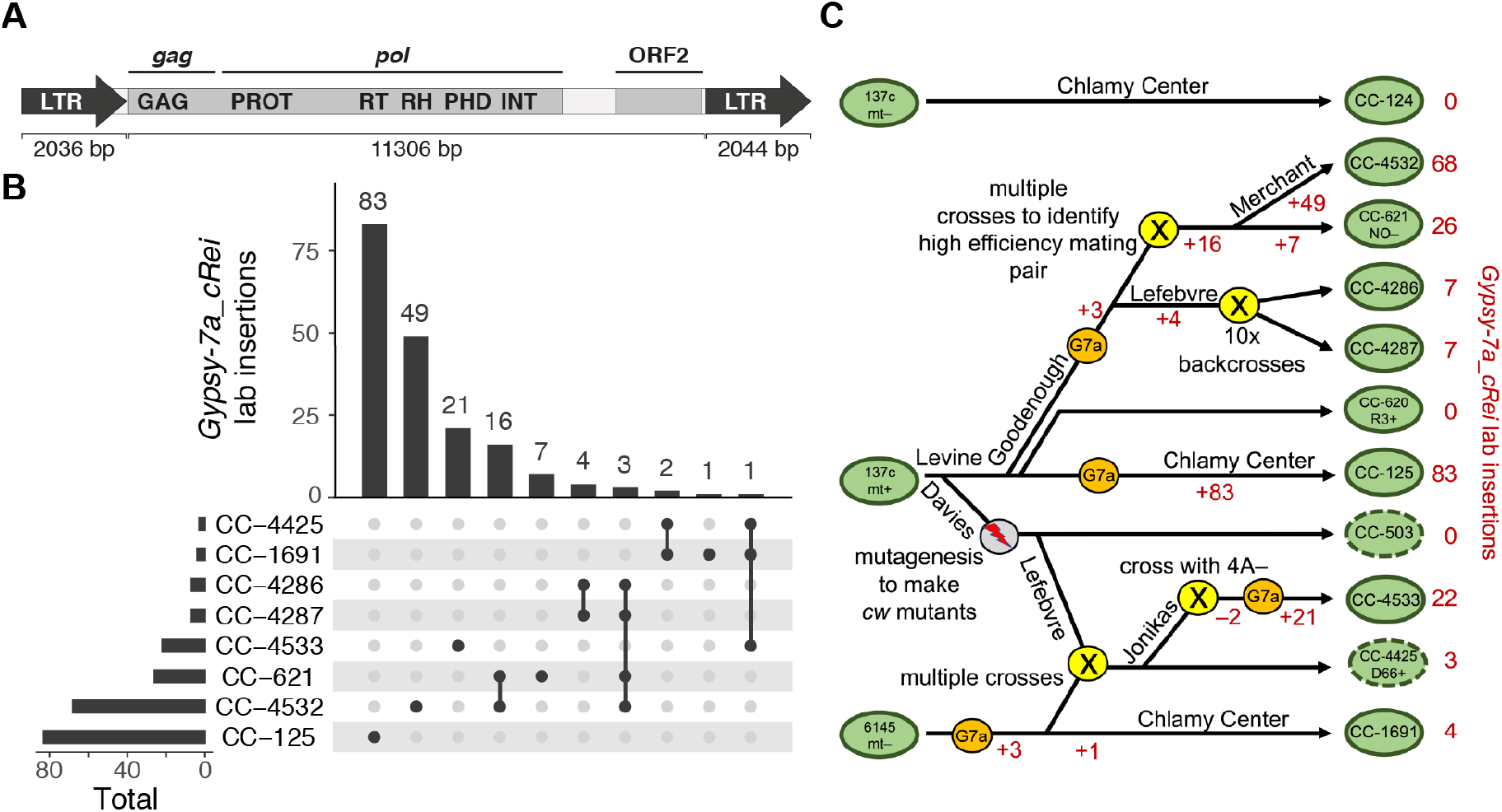
*Gypsy-7a_cRei* insertions and the strain history of 137c+. **(A)** Structure of the 15.4 kb *Gypsy* LTR retrotransposon. LTR subparts are shown as block arrows, note that the left LTR is missing the final 8 bp of the right LTR. The two ORFs are highlighted within the 11.3 kb internal section and the *gag* and *pol* sections of the polyprotein are highlighted. Protein domains: GAG = group-specific antigen, PROT = pepsin-like aspartate protease, RT = reverse transcriptase, RH = RNAse H, PHD = plant homeodomain finger, INT = integrase. **(B)** Upset plot (Lex et al. 2014) showing the number of shared and strain-specific laboratory insertions of *Gypsy-7a_cRei* in select laboratory strains. Ancestral copies of *Gypsy-7a_cRei* are excluded. **(C)** Schematic representing a putative strain history of several inter-related laboratory strains. Presented is the most parsimonious interpretation of the shared and independent insertions **(B)** coupled with known strain histories. Green ovals represent strains as indicated, with a dashed line indicating cell wall defective strains. A gray circle indicates MNNG mutagenesis that produced *cw* mutants. Yellow circles indicate crosses as labeled. Orange circles indicate likely activation of *Gypsy-7a_cRei* (“G7a”). Changes in the net number of *Gypsy-7a_cRei* loci due to addition by retrotransposition (+) or loss during crossings (–) are indicated in red. The names of several key Chlamydomonas researchers (R.P. Levine, D.R. Davies, U.W. Goodenough, P.A. Lefebvre, S.S. Merchant) are indicated where relevant.

We attempted to reconcile the distribution of the *Gypsy-7a_cRei* laboratory insertions with previously described strain histories (Pröschold et al. 2005; Gallaher et al. 2015). The result of this analysis is presented as the proposed strain history in Figure 8C. Since all insertions were unique to CC-125, we hypothesize that *Gypsy-7a_cRei* became active in the 137c+ culture that became CC-125 after being separated from the cultures that became CC-503 and CC-620, which occurred several decades ago. *Gypsy-7a_cRei* became active independently again in a strain from the laboratory of Ursula Goodenough (CC-621), and it remains active and continues to expand in strains derived from that strain, e.g. CC-4286 and CC-4287 from Paul Lefebvre and CC-4532 from Sabeeha Merchant. A third reactivation of *Gypsy-7a_cRei* likely occurred in Ruth Sager’s 6145 strain, which eventually became CC-1691. This event contributed novel insertions to strain D66+ (CC-4425), which in turn contributed a single laboratory insertion to Martin Jonikas’ strain, CC-4533. This last strain, the parental strain of the Chlamydomonas Library Project (CLiP), may represent a fourth reactivation of *Gypsy-7a_cRei* (or an increase in transposition frequency), since it carries 21 private insertions despite being separate from CC-4425 by approximately a decade.

Aside from *Gypsy-7a_cRei*, the most active TE family was *MRC1*, with 17 insertions in CC-503 v6 and 16 insertions in CC-4532 v6 (Supplemental Tables S12 and S13). *MRC1* was originally described as a non-autonomous LTR element (Kim et al. 2006), however we recently reclassified it as a non-autonomous member of the newly described *Chlamys* clade of *Penelope*-like elements (Craig et al. 2021b). Gallaher et al. (2015) and Neupert et al. (2020) reported activity of *MRC1*, and it may generally be one of the most active TEs in the laboratory. We identified four active DNA transposons that have been described previously, namely one insertion each of *Gulliver* (Ferris 1989), *Tcr1* (Schnell and Lefebvre 1993) and *Tcr3* (Wang et al. 1998) (*hAT*, *Kyakuja* and *EnSpm* superfamilies, respectively), and three insertions of the non-autonomous *hAT* family *Bill* (Kim et al. 2006). The eight remaining TEs have only been described in Repbase (Bao et al. 2015) or the more recent Chlamydomonas TE library (Craig 2021). Taken collectively, these results suggest that TE activity between laboratory strains can be highly heterogenous, with the potential for rapid TE proliferation to cause significant increases in genome size and to disrupt genic sequence. It is presently unclear why suppression of *Gypsy-7a_cRei* is unstable in certain strains, and why this family exhibits a far higher transposition frequency than other active TEs upon activation. Similar copy number variation among laboratory strains has been reported previously for the active nonautonomous DIRS retrotransposon *TOC1* (Day et al. 1988), although curiously we did not find any de novo insertions of this element in CC-503 v6 nor CC-4532 v6.

### Transposable element genes and the *TET/JBP* gene *CMD1*

Although the 14 active TE families observed in CC-4532 v6 and CC-503 v6 may be representative of TEs active among laboratory strains, they represent only a fraction of the TEs that are active in the species. Our recent annotation of TEs identified more than 220 families in the Chlamydomonas genome, with almost all TE copies showing signatures of recent activity in the species (Craig 2021). This result means that if a TE contains a gene (or genes), the coding sequence is unlikely to be disrupted by mutation. While TE genes may have unusual features (e.g. ribosomal frameshifts, especially in retrotransposons), many are readily identified by gene prediction algorithms. Unknowingly including TE genes within annotations can confound analyses, such as analyses of methylation, chromatin states or small RNA targeting, where substantial differences may be expected between non-TE and TE genes. Annotation projects therefore generally aim to exclude as many TE genes as possible, while highly curated annotations of model organisms may include TEs as defined entities. We applied a balanced approach in the v6 annotations, in which we incorporated a set of highly supported TE genes, while filtering out lower confidence genes that would require curation.

We previously identified ∼1,000 genes in v5.6 that are likely part of TEs (Craig et al. 2021a). When comparing v5.6 genes and TE coordinates, the distribution of their intersect is highly bimodal; 1,023 genes have a >30% overlap between their coding sequence (CDS) and TEs, and 908 genes have >80% overlap (Figure 9A). We obtained similar distributions in the preliminary v6 annotations, indicating that most genes can be cleanly divided into TE and non-TE subsets. To designate high-confidence TE genes, we required a gene with a high CDS-TE overlap to have either sequence similarity to a known TE-encoded protein or a functional domain. This analysis resulted in 810 TE genes being included in CC-4532 v6.1 (Figure 9B, Table 2) and 647 in CC-503 v6.1 (Supplemental Figure S13), which are integrated in the associated GFF3 files under the field “transposable_element_gene”. Users should be aware that these TE gene sets are not exhaustive, and projects requiring TE coordinates in general should use annotations derived from the dedicated repeat library (Supplemental Dataset 4).

**Figure 9.**
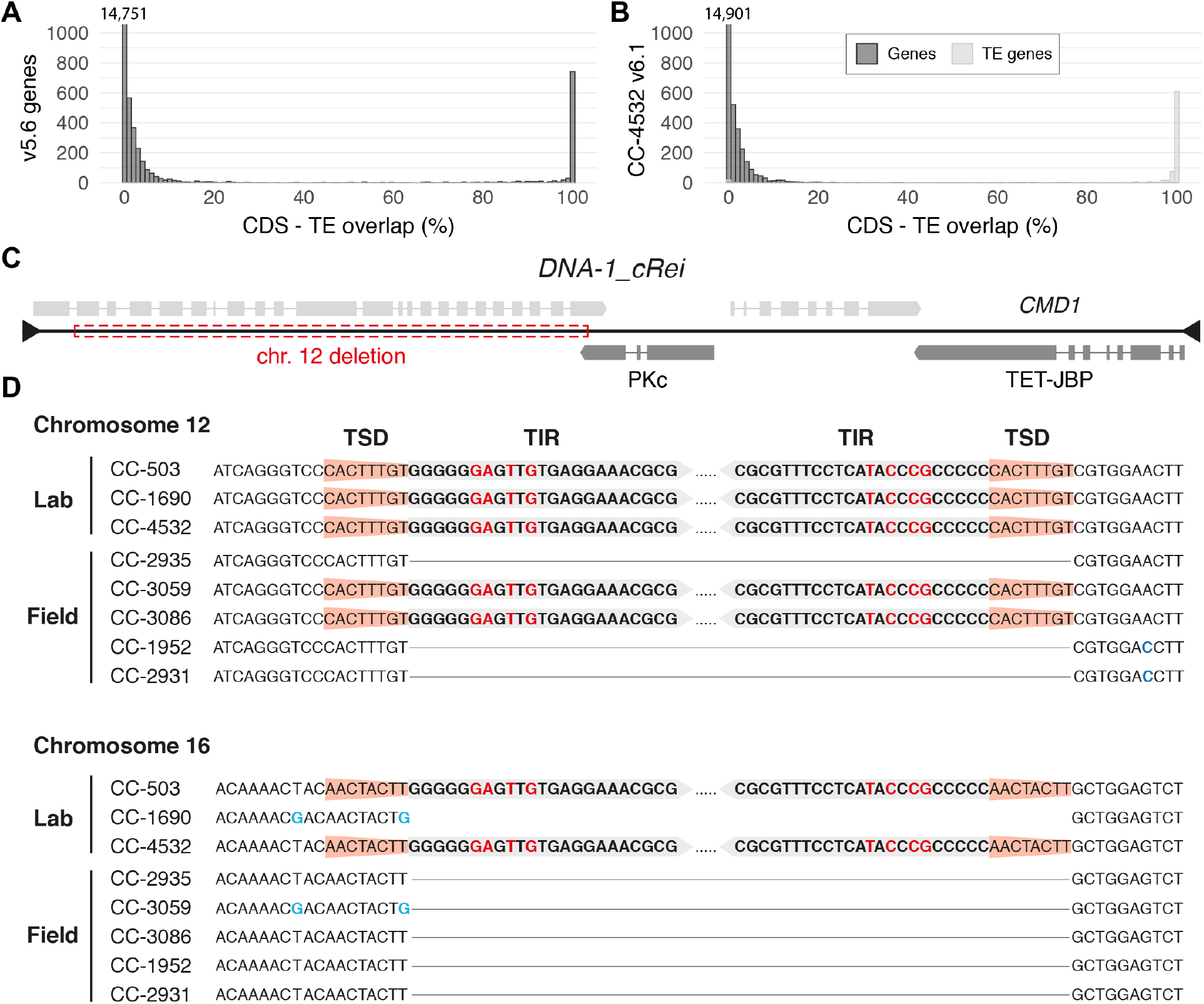
Transposable element genes and *CMD1*. **(A)** Overlap between gene coding sequence (CDS) and TEs in v5.6. The number of genes with 0% overlap is written above the first bar. **(B)** Overlap between gene coding sequence (CDS) and TEs in CC-4532 v6.1. Genes were split into non-TE and TE genes. **(C)** Schematic representation of the *DNA-1_cRei* transposon that contains *CMD1*. Gene models were derived from Iso-Seq (Supplemental Figure S14), forward strand genes are shown above black line, reverse strand genes below. Protein domains are highlighted (PKc = protein kinase catalytic domain, TET-JBP = ten-eleven translocation/J-binding protein). The ∼8 kb region deleted from the chromosome 12 copy is shown by the red box. **(D)** Sequence context of the chromosome 12 and 16 copies of *DNA-1_cRei*. The 8 bp target site duplications (TSDs) and the first 23 bp of the 67 bp terminal inverted repeats (TIRs) at each terminus are highlighted. Mismatches between the TIRs are shown in red text, and single nucleotide variants present among laboratory and field isolates are shown in blue text. Regions lacking insertions are shown as solid black lines. Variants were manually curated from PacBio and Illumina data.

Most of the TE genes encode proteins with typical TE domains, such as reverse transcriptases and integrases. These genes may be included or excluded in analyses as desired. TE-encoded proteins are capable of important and wide-ranging interactions with their host genomic and cellular environments (Cosby et al. 2019), and we were motivated to include TE genes due to their potential functional relevance. The *TET/JBP* (ten-eleven translocation/J-binding protein) gene *CMD1* (5mC-modifying enzyme, Cre12.g553400) is such an example, as the corresponding protein is responsible for DNA demethylation via a novel biochemical pathway, converting 5mC to the previously undescribed epigenetic mark C^5^-glyceryl-methylcytosine (5gmC) using a vitamin C co-substrate (Xue et al. 2019; Li et al. 2021). Genome-wide 5mC levels doubled in a *cmd1* mutant relative to its isogenic wild type, which resulted in lower expression of certain genes and susceptibility to photodamage (Xue et al. 2019). Notably, TET-JBP proteins have been described in *KDZ* DNA transposons of fungi, where their DNA demethylation activity may play a role in self-regulation and the regulation of other TEs (Iyer et al. 2014). Similarly, we found that Chlamydomonas *CMD1* appears to be part of a previously undescribed DNA transposon, *DNA-1_cRei*. This 18.4 kb element is highly unusual, since it contains four expressed genes (including *CMD1*) although none encodes a recognizable transposase and we were thus unable to classify it to any known clade of DNA elements (Figure 9C, Supplemental Figure S14). Nonetheless, *DNA-1_cRei* possesses all the characteristics of a DNA transposon, namely 67 bp imperfect terminal inverted repeats, precise insertion polymorphisms among both laboratory strains and field isolates, and 8 bp target site duplications (Figure 9D). In CC-4532 v6 and CC-503 v6, *DNA-1_cRei* is present in two copies, one on chromosome 12 that features an ∼8 kb internal deletion, and a second presumably full-length copy on chromosome 16 (containing a second copy of *CMD1*, Cre16.g654100). The second copy is absent in CC-1690 (Figure 9D, Supplemental Figure S14), although this is a haplotype 2 region in CC-1690 and the absence is likely ancestral and unrelated to TE activity in the laboratory. In addition to the two *CMD1* copies, the Chlamydomonas genome contains at least ten other *TET/JBP* genes (Aravind et al. 2019), most of which are part of two Helitron TE families (*Helitron2-2_cRei* and *Helitron2-3_cRei* (Craig 2021)). It is possible that the low methylation found outside of hypermethylated regions is partly an outcome of the presence of multiple TEs containing TET/JBP genes. This example demonstrates that TE genes can potentially underlie important biological processes. These genes can vary in copy number between strains, and their inclusion in the wider annotation should facilitate the study of novel aspects of Chlamydomonas biology.

### The present and future of the Chlamydomonas Genome Project

For almost two decades, the Chlamydomonas Genome Project has been based on the *mt+* strain CC-503. In version 6, we have presented near-complete assemblies for both CC-503 and the *mt–* strain CC-4532. Following the discovery of numerous structural mutations affecting CC-503, CC-4532 v6 has been chosen to serve as the reference genome. Despite its replacement, CC-503 v6 remains a valuable resource, especially for the *MT+* allele and the existing organelle genomes that were appended to CC-4532 v6.

It is now clear that laboratory strains can differ extensively from each other, both genetically and phenotypically. Most of this variation stems from the mosaic of two haplotypes that comprise the genome of each strain (Gallaher et al. 2015). These developments have led to the “know thy strain” maxim: researchers are encouraged to consider the genetic differences that exist between the reference genome and the strains used in experimental work (Salomé and Merchant 2019). Our results suggest that these differences should not only be considered with respect to ancestral variation between the haplotypes, but also to derived variation arising by laboratory mutation. Although CC-503 may be an extreme case, the CC-4532 genome harbors 10 structural mutations and more than 100 TE insertions. Indeed, analyses by Gallaher et al. (2015) and Flowers et al. (2015) previously inferred the presence of many derived structural variants among strains, including several large duplications. While many of the most characteristic laboratory phenotypes were caused by mutations (e.g. *nit1* and *nit2*), it is likely that all strains have experienced unique structural mutations (including TE proliferation at various rates), many of which disrupt genes. It is also possible that independently maintained cultures of the same strain may differ due to independent mutations, which makes it desirable to keep frozen stocks of any strain used for long-term projects, e.g. generation of mutant libraries. Laboratory strains have been maintained clonally for as many as 75 years and mutations are an unavoidable consequence, especially if strains are evolving under relaxed selection. Recent work has in fact demonstrated that an unexpectedly large proportion of mutations in Chlamydomonas are beneficial under laboratory conditions, potentially due to many genes being dispensable in such an artificial environment (Böndel et al. 2019; Böndel et al. 2022). It has also been estimated that ∼5-10% of all de novo mutations in Chlamydomonas experimental lines are >50 bp (López-Cortegano et al. 2022), supporting a prominent role for structural evolution in the laboratory. The implications of “laboratory domestication” have been considered in other model systems such as *Caenorhabditis elegans* (Sterken et al. 2015), and laboratory mutations should be carefully considered when evaluating experimental results.

Although CC-4532 v6.1 is the highest quality release to date, there remains room for improvement, particularly in gene annotation. The availability of various ‘omics datasets for Chlamydomonas and related species continues to grow, presenting opportunities to integrate new annotation evidence. This may be especially important for the structural and functional annotation of alternative isoforms and polycistronic loci, which have not been considered in depth in this work. Work is ongoing to incorporate various additional annotations as browser tracks on Phytozome (e.g. noncoding RNA genes), and we re-iterate that genome users are encouraged to contact with any curated improvements.

With the continuous developments in long-read sequencing, we are entering an exciting era of Chlamydomonas genomics. Complete “telomere-to-telomere” Chlamydomonas genomes are within reach, and a pan-genome project has recently been initiated, aiming to produce genomes for several additional laboratory strains and field isolates (https://www.chlamycollection.org/the-chlamydomonas-pan-genome-needs-you/). As demonstrated herein, many insights can only be gleaned by comparing the genomes of different strains, and we can expect substantial benefits from sequencing additional strains and isolates moving forward. With respect to the two laboratory haplotypes, a “laboratory pan-genome” could be produced where all haplotype 1 and 2 regions are represented, capturing all ancestral variation present among laboratory strains. This could potentially take the form of consensus assemblies for each haplotype, with data from several strains used to infer the ancestral state at the time of isolation and to identify derived mutations in specific strains. Furthermore, similar to resources developed for several important plants (Bayer et al. 2020), the species-level pan-genome aims to incorporate the far greater diversity present among Chlamydomonas field isolates (Flowers et al. 2015; Craig et al. 2019). Such prospects are expected to reveal novel aspects of Chlamydomonas biology, continuing the development of the species as an integral model in plant and algal biology.

## METHODS

### Strains and DNA sequencing

CC-503 was obtained from the Chlamydomonas Resource Center in 2012. CC-4532 has been propagated in Sabeeha Merchant’s group since the late 1990s (see Gallaher et al. (2015)), when it was received from Ursula Goodenough. Cultures were grown as described previously (Gallaher et al. (2015).

Genomic DNA was extracted from frozen cell pellets and used for library preparation and sequencing at the Joint Genome Institute. Sequencing was performed on the PacBio Sequel platform in CLR (continuous long reads) mode, generating ∼127x and 176x coverage for CC-503 and CC-4532, respectively (CC-503 mean read length 3.58 kb; CC-4532 mean read length 9.88 kb). Additional Illumina sequencing was performed on the HiSeq2000 platform (150 bp paired-end reads, ∼400 bp insert) to ∼50x (CC-503) and 55x (CC-4532) coverage as reported in Gallaher et al. (2015).

### Assembly of CC-4532 v6 and CC-503 v6 genomes

Preliminary contig-level assemblies were produced from the PacBio datasets. CC-503 was assembled using MECAT version 1.1 (genomeSize=130000000 ErrorRate=0.02 Overlapper=mecat2asmpw) (Xiao et al. 2017) and CC-4532 using Canu version 1.8 (genomeSize=120000000) (Koren et al. 2017). Reads were mapped to the raw assembly using BLASR, and error correction was performed using a single pass of Arrow correction from the GenomicConsensus toolkit. The remaining homozygous SNP/indel consensus errors were corrected using the strain-appropriate Illumina data. Illumina reads were aligned using bwa mem (Li 2013) and homozygous SNPs and indels were identified using the GATK’s UnifiedGenotyper tool (McKenna et al. 2010), and were subsequently corrected. The corrections were verified by mapping the Illumina reads to corrected consensus sequence.

We used the CC-1690 assembly (O’Donnell et al. 2020) to scaffold the preliminary contigs of each assembly to chromosomes. Contigs were mapped to the CC-1690 assembly using minimap2 v2.17 (-ax asm5) (Li 2018) to produce PAF (Pairwise mApping Format) files. These mapping data were used to manually order and orientate uniquely mapping contigs (i.e. the majority of the contig received a mapping quality of 60) relative to each CC-1690 chromosome. Any inconsistencies between the contigs and CC-1690 chromosomes were inspected against the raw PacBio reads from the relevant strain using IGV v2.7.2 (Robinson et al. 2011). In a small number of cases a misassembled contig was split, while for CC-503 some genuine inconsistencies caused by structural mutations were supported by the raw reads and maintained. Several short contigs that mostly contained satellite DNA were manually removed since they appeared to duplicate a region already assembled on a larger contig. Other short contigs entirely consisting of subtelomeric repeats, which generally did not map uniquely, were assigned to chromosome termini by specific alignment and phylogenetic analysis (see Chaux-Jukic et al. (2021)).

Gap lengths between contigs were estimated relative to CC-1690 and the appropriate number of Ns was inserted between contigs. Occasionally the estimated “gap” length was negative, suggesting redundant sequence at the termini of neighboring contigs. These contig termini were compared, trimmed to remove redundant sequence, and subsequently merged where possible. Arbitrary gaps of 100 Ns were inserted between contigs which could not be successfully merged.

### Repeat annotation

TE sequence was identified in each assembly by providing the latest Chlamydomonas repeat library to RepeatMasker v4.0.9 (Smit et al. 2013-2015). This library features updated consensus models for all Chlamydomonas repeats available in Repbase (https://www.girinst.org/repbase/) together with >100 newly curated repeats (Craig 2021). Any putative TE divergent by >20% from its consensus sequence was removed. *ZeppL* clusters were identified as the span from the first two consecutive *ZeppL-1_cRei* copies to the final two consecutive *ZeppL-1_cRei* copies on each chromosome (except for chromosome 15 where three distinct clusters were observed, see Results).

Microsatellites and satellite DNA were primarily identified using Tandem Repeats Finder (Benson 1999) with parameters “2 7 7 80 10 50 1000” (i.e. a minimum alignment score of 50 and a maximum monomer size of 1000 bp). Tandem repeats consisting of ≥2 monomers were split to microsatellites (monomers <10 bp) and satellite DNA (monomers ≥10 bp), and if a region was called as both, precedence was given to satellite DNA since shorter monomers are frequently nested within larger ones. Satellite DNA annotations were supplemented with curated satellites identified by RepeatMasker from the repeat library, several of which have monomers longer than the detection limit of Tandem Repeats Finder.

Genome-wide CG methylation was quantified for the CC-1690 assembly following Chaux-Jukic et al. (2021). Briefly, the raw signal of the CC-1690 Nanopore reads (i.e. fast5 files) generated by Liu et al. (2019) were mapped to the CC-1690 assembly using Tombo (https://nanoporetech.github.io/tombo/) and CG methylation was called using DeepSignal (Ni et al. 2019).

### Validation of assembly improvements

Misassemblies in the v5 assembly were identified by mapping the v5 contigs to the chromosomal CC-503 v6 and CC-4532 v6 assemblies using minimap2 as described above. Genomic coordinates of intra- and inter-chromosomal inconsistencies were assessed manually from the PAF files and converted to input files for Circos (Krzywinski et al. 2009) and karyoploteR (Gel and Serra 2017) to produce figures 1, 2 and S1.

To enable convenient liftover of coordinates between assemblies, a 5-way Cactus whole-genome alignment (WGA) (Armstrong et al. 2020) was produced including the v4, v5, CC-503 v6, CC-4532 v6 and CC-1690 assemblies. Each assembly was soft masked for repeats by providing coordinates of TEs and tandem repeats (see above) to BEDtools v2.26.0 maskfasta (-soft) (Quinlan and Hall 2010). An arbitrary guide tree for Cactus was provided as “(CC-4532_v6:0.001,(CC-1690:0.001,(CC-503_v4:0.001,(CC-503_v5:0.001,CC-503_v6:0.001):0.001):0.001):0.001)”, and all assemblies were set to reference quality. Liftover of genomic coordinates in BED (Browser Extensible Data) format could then be performed between any two assemblies in the HAL (Hierarchical ALignment) format WGA using the HAL tools command halLiftover (Hickey et al. 2013). This approach was used to convert v5 coordinates of hypermethylated regions (Lopez et al. 2015) to CC-4532 v6 (Figure 1) and CC-1690 (Supplemental Figure S3) coordinates. Coordinates of v5 assembly gaps were also converted to CC-4532 v6 coordinates to investigate the sequence properties of filled gaps in the updated assembly (see Figure 4).

The genotyping data from Kathir et al. (2003) were kindly provided by Paul Lefebvre. The genomic coordinates (v5 assembly, as chromosome and position, in bp) were determined for all markers by BLAST search in Phytozome using the sequence deposited for each marker (https://www.chlamycollection.org/BAC/MARKER_index.htm), or by keyword search using the gene name in Phytozome. The markers were then ordered based on their assigned v5 chromosome and position. All genotyping data were assembled into a tab-delimited file and used as input for R/QTL (Broman et al. 2003) with the functions *read.cross*, *est.rf*, *plotMap*, *plotRF*, and *summaryMap*. The genotyping data for the CC2935 × CC2936 progeny (12 full tetrads) were obtained from Liu et al. (2018). Since the genotypes were encoded as either 1 or 2, a matrix (of the same size as the genotype matrix) was calculated whereby each n+1 position received the difference between genotype at the n+1 position and the genotype at position n. Any SNP with a value not equal to zero was retained to estimate the genetic map, as described above. The genomic coordinates of mismapped markers or SNPs were corrected based on the CC-503 v6 or CC-1690 assemblies before re-running the genetic map construction, as above. The quality of the assemblies was assessed by plotting the recombination frequencies across the entire genome (*plotRF*) and by calculating the total length of the genetic map (*summaryMap*).

### Preliminary structural annotations

Protein-coding genes for the CC-4532 v6 and CC-503 v6 assemblies were annotated using a combination of tools. Input data were ∼1.6 billion 150 bp paired-end RNA-seq reads from the JGI Gene Atlas (strain CC-1690), ∼6.4 million 454-sequenced ESTs generated in previous versions (CC-1690) and ∼1.6 million PacBio Iso-Seq reads (pooled samples from CC-4532, CC-5390, CC-4348, CC-4349, CC-4565, CC-4566 and CC-4567, see Gallaher et al. (2021)). The Gene Atlas samples will be described by Sreedasyam et al. (unpublished) and Schmollinger and Merchant (unpublished), and can be browsed at https://phytozome-next.jgi.doe.gov/geneatlas/. Specifically for the CC-4532 v6 annotation, ∼520 million unpaired 50 bp RNA-seq reads were included that were generated from CC-4532 grown under a range of conditions including heterotrophic and photoautotrophic growth, and in Fe replete and Fe limited media (NCBI SRA accessions PRJNA842032 and PRJNA717804). The RNA-seq and 454 reads were first assembled using PERTRAN (Shu et al. 2013), which conducts genome-guided transcriptome short-read assembly via GSNAP (Wu and Nacu 2010) and builds splice alignment graphs after alignment validation, realignment and correction. Iso-Seq circular consensus sequencing (CCS) reads were corrected and collapsed using a pipeline that aligns CCS reads to the genome with GMAP (Wu and Watanabe 2005), performs intron correction for small indels in splice junctions (if any), and clusters alignments where all introns are shared for multi-exonic transcripts, or have 95% overlap for single-exon transcripts. A combined assembly of all transcriptomic data was then produced using PASA (Haas et al. 2003), yielding 287,891 assembled transcripts for CC-4532 v6 and 293,991 of CC-503 v6.

Putative gene loci were determined using PASA and its associated transcriptome assembly, or by EXONERATE (Slater and Birney 2005) alignments of proteins from *Chlamydomonas incerta, Chlamydomonas schloesseri, Edaphochlamys debaryana, Volvox carteri, Gonium pectorale, Dunaliella salina, Chlamydomonas eustigma, Monoraphidium neglectum, Chlorella variablis, Coccomyxa subellipsoidea, Bathycoccus prasinos, Auxenochlorella protothecoides, Micromonas commoda, Arabidopsis thaliana, Glycine max, Oryza sativa, Setaria viridis, Solanum lycopersicum, Vitis vinifera*, human, mouse and Swiss-Prot eukaryote proteomes. Transcripts and proteins were aligned against assemblies after softmasking TEs. Gene models were predicted using four tools: either directly from the PASA transcripts by identifying ORFs, or using FGENESH+/FGENESH_EST using parameters trained for *V. carteri* (Salamov and Solovyev 2000), AUGUSTUS v3.3.3 using “chlamy2011’’ species parameters (Stanke et al. 2006; Stanke et al. 2008) or EXONERATE using the protein2genome:bestfit model (Slater and Birney 2005). At a specific locus, the most well-supported gene model from one of the above approaches was retained. Gene models were scored positively for a total score of 7, weighted by the following criteria. First, transcriptomic support contributed a maximum score of 3: a score of 0-1 based on the percentage of exon coverage by the PASA assembly, a score of 0-1 for intron support ratio (the number of introns in coding sequence with both splice sites supported relative to the total number of introns in coding sequence), and a score of 0-1 for splicing support ratio (the number of supported splice sites in coding sequence relative to the total number of splice sites in coding sequence). Second, protein homology support contributed a maximum score of 3: 0-1 for the “Cscore” (a ratio of the blastp score between the predicted protein and its best hit in the proteomes listed above, and the blastp score of that hit against the predicted protein), 0-1 for protein homology coverage (the length of alignment between the predicted protein and its best blastp hit, relative to the mean length of the predicted protein and its best blastp hit), and 0-1 for the percentage of predicted peptide coverage by EXONERATE protein alignments. Finally, a score of 0.3 was awarded for the presence of a start codon, 0.2 for a stop codon, and 0.5 for full-length transcriptome support (every intron in coding sequence supported). Models were negatively scored by 0-1 for percent overlap between coding sequence and TEs. The selected gene models were then subject to improvement using PASA, which included adding UTRs, splicing correction and addition of alternative transcripts.

Finally, poorly supported PASA-improved transcript models were filtered out using several of the metrics described above. Transcript models were retained if they had <20% overlap between coding sequence and TEs, and they had a Cscore >=0.5 and protein homology coverage >=0.5, or they received transcriptomic support (>=0.8 for intron support ratio). For gene models where coding sequence had >=20% overlap with TEs, a Cscore >=0.9 and protein homology coverage >=0.7 were required. Pfam domains were then identified for each retained protein using InterProScan (Jones et al. 2014). Incomplete gene models (missing a start or stop codon) with a combined Cscore and protein homology score of <1.9 were removed. Any gene models with low protein homology (a combined Cscore and protein homology score <1.6) and without full-length transcriptome support were also removed. Finally, short single-exon models (<300 bp) that lacked recognized protein domains and had poor expression (<15 reads) were removed.

### Curation of structural annotations

The preliminary gene models for CC-4532 v6 and CC-503 v6 were subject to several steps of improvement. First, despite filtering out models with little homology or transcriptomic support (see above), the annotations each contained hundreds of short models with dubious coding potential. To quantify coding potential, the analyses of Craig et al. (2021a) were repeated. Briefly, for each annotation all genes were split to two classes, either “control” (genes with at least one algal homolog or a protein domain) or “test” (genes with neither a homolog nor protein domain). Algal homologs were identified using OrthoFinder v2.2.7 (Emms and Kelly 2015). Four metrics were then calculated per gene: the PhyloCSF score (Lin et al. 2011), the ratio of genetic diversity at zero-fold and four-fold degenerate sites (ρχ0D/4D), codon usage as quantified by the index of translation elongation (*I*_TE_) (Xia 2015), and a “Kozak score” that quantifies the strength of the Kozak-like sequence surrounding the start codon (Cross 2015). To run PhyloCSF, two 8-way Cactus WGAs were produced, one with CC-4532 v6 plus seven outgroup genomes (*C. incerta, C. schloesseri, E. debaryana, G. pectorale, V. carteri, Yamagishiella unicocca* and *Eudorina* sp. 2016-703-Eu-15), and second with CC-503 v6 and the same seven genomes. The guide tree was identical to that of Craig et al. (2021a), as was post-processing of the WGAs and the calculation of PhyloCSF scores. Genetic diversity (ρχ) was estimated for preliminary gene models from both CC-4532 and CC-503 using whole-genome re-sequencing data from a population sample of 17 Quebec field isolates, with SNPs called and filtered for both assemblies following Craig et al. (2019). *I*_TE_ was calculated using the optimal codon usage table previously produced from v5.6 genes (Craig et al. 2021a). To calculate Kozak scores, a Kozak-like sequence consensus logo, which summarizes the frequency of nucleotides 5 bp up- and downstream of start codons, was produced using a random half of the control set genes in each annotation with WebLogo 3 (Crooks et al. 2004). The sequence context of the start codons for genes in the test set and the other half of the control set were then scored for their match to the consensus logo, with a higher score indicating a stronger Kozak-like sequence. Combining these data, a test set gene was deemed to be low coding potential if it had a PhyloCSF score <1 and failed two of the following conditions: ρχ0D/4D >95^th^ percentile of ρχ0D/4D of the control genes, *I*_TE_ <5^th^ percentile of *I*_TE_ values for the control genes, or a Kozak-like score <0.25 (see Supplemental Figures S5 and S6). Low coding potential genes with three or more exons, or ORFs ≥900 bp (after subtracting tandem repeats), were retained in the main gene sets.

Second, analyses were performed to avoid omitting from the v6 annotations any well-supported genes from previous annotations. A dataset was produced that included v4 genes that were absent in the v5 annotations (Blaby and Blaby-Haas 2017), the entire v5.6 gene set, and 142 high confidence genes identified by Craig et al. (2021a) in the v5 assembly that were missing from v5.6. Any genes with coding sequence overlapping >30% with TEs were removed and all genes were required to have strong evidence of coding potential (either an algal homolog, a recognized protein domain or a PhyloCSF score ≥100). The coding sequence coordinates of each gene were converted from the source assembly (v4 or v5) to the target assemblies (CC-4532 v6 and CC-503 v6) using halLiftover and the 5-way WGA (see above). We considered genes to be potentially missing in the target genome if ≥90% of coding sequence could be successfully lifted over, and <10% of the converted coordinates overlapped existing coding sequence in the target genome. Additionally, both v6 preliminary annotations were checked against each other i.e. all CC-503 v6 preliminary gene models with strong coding potential were converted to CC-4532 v6 coordinates and checked for overlap with CC-4532 gene models, and vice versa. Any genes from a v6 annotation were given priority over models from v5 or v4 (i.e. in cases where they overlapped in the target v6 assembly).

We attempted to add the resulting missing genes to the relevant assembly using three approaches. If the coding sequence liftover was entirely one-to-one (i.e. every site within each exon perfectly aligned) the gene was simply incorporated based on the converted coordinates. If not (i.e. ≥90% was lifted over, but <100%), we searched for the gene in question among the discarded low-scoring gene models from the preliminary annotations (see above). A match for a missing gene was defined as any discarded model with a significant blastp hit (e-value ≤0.01 and percent identity ≥95%) that had ≥80% intersect between coding sequence coordinates in the target assembly. If neither approach was successful, the transcript of the missing gene was mapped as a cDNA to the relevant target assembly using GMAP (parameters: “—cross-species, --max-intronlength-ends 200000, -z sense_force” (Wu and Watanabe 2005). Gene models were included if the mapped cDNA had a valid ORF in the target genome.

Finally, we hand curated a number of gene models, including those for selenoproteins, all polycistronic loci described by Gallaher et al. (2021), and *MT* genes. Any gene that was either added by the automated pipeline described above or was manually curated was given the suffix “_EX” in the associated locus IDs.

### Annotation of transposable element genes

In the preliminary annotations, gene models that had >20% coding sequence overlap with TEs were filtered out, unless they had very strong sequence identity relationships (note that many genomes of other algal species do include TE genes in their main annotations, and therefore strong sequence identity is not unexpected). These genes with homology evidence were subsequently separated from the main annotation if their coding sequence overlapped with TEs by ≥80%, or if they encoded proteins in which >30% of Pfam domains were TE-associated. For each annotation, this gene set was then reduced to a conservative set of TE genes. First, a gene was retained if it had any protein domain (i.e. not only TE-associated domains, see Figure 9). Second, each protein was queried against the Repbase TE protein database using blastp and retained a) if percent identity between the query and hit was ≥60%, if the best hit had an e-value:′,0.001 and its length spanned ≥50% of both query and target lengths, or b) if percent identity was <60%, if the hit spanned ≥20% of each length. For simplicity, this conservative set of TE genes was manually reduced to one transcript isoform based on inspection of transcriptomic support. Genes that did not meet these criteria were discarded (see examples in Figure S14).

### ChIP-seq and proteomics

Intergenic H3K4me3 ChIP-seq peaks called against the v5 assembly were retrieved from Strenkert et al. (2021). Peak coordinates from the three time points in their experiment were merged and subsequently converted to CC-4532 v6 coordinates using halLiftover (see above, a peak was defined as successfully lifted over if ≥90% of sites were converted). Following Strenkert et al. (2021), distance from the midpoint of each peak to the nearest TSS was calculated, and a peak was assigned to a TSS if it was within 750 bp. Peaks that were still classed as intergenic after this analysis were compared to the TE annotation and conservatively called as TE-associated if ≥80% of sites within the peak overlapped a single TE copy.

The proteomics analysis was performed as in Gallaher et al. (2018), using datasets generated in that study. Briefly, peptides were identified by mass spectrometry and compared to the v5.6 and CC-4532 v6.1 predicted proteins. The total number of gene models encoding proteins with at least one assigned peptide was estimated, as was the total number of unique peptides assigned to each annotation, and the total number of peptides assigned overall.

### Identification of structural mutations and transposable element insertions

Structural variants (i.e. >50 bp) were called between the CC-503 v6 and CC-4532 v6 assemblies using MUM&Co (O’Donnell and Fischer 2020), which identifies putative variants from MUMmer alignments (Kurtz et al. 2004). MUM&Co was run on each chromosome individually, and for chromosomes 2 and 9 the CC-503 v6 chromosomes were split at the translocation breakpoints and the relevant parts of each chromosome were included. All variant calls were then visualized and curated in IGV by comparing the CC-503 v6, CC-4532 v6 and CC-1690 assemblies (using alignments produced by minimap2, as performed above) and raw PacBio reads. Variants called within tandem repeats or within regions where CC-4532 carried haplotype 2 were not considered. Confirmed variants were polarized as mutations by comparison of the three assemblies i.e. the allele present in two assemblies (one of the v6 assemblies and CC-1690) was assumed to be ancestral. Structural mutations unique to CC-1690 were not called.

Structural mutations identified in CC-503 v6 were subsequently compared to past assembly versions and were called as consistent (present) or inconsistent (absent). Genes putatively affected by structural mutations were identified from the assembly and annotation featuring the ancestral state i.e. genes from the CC-4532 v6.1 annotation were identified at the regions overlapping CC-503 v6 structural mutations (see Supplemental Tables S10-13).

TE insertions were called as specific cases of insertion variants called by MUM&Co. All insertions were compared against the annotations derived from the Chlamydomonas repeat library (see above) and called as TE insertions where genomic coordinates of a TE perfectly intersected those of the insertion. Similarly, a small number of “deletions” were called as excision events of cut-and-paste DNA transposons (e.g. *Gulliver*).

To identify insertions of *Gypsy-7a_cRei* in laboratory strains, whole-genome re-sequencing data from 14 strains (Gallaher et al. 2015) were aligned using bwa mem (Li 2013) against a version of CC-4532 v6 that had been hardmasked for TEs and had the Chlamydomonas repeat library appended. Read pairs with at least one read mapped to the *Gypsy-7a_cRei* sequence (included in the repeat library) and a mapping quality score >10 were extracted with samtools view (v1.15) (“-b -h -P -q 10”) (Danecek et al. 2021). The resulting BAM files were used to generate bedgraph files of read coverage using bedtools genomecov v2.30 (“-bg -split”) (Quinlan and Hall 2010). Peaks with <5 reads were filtered out. The resulting tracks were visualized in IGV v2.9.4 (Robinson et al. 2011). Peaks of coverage were manually identified for each strain.

### Phylogenetic analysis of RecQ3 helicases

Peptide sequences were collected by searching for similar proteins to Cre16.g801898, Cre16.g673393, and AT4G35740 using the Phycocosm (Grigoriev et al. 2021), Phytozome (Goodstein et al. 2012) and NCBI databases. Sequences were aligned with MAFFT (v7.305) (Katoh and Standley 2013) through the CIPRES web portal (Miller et al. 2010) and phylogenetic reconstruction was performed using W-IQ-TREE with default parameters (Trifinopoulos et al. 2016). The consensus tree was visualized in iTOL (Letunic and Bork 2019), and the sequences from the subtree representing the RecQ3 subfamily were extracted, realigned, and used to build a RecQ3 phylogeny. The consensus tree was visualized in iTOL.

## SUPPLEMENTAL FILES

**Supplemental Figure S1.** Misassemblies in version 5 and their resolution in version 6.

**Supplemental Figure S2.** Full recombination frequency plots for the estimation of the genetic maps.

**Supplemental Figure S3.** CG methylation and repeat landscape of the CC-1690 assembly.

**Supplemental Figure S4.** Browser view of a v5.6 split gene model merged in CC-4532 v6.1.

**Supplemental Figure S5.** Analyses of coding potential for CC-4532 v6.1.

**Supplemental Figure S6.** Analyses of coding potential for CC-503 v6.1.

**Supplemental Figure S7.** CC-4532 v6 haplotype 2 regions and unique structural mutations.

**Supplemental Figure S8.** Genomic distribution of haplotype 1 and 2 among laboratory strains.

**Supplemental Figure S9.** Summary of indels present at CC-503 reciprocal translocation/inversion double-strand breaks and repair points.

**Supplemental Figure S10.** Browser views of genes at double-strand breaks associated with the CC-503 reciprocal translocation/inversion mutation.

**Supplemental Figure S11.** Browser views of whole-genome re-sequencing data at double-strand breaks associated with the CC-503 reciprocal translocation/inversion mutation.

**Supplemental Figure S12.** Browser view of the CC-503 specific deletion of a prolyl 4-hydroxylase gene.

**Supplemental Figure S13.** Intersect between coding sequence of CC-503 v6.1 gene models and transposable elements.

**Supplemental Figure S14.** Browser view of *DNA-1_cRei* and *CMD1* on chromosome 16.

**Supplemental Table S1.** Summary statistics, gene density and repeat content of the CC-4532 v6 chromosomes.

**Supplemental Table S2.** Summary statistics of the CC-4532 v6 genome split by site class with respect to the CC-4532 v6.1 annotation.

**Supplemental Table S3.** Metrics and sequence context of all CC-4532 v6 assembly gaps.

**Supplemental Table S4.** Putative centromere metrics of the CC-1690 and CC-4532 v6 assemblies.

**Supplemental Table S5.** Comparison of proteomic validation of v5.6 and CC-4532 v6.1 proteins.

**Supplemental Table S6.** Automated and expert annotations of v5.6 and CC-4532 v6.1 structural annotations.

**Supplemental Table S7.** Haplotype 2 regions in CC-4532 v6.

**Supplemental Table S8.** Haplotype 2 coordinates in v5 and CC-4532 v6, and changes between the assembly versions.

**Supplemental Table S9.** Mating type locus R domain genes in CC-4532 v6.1 and CC-503 v6.1.

**Supplemental Table S10.** Curated structural mutations in the CC-503 v6 assembly.

**Supplemental Table S11.** Curated structural mutations in the CC-4532 v6 assembly.

**Supplemental Table S12.** Curated TE insertions/excisions in the CC-503 v6 assembly.

**Supplemental Table S13.** Curated TE insertions/excisions in the CC-4532 v6 assembly.

**Supplemental Table S14.** Approximate coordinates of *Gypsy-7a_cRei* copies among laboratory strains.

**Supplemental Dataset S1.** Master annotation table of CC-4532 v6.1.

**Supplemental Dataset S2.** Gene symbols moved to previous identifiers.

**Supplemental Dataset S3.** Proteins, alignment and phylogeny for RECQ3 analysis.

**Supplemental Dataset S4.** Latest Chlamydomonas repeat library (v3.4).

## ACKNOWLEDGMENTS

The work (proposals: 10.46936/10.25585/60007932, 10.46936/10.25585/60001051, and 10.46936/10.25585/60000843) conducted by the U.S. Department of Energy Joint Genome Institute (https://ror.org/04xm1d337), a DOE Office of Science User Facility, is supported by the Office of Science of The U.S. Department of Energy operated under Contract No. DE-AC02-05CH11231.

We would like to thank the following for their expert advice in assigning descriptions and gene symbols to the v6.1 annotations: Jean Alric, Marius Arend, Ariane Atteia, Olga Baidukova, Steven G. Ball, Matteo Ballotari, Gabriella Benko, Christoph Benning, Robert Bloodgood, Alexandre-Viola Bohne, Pierre Cardol, Yen Peng Chew, Yves Choquet, José L. Crespo, David Dauvillée, Dion Dunford, Susan K. Dutcher, Emilio Fernández-Reyes, Aurora Galván, Michel Goldschmidt-Clermont, Arthur R. Grossman, Patrice P. Hamel, Thomas Happe, Peter Hegemann, Michael Hippler, Martin Jonikas, Steve King, J. Clark Lagarias, Stéphane D. Lemaire, Younghua Li-Beisson, Takuya Matsuo, David Mitchell, Aurora Nedelcu, Jöerg Nickelsen, Adrian Nievergeld, Krishna K. Niyogi, Junmin Pan, Dhruv Patel, Matthew C. Posewitz, Claire Remacle, Nicolas Rouhier, Emanuel Sanz-Luque, Michael Schroda, James Umen, Setsuko Wakao, Florent Waltz, Robert Willows, Felix Willmund, George B. Witman, Francis-Andre Wollman, Katia Wostrikoff, and William Zerges. We would like to thank Julianne Oshiro and Jordan L. Chastain for assistance tracking down PMID accessions for incorporation into Phytozome.

## AUTHOR CONTRIBUTIONS

JWJ, RJC and OV contributed to genome assembly. SS, RJC, SDG, OV and DMG performed gene annotation and post-processing. SDG, JK, JG, KB, CD and YY performed and managed nucleic acids preparation and sequencing. RJC, SDG, OV, PS, CEB, SP, SO and DS performed bioinformatics analyses. SDG, SSM and OV curated gene symbols and contributed annotation. JS and SSM conceived, coordinated and supervised the study. RJC wrote the manuscript with major contributions from SDG, PS, OV and SSM. All authors read and commented on the manuscript.

## DATA AVAILABILITY

CC-4532 v6 is available at Phytozome (https://phytozome-next.jgi.doe.gov). Sequencing reads are available at accessions listed in the text or are in the process of deposition. Work is ongoing to release CC-503 v6 and to incorporate additional browser tracts for the CC-4542 v6 release.

## APPENDIX: GENE SYMBOL NAMING RULES

1) **A gene symbol should communicate the function of that gene’s encoded protein whenever possible**. Gene symbols that describe a phenotype of a mutant at that locus are acceptable, but not preferred, if a function is known. E.g. Cre03.g188250 was previously named *STA6,* because this was the sixth locus identified in a screen for starchless mutants. It has since been demonstrated that this gene encodes the ADP-glucose pyrophosphorylase small subunit, and so it has been renamed *AGP4*.

2) **Gene symbols should never be assigned based solely on the presence of a common domain**. In cases such as the 27 genes that encode proteins with a predicted serine/threonine kinase domain, use the locus ID (CreXX.gNNNNNN) to identify the gene. Only assign a gene symbol once there is some evidence to assign a function to that gene.

3) **Whenever possible, gene symbols should reflect orthologies to genes characterized in other model organism**s. For example, *VKE1* (Cre12.g493150) was renamed *LTO1* to reflect its orthology to a gene with that name in Arabidopsis.

4) **The gene symbol should suggest, but is not required to match, the encoded protein’s name**. For example, the RuBisCO small subunit 1 protein is encoded by *RBCS1* (Cre02.g120100).

5) **A gene symbol for a nuclear gene should consist of three or four capital letters and an Arabic numeral** for the WT allele. In very rare circumstances, as few as two and as many as six letters are allowed if it is necessary to provide continuity to historical data. E.g. *FA2* (Cre07.g351150), *KU80* (Cre10.g423800), *NDUFAF6* (Cre03.g194300) and *RAPTOR1* (Cre08.g371957). We note here that previous efforts to assign gene symbols in Chlamydomonas applied a more stringent three-character only rule (Dutcher 1995). We argue that this rule should be relaxed, since three-character gene symbols often obscure orthologous relationships to genes in other species (e.g. *CRI1* (Cre16.g651923) was renamed *CRTISO1* to highlight its orthology to that gene in other species). More importantly, the insistence on three letters has created several naming collisions that were corrected in v6.1. For example, the “COP” in *COP1* (Cre02.g085050) stands for “constitutive photomorphogenesis protein”, while the same three letters are used for the Chlamyopsin genes like *COP2*. In v6.1, Cre02.g085050 was renamed *PCOP1* for “Plant-type constitutive photomorphogenesis protein”, thus preserving “COP” for the Chlamyopsin genes. Similarly, in v5.6 “MCP” was used for both mitochondrial substrate carrier proteins and Molybdenum cofactor carrier proteins.

6) **Mutant alleles are indicated by the same characters as the WT gene, but in lower case, followed by a dash and an allele number**. E.g. *arg7-1* and *arg7-8* for two mutant alleles of *ARG7* (Cre01.g021251). Because a dash is used to signify a mutant allele, dashes must never be used as part of a WT gene symbol.

7) **Gene symbols for organelle-expressed genes consist of three lower case letters and one upper case letter.** E.g., CreCp.g802313, which encodes the large subunit of RuBisCO has *rbcL* for a gene symbol. This follows the naming convention of prokaryotes.

8) **All genes get a number to distinguish members of a gene family**. This is applied universally, even to those that have not had one historically. E.g. in v6.1, *METC* (Cre16.g669550) has become *METC1, PSAD* (Cre05.g238332) is now *PSAD1*. This is not simply a matter of fastidiously applying a rigid format. Including a number, typically starting with “1”, helps to future-proof the annotations. Since 76% of genes still lack a gene symbol, any gene symbol that is currently one-of-a-kind may in the future be found to be part of a larger family. This is especially true when the Chlamydomonas gene symbols are used to help guide the annotations in other species, which may have multiple co-orthologous genes.

9) **Sequential letters should be used in place of numbers for symbols that end in a number.** For some gene families, it is informative to have a gene symbol root that ends in an Arabic numeral. For example, Cre09.g415450 and Cre13.g586500 both encode kinesin-17 family members; called *KIN17-1* and *KIN17-2* in v5.6. Unfortunately, this approach is incompatible with using a dash and Arabic numeral exclusively to identify mutant alleles (see rule 5 above). To resolve this, the dash and numeral were replaced with sequential letters in v6.1; in this case, *KIN17A* and *KIN17B*. The same strategy was used for gene symbols with a period. E.g. The *NRT2.4* (Cre03.g150101) and *NRT2.5* (Cre03.g150151) genes were renamed *NRT2D* and *NRT2E*, respectively. Note that in this case, the protein names remain unchanged as NRT2.4 and NRT2.5 (Léran et al. 2014).

10) **Gene symbols may not contain any other punctuation marks (periods, dashes, colons, etc.), Greek letters, subscripts or superscripts.** The use of non-Unicode characters (e.g. Greek letters) can frequently wreak havoc with the tools used in computational biology. If necessary, Greek letters should be replaced with the closest Roman equivalent.

11) **Gene symbols should not include “Chlamydomonas”**. E.g. Cre02.g092150 had been named *CSL* in v5.6 for “Chlamydomonas SHOC2/SUR8-like LRR”, but this symbol was demoted to alias in v6.1. Refraining from using the genus or species in the gene symbol helps preserve the universality of symbols when used to guide the annotation of other species.

12) **Make sure any new gene symbol roots are not used for another gene family**. When the function has been identified for a previously unnamed gene, try to copy or adapt the gene symbol from orthologs in other model species. When the gene’s function is novel, choose a communicative three to four letter root. Make sure that root is not in use in Chlamydomonas or other model species. We recommend checking uniprot.org for this purpose.

## Notes

### Competing Interest Statement

The authors have declared no competing interest.

