## Supplemental Figures for "The Chlamydomonas Genome Project, version 6: reference assemblies for mating type *plus* and *minus* strains reveal extensive structural mutation in the laboratory"

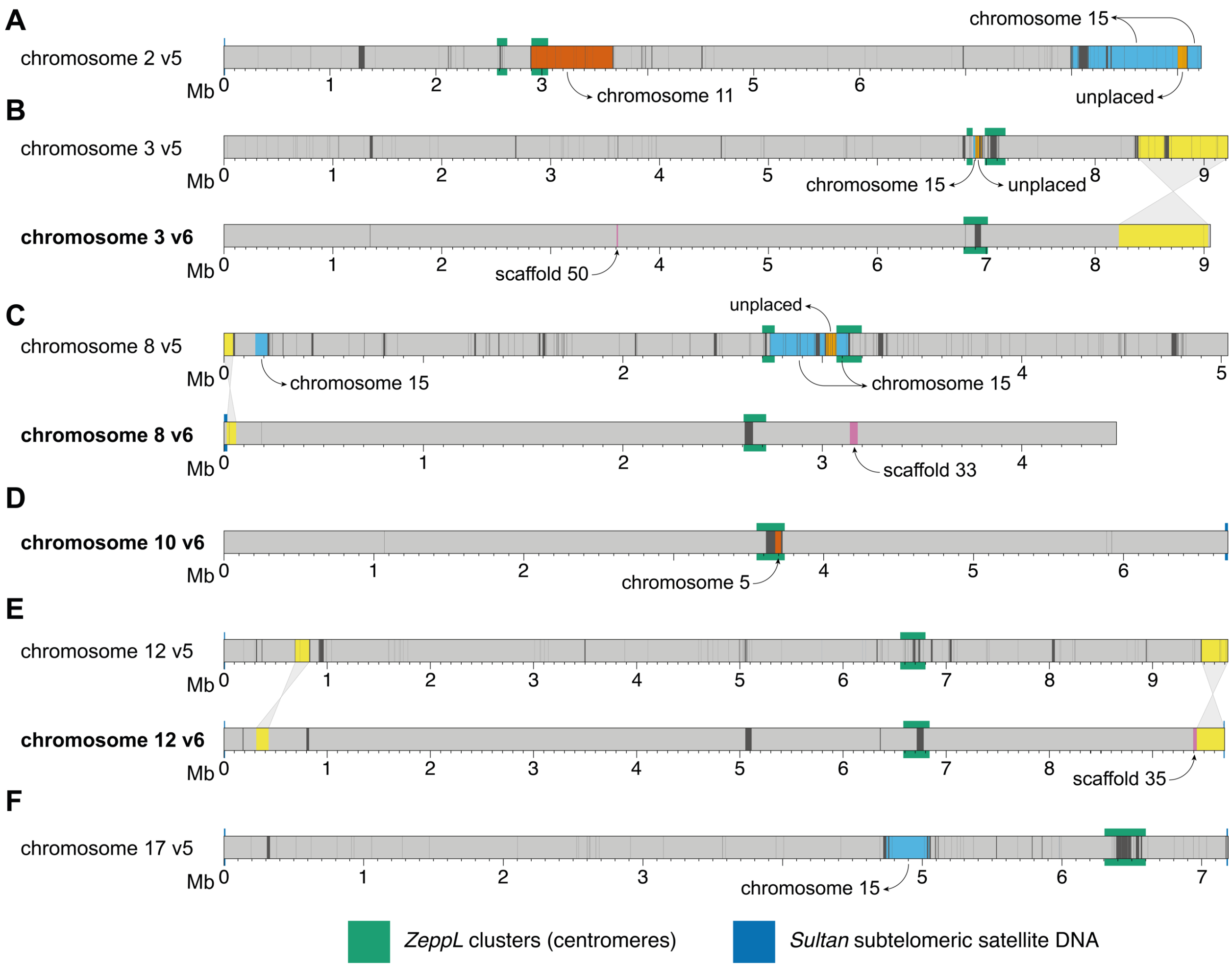

**Supplemental Figure S1. Misassemblies in version 5 and their resolution in version 6.**
(Supports Figures 1 and 2).
Overview of major chromosomal changes between the assemblies, note that several small local corrections are not shown. CC-503 v6 chromosomes are shown to simplify mapping to v5. Chromosomes 5 and 11 are shown in Figure 2, and changes to chromosome 9 in v6 are shown in Figure 7.
**(A)** Chromosome 2 (see Figure 7 for sequence gained in v6). Note two centromeric regions in v5.
**(B)** Chromosome 3. Note two centromeric regions in v5.
**(C)** Chromosome 8. Note two centromeric regions in v5.
**(D)** Chromosome 10 (v5 not shown since only v6 change is the movement of a single sequence from chromosome 5).
**(E)** Chromosome 12.
**(F)** Chromosome 17 (v6 not shown since only change is the movement of a single sequence to chromosome 15).

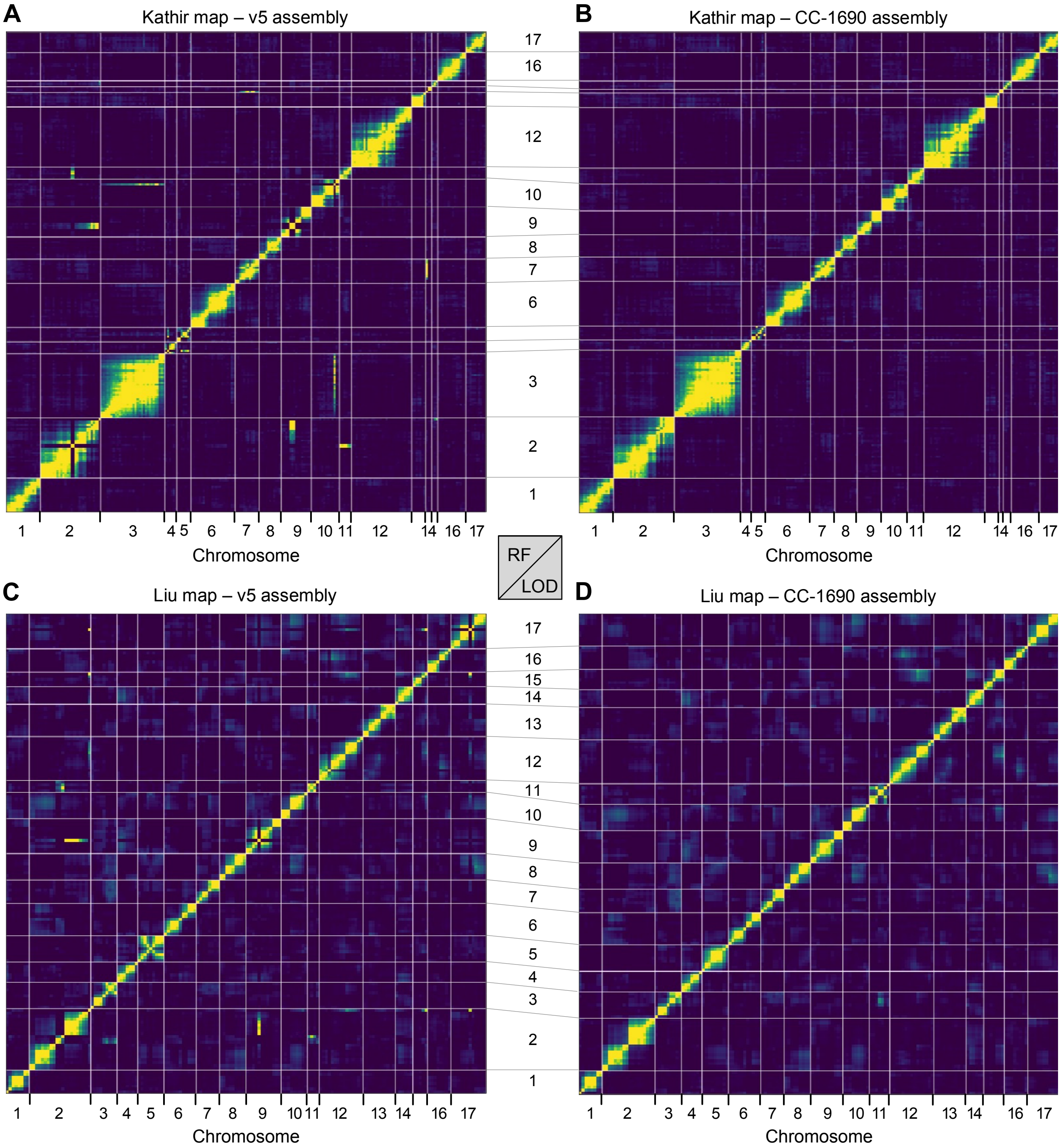

**Supplemental Figure S2. Full recombination frequency plots for the estimation of the genetic maps.** (Supports Figure 2).

**(A, B)** Full RF plots for the molecular markers used by Kathir et al. (2003) with the genomic coordinates from the v5 **(A)** or CC-1690 **(B)** assemblies.

**(C, D)** Full RF plots for the SNPs extracted from Liu et al. (2018) with the genomic coordinates from the v5 **(C)** or CC-1690 **(D)** assemblies.

Strong linkage is indicated by a yellow color; absence of linkage is shown as dark blue. RF: recombination fraction. LOD: logarithm of the odds.

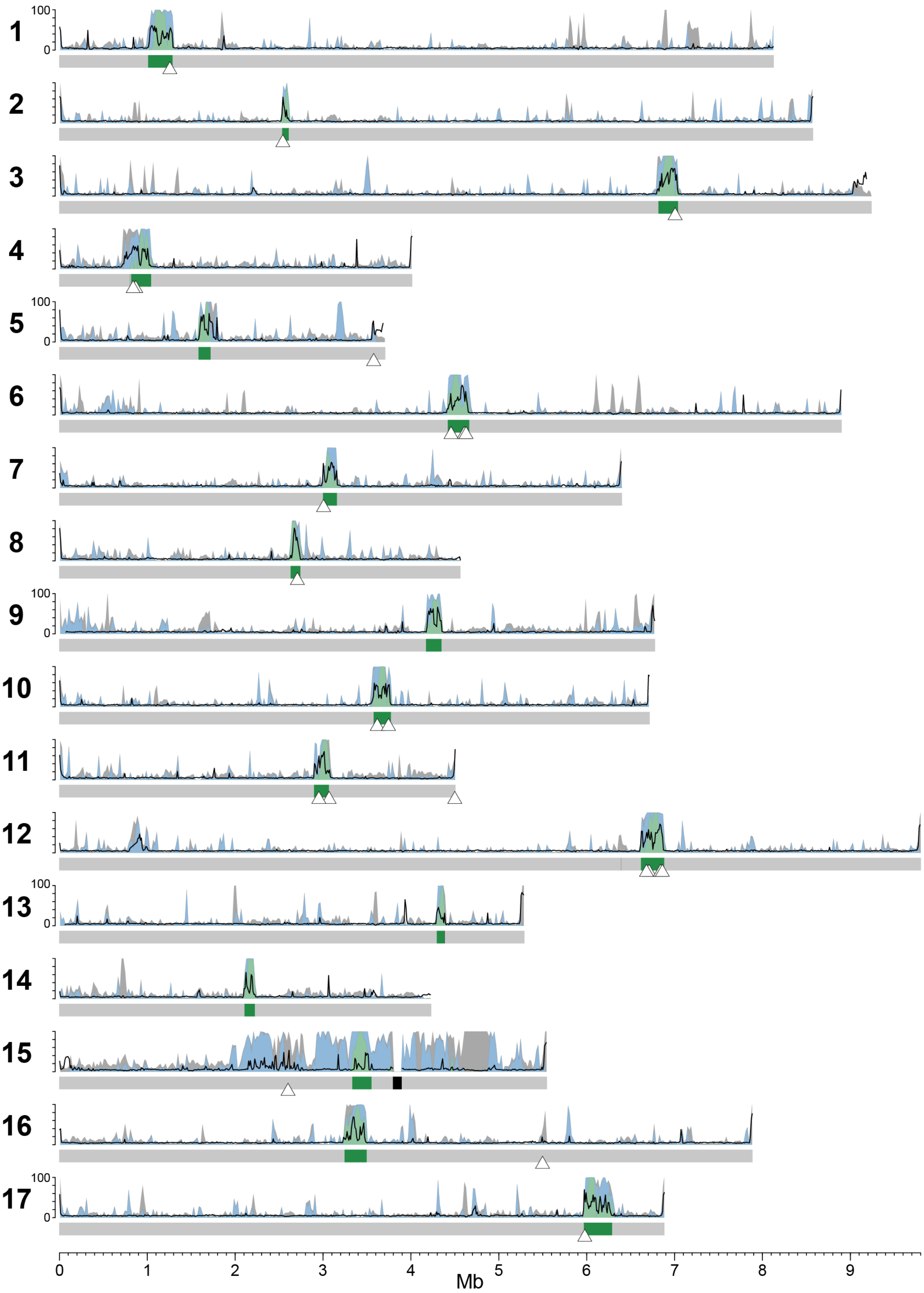
100
0
0
1
2
3
4
5
6
7
8
9
Mb

**Supplemental Figure S3. CG methylation and repeat landscape of the CC-1690 assembly.**
(Supports Figure 1).
Each gray bar represents a CC-1690 chromosome with the putative centromeres colored green and assembly gaps black. The line graph represents CG methylation (0-100%) in 10 kb windows. The stacked density plot represents repeat content in 20 kb windows, green represents *ZeppL-1_cRei*, blue any other TE, and gray any tandem repeat (microsatellites and satellites). White triangles represent the hypermethylated regions previously identified by Lopez et al. (2015).

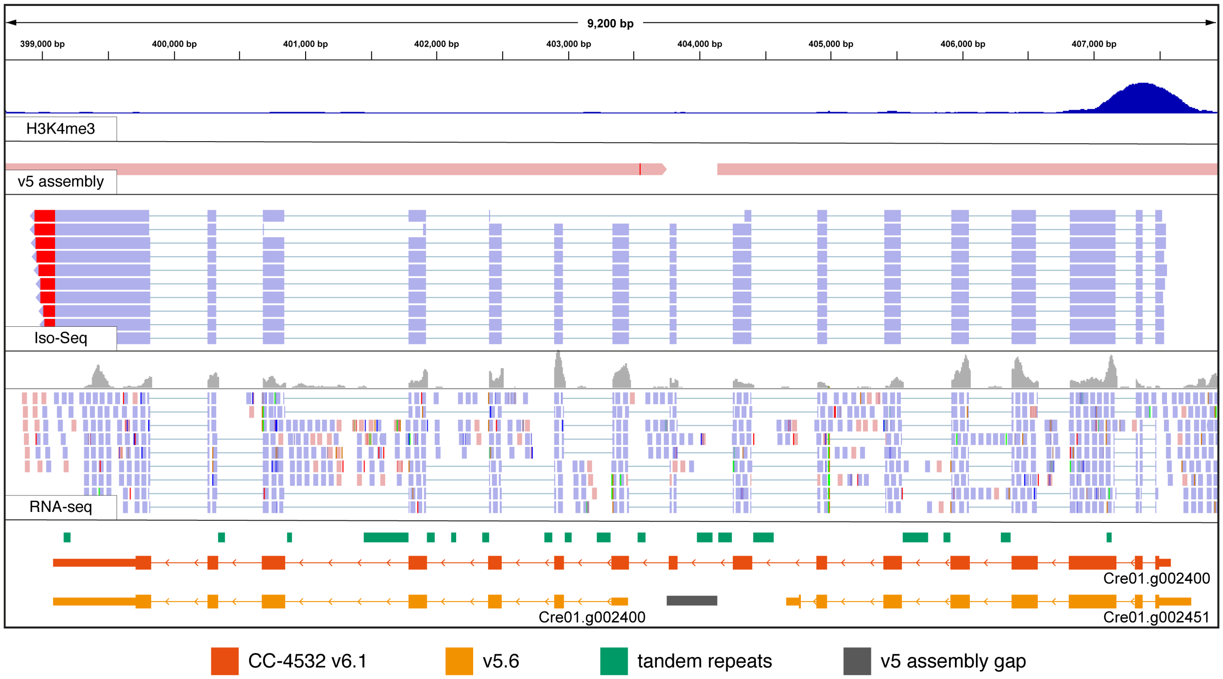

**Supplemental Figure S4. Browser view of a v5.6 split gene model merged in CC-4532 v6.1.** (Supports Figure 5).

Cre01.g002400 is a putative triglyceride lipase homologous to *HIL1* (*HEAT INDUCIBLE LIPASE1*) in *Arabidopsis thaliana*. H3K4me3 ChIP-seq marks promoters. The v5 assembly track shows an alignment of v5 contigs to CC-4532 v6, with the assembly gap appearing as an unmapped region. Coordinates for v5.6 gene models (orange) were lifted over to CC-4532 v6. Thick blocks represent coding sequence, thin blocks UTRs, and conjoining lines introns. Red mismatches at the end of Iso-Seq reads correspond to poly(A) tails.

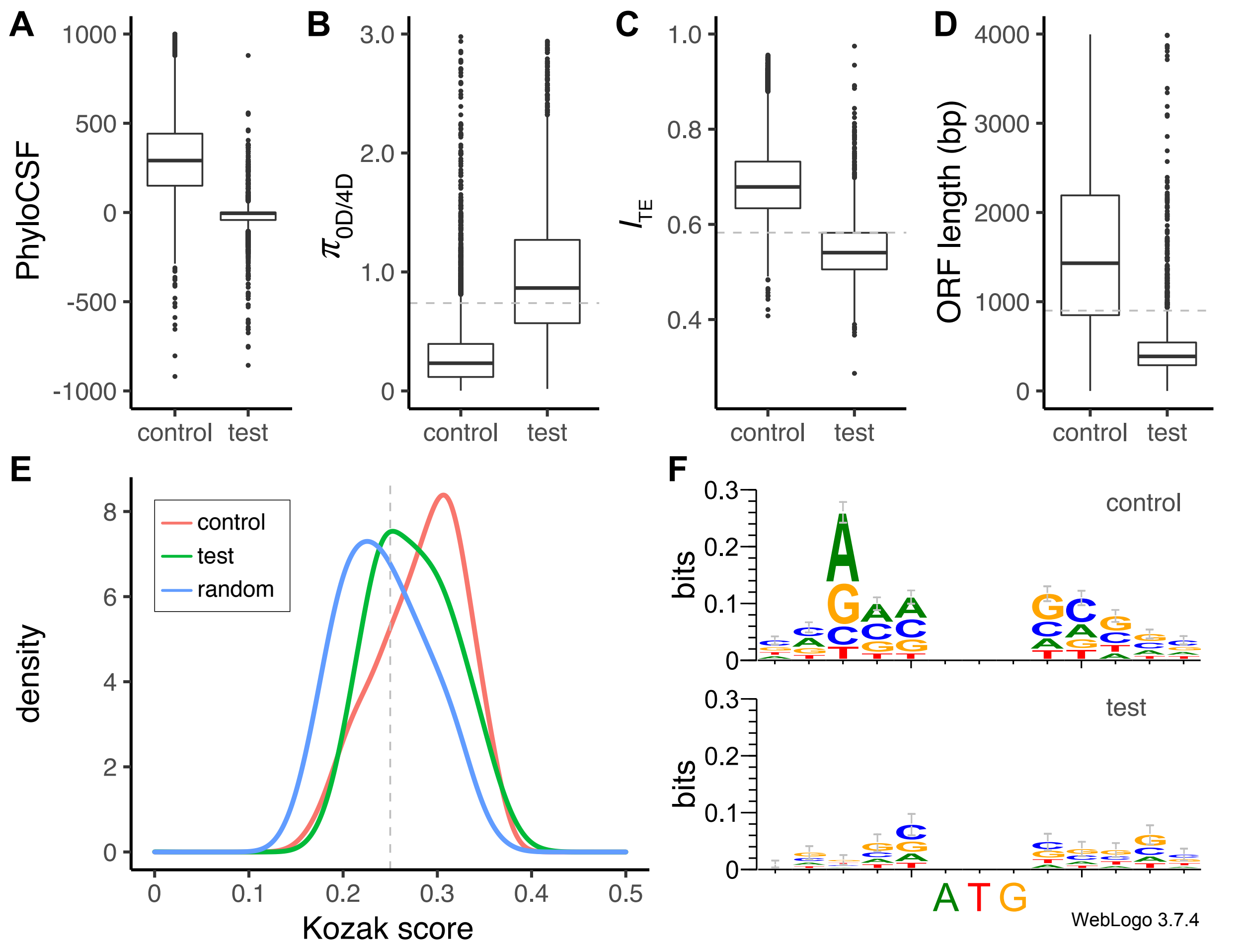

**Supplemental Figure S5. Coding potential analyses for CC-4532 v6.1.**
(Supports Table 2).
Genes from the preliminary annotation of CC-4532 were split to control (algal homolog or functional domain, N = 15,237) or test (no homolog/domain, N = 2,362). See Methods for further details.
**(A)** PhyloCSF scores for control and test set genes. Scores were calculated based on an 8-species whole-genome alignment. Test set genes scoring <1 failed (N = 1,982).
**(B)** Ratio of genetic diversity ($\pi$) at zero-fold and four-fold degenerate sites, based on whole-genome re-sequencing data of 17 Chlamydomonas Quebec field isolates. Test set genes with ratios >0.738 (the 95$^{th}$ percentile of control set) failed (N = 1,708).
**(C)** Codon usage as quantified by the index of translation elongation ($I_{TE}$). Test set genes with values <0.582 (the 5$^{th}$ percentile of control set) failed (N = 1,776).
**(D)** ORF length excluding any microsatellite or satellite DNA. Test set genes with ORFs <900 bp (N = 2,154) that failed an appropriate number of other tests were included in the low coding potential set.
**(E)** Kozak scores for the 5 bp up and downstream of start codons, calculated by comparison to a reference Kozak-like sequence. Control distribution was estimated from the half of the control set genes not used to generate the reference, and the random distribution was generated from 10,000 random sequences with an expected GC content ~64%. Test set genes with scores <0.25 failed (N = 1,379).
**(F)** Kozak sequence logos produced using WebLogo 3 (Crooks et al. 2004). Control logo was generated from a random half of control set genes and used as the reference for generating the distributions above.

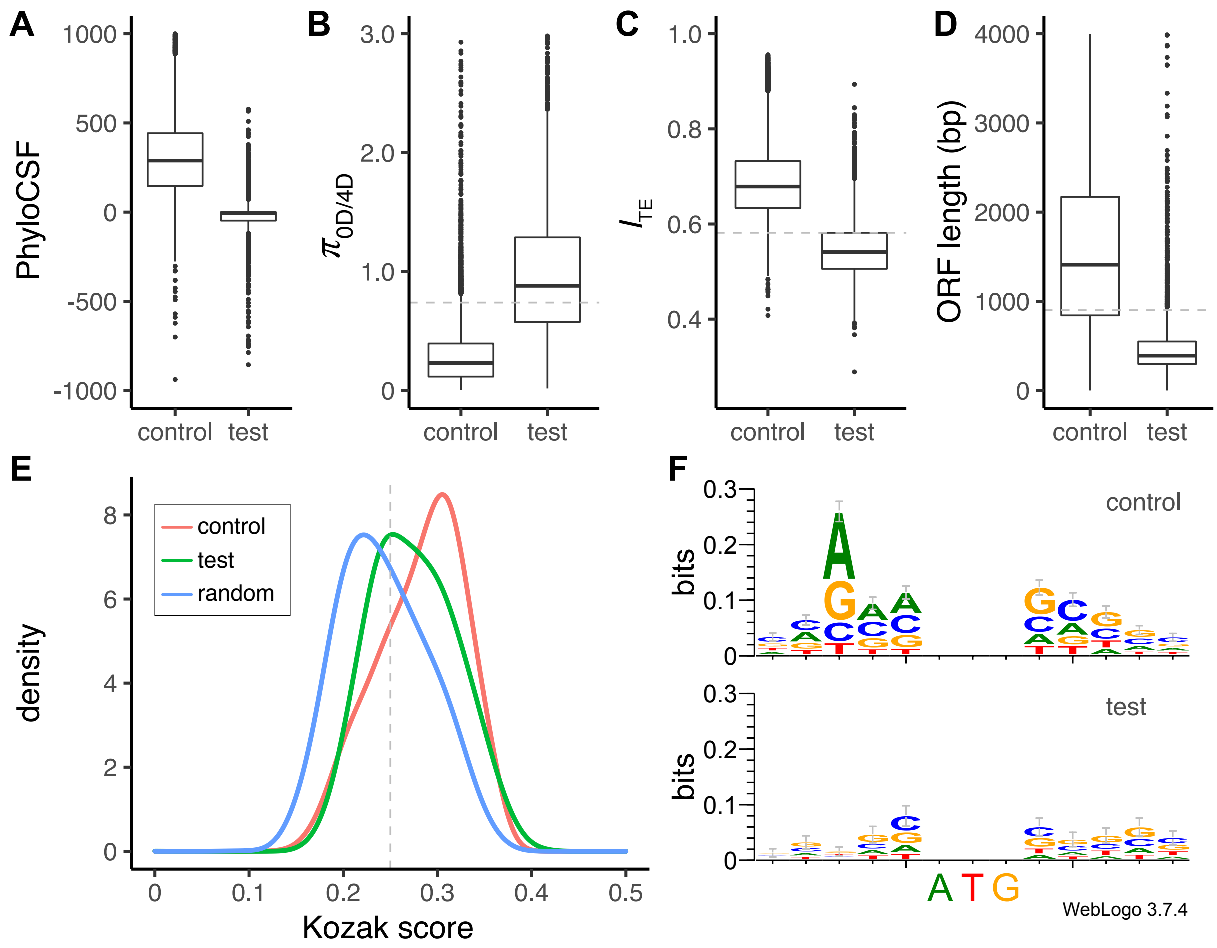

**Supplemental Figure S6. Coding potential analyses for CC-503 v6.1.**
(Supports Table 2).
Genes from the preliminary annotation of CC-503 were split to control (algal homolog or functional domain, N = 15,159) or test (no homolog/domain, N = 2,418). See Methods for further details.
**(A)** PhyloCSF scores for control and test set genes. Scores were calculated based on an 8-species whole-genome alignment. Test set genes scoring <1 failed (N = 2,022).
**(B)** Ratio of genetic diversity (π) at zero-fold and four-fold degenerate sites, based on whole-genome re-sequencing data of 17 Chlamydomonas Quebec field isolates. Test set genes with ratios >0.738 (the 95th percentile of control set) failed (N = 1,760).
**(C)** Codon usage as quantified by the index of translation elongation ($I_{TE}$). Test set genes with values <0.582 (the 5th percentile of control set) failed (N = 1,815).
**(D)** ORF length excluding any microsatellite or satellite DNA. Test set genes with ORFs <900 bp (N = 2,185) that failed an appropriate number of other tests were included in the low coding potential set.
**(E)** Kozak scores for the 5 bp up and downstream of start codons, calculated by comparison to a reference Kozak-like sequence. Control distribution was estimated from the half of the control set genes not used to generate the reference, and the random distribution was generated from 10,000 random sequences with an expected GC content ~64%. Test set genes with scores <0.25 failed (N = 1,407).
**(F)** Kozak sequence logos produced using WebLogo 3 (Crooks et al. 2004). Control logo was generated from a random half of control set genes and used as the reference for generating the distributions above.

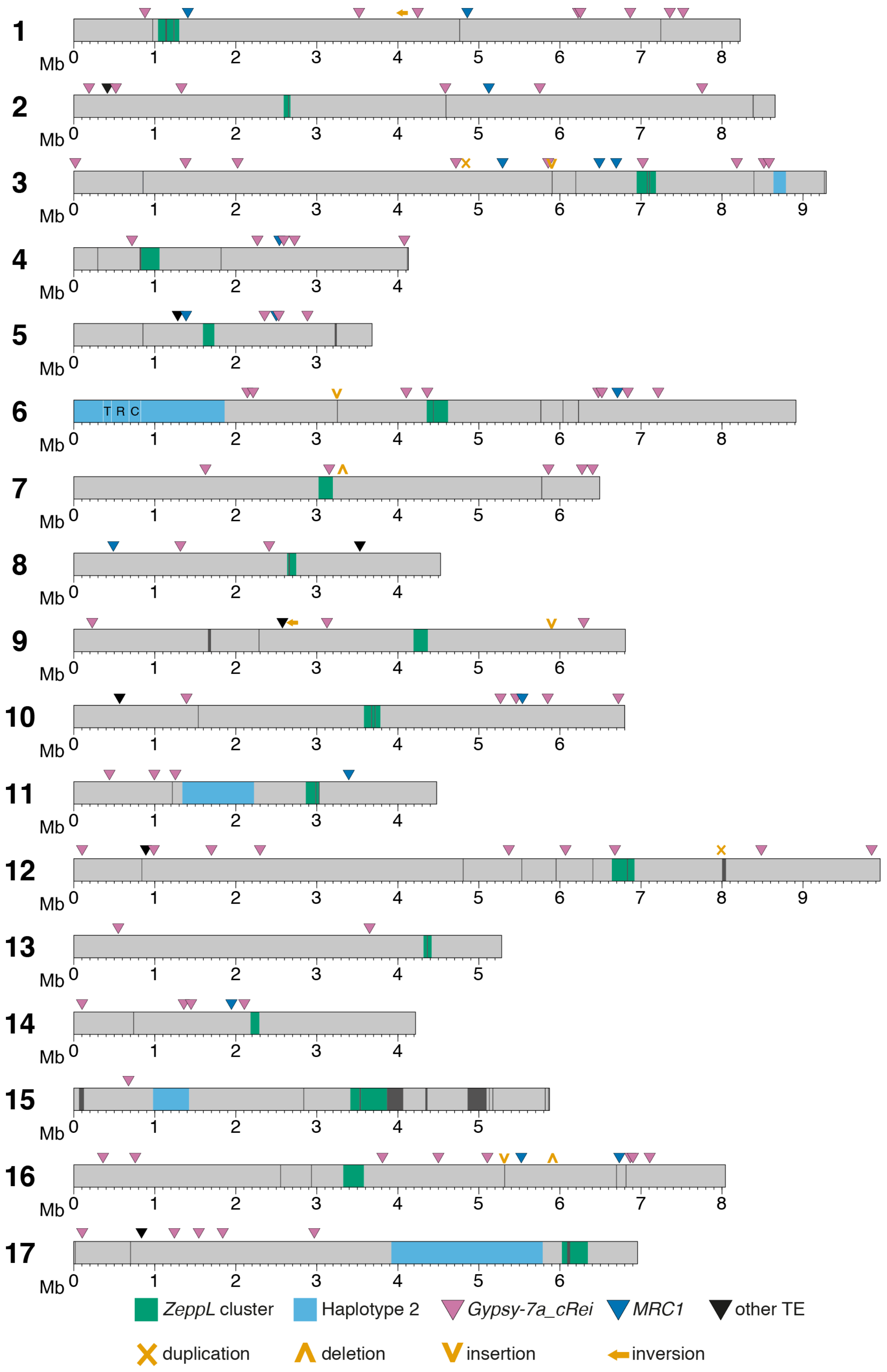
Mb
T R C
ZeppL cluster
Haplotype 2
Gypsy-7a_cRei
MRC1
other TE
duplication
deletion
insertion
inversion

**Supplemental Figure S7. CC-4532 v6 haplotype 2 regions and unique structural mutations.** (Supports Figure 1 & 8).

Overview of the CC-4532 v6 chromosomes, showing putative centromeres (*ZeppL* clusters), haplotype 2 regions, assembly gaps (dark grey regions), de novo TE insertions and structural mutations. See Supplemental Tables 11 and 13 for TE and mutation coordinates, respectively. The three domains (T, R and C) of the *MT*$^{-}$ locus are marked by white lines within the haplotype 2 region of chromosome 6.

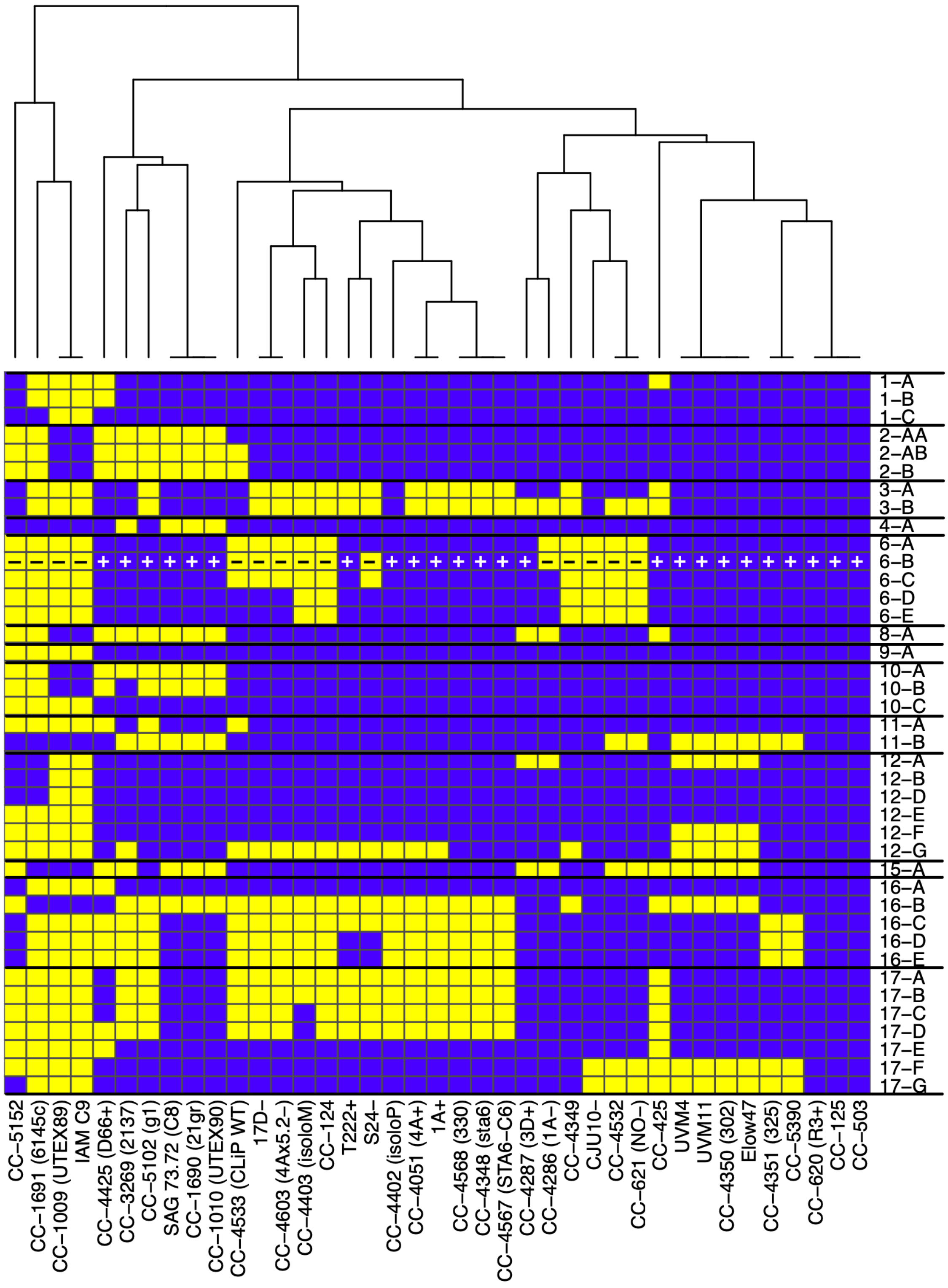

**Supplemental Figure S8. Genomic distribution of haplotype 1 and 2 among laboratory strains.** Each cell represents a block of sequence that is either haplotype 1 (blue) or 2 (yellow) in each strain, with chromosomes shown on the y-axis. Note that the cells are not to scale, for genomic coordinates of each block and changes relative to v5 see Supplemental Table 8.

| Relative to: | Double-strand break / repair event | Indel | Affected genes |
|---|---|---|---|
| CC-1690 / CC-4532 v6 | DSB1 | 182 bp deletion | Cre09.g390100 |
| | DSB2 | 1,950 bp deletion | Cre09.g393284 |
| | DSB3 | 62 bp deletion | Cre09.g387023 |
| CC-503 v6 | repair | 2 bp insertion | |
| | repair | 14 bp insertion | |
| | repair | 56 bp insertion | |

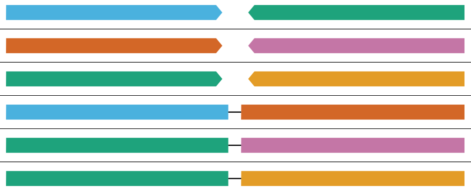

**Supplemental Figure S9. Summary of indels present at CC-503 reciprocal translocation/inversion double-strand breaks and repair points.**
(Supports Figure 7).
Each DSB was associated with a deletion, and each repair with an insertion. For DSBs, CC-4532 v6.1 gene IDs are listed to represent the predicted non-mutant genes.

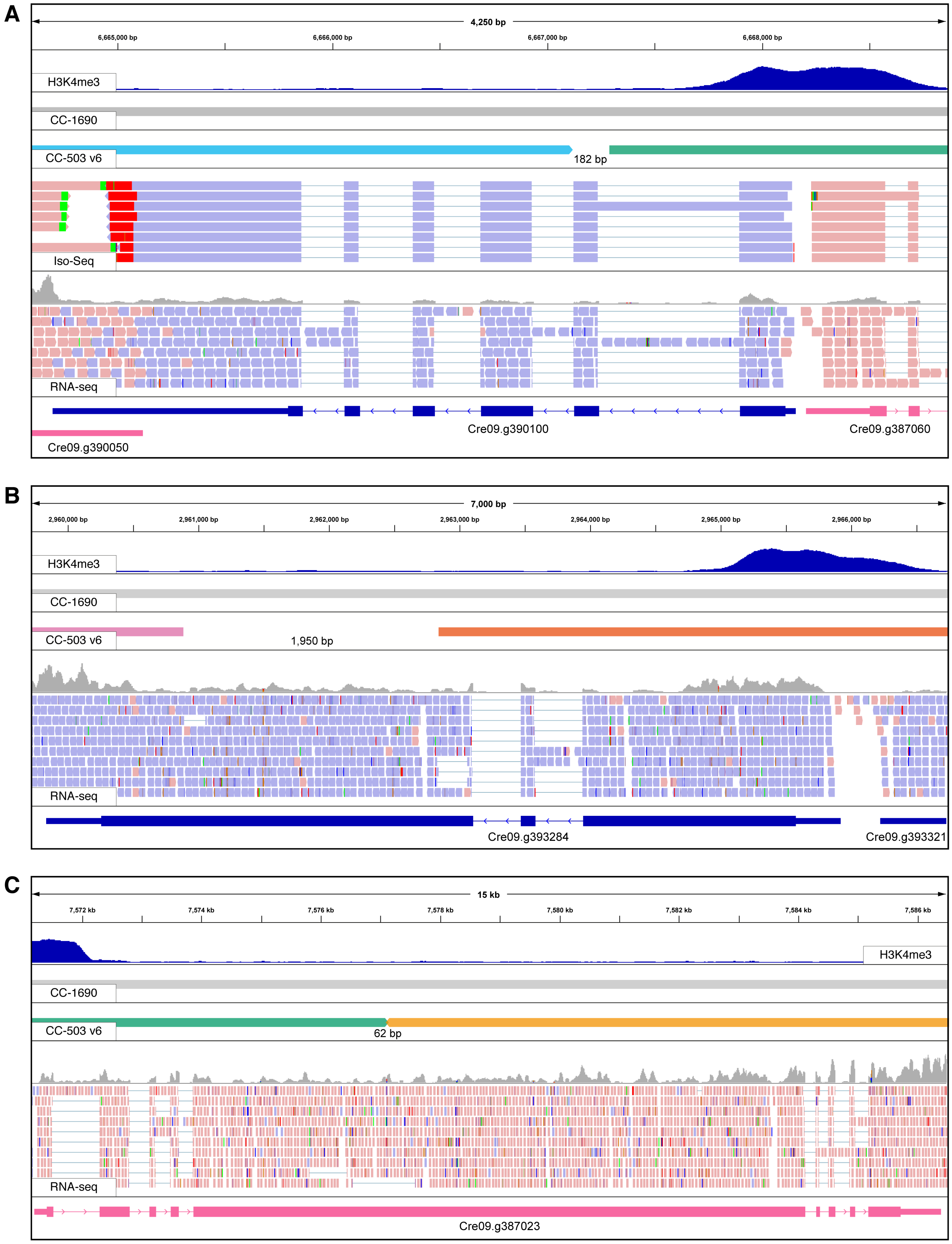
A
4,250 bp
6,665,000 bp
6,666,000 bp
6,667,000 bp
6,668,000 bp
H3K4me3
CC-1690
CC-503 v6
182 bp
Iso-Seq
RNA-seq
Cre09.g390100
Cre09.g387060
Cre09.g390050
B
7,000 bp
2,960,000 bp
2,961,000 bp
2,962,000 bp
2,963,000 bp
2,964,000 bp
2,965,000 bp
2,966,000 bp
H3K4me3
CC-1690
CC-503 v6
1,950 bp
RNA-seq
Cre09.g393284
Cre09.g393321
C
15 kb
7,572 kb
7,574 kb
7,576 kb
7,578 kb
7,580 kb
7,582 kb
7,584 kb
7,586 kb
H3K4me3
CC-1690
CC-503 v6
62 bp
RNA-seq
Cre09.g387023

**Supplemental Figure S10. Browser views of genes at double-strand breaks associated with the CC-503 reciprocal translocation/inversion mutation.**
(Supports Figure 7).
Genomic regions are shown relative to the CC-4532 v6.1 assembly (i.e. the ancestral state). Data tracts from top to bottom are: i) H3K4me3 ChiP-seq data marking promoters, ii), alignment of CC-1690 assembly, iii) alignment of CC-503 v6 assembly (colours match Figure 7 and deletion lengths are labelled), iv) Iso-Seq (only **A**, no full-length reads for **B**, **C**), v) RNA-seq, vi) CC-4532 v6.1 gene models.
**(A)** DSB1, coordinates are chromosome 2: 6,664,000 - 6,668,876 bp.
**(B)** DSB2, coordinates are chromosome 9: 2,959,720 - 2,966,763 bp.
**(C)** DSB3, coordinates are chromosome 2: 7,571,150 - 7,586,550 bp.

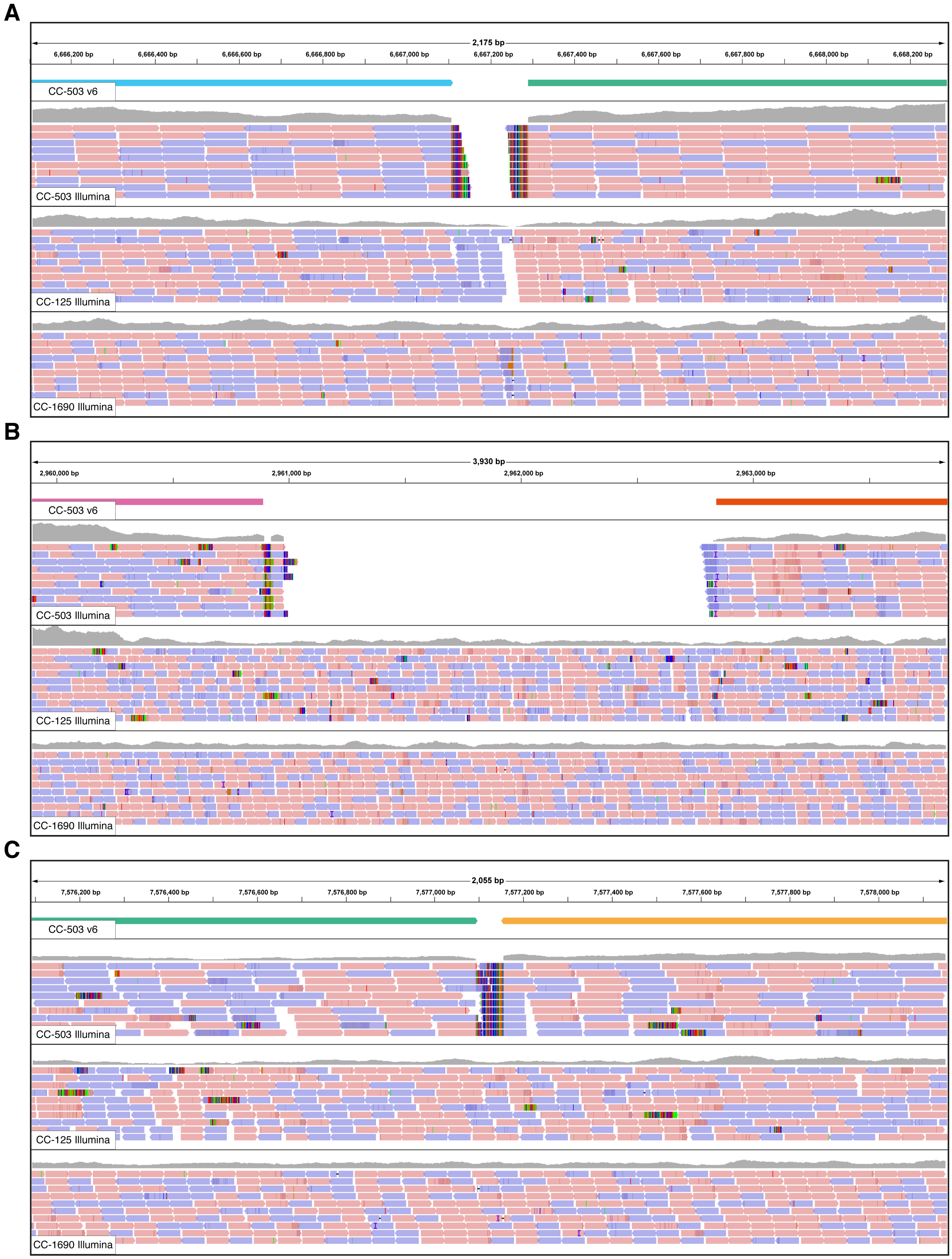
A
2,175 bp
6,666,200 bp
6,666,400 bp
6,666,600 bp
6,666,800 bp
6,667,000 bp
6,667,200 bp
6,667,400 bp
6,667,600 bp
6,667,800 bp
6,668,000 bp
6,668,200 bp
CC-503 v6
CC-503 Illumina
CC-125 Illumina
CC-1690 Illumina
B
3,930 bp
2,960,000 bp
2,961,000 bp
2,962,000 bp
2,963,000 bp
CC-503 v6
CC-503 Illumina
CC-125 Illumina
CC-1690 Illumina
C
2,055 bp
7,576,200 bp
7,576,400 bp
7,576,600 bp
7,576,800 bp
7,577,000 bp
7,577,200 bp
7,577,400 bp
7,577,600 bp
7,577,800 bp
7,578,000 bp
CC-503 v6
CC-503 Illumina
CC-125 Illumina
CC-1690 Illumina

**Supplemental Figure S11. Browser views of whole-genome re-sequencing data at double-strand breaks associated with the CC-503 reciprocal translocation/inversion mutation.**
(Supports Figure 7).
Genomic regions are shown relative to the CC-4532 v6.1 assembly (i.e. the ancestral state). Data tracts from top to bottom are: i) alignment of CC-503 v6 assembly (colors match Figure 7), ii), alignment of CC-503 Illumina re-sequencing data, iii) alignment of CC-125 Illumina re-sequencing data, iv) alignment of CC-1690 Illumina re-sequencing data.
**(A)** DSB1, coordinates are chromosome 2: 6,666,106 - 6,668,290 bp.
**(B)** DSB2, coordinates are chromosome 9: 2,959,894 - 2,963,841 bp.
**(C)** DSB3, coordinates are chromosome 2: 7,576,093 - 7,578,156 bp.

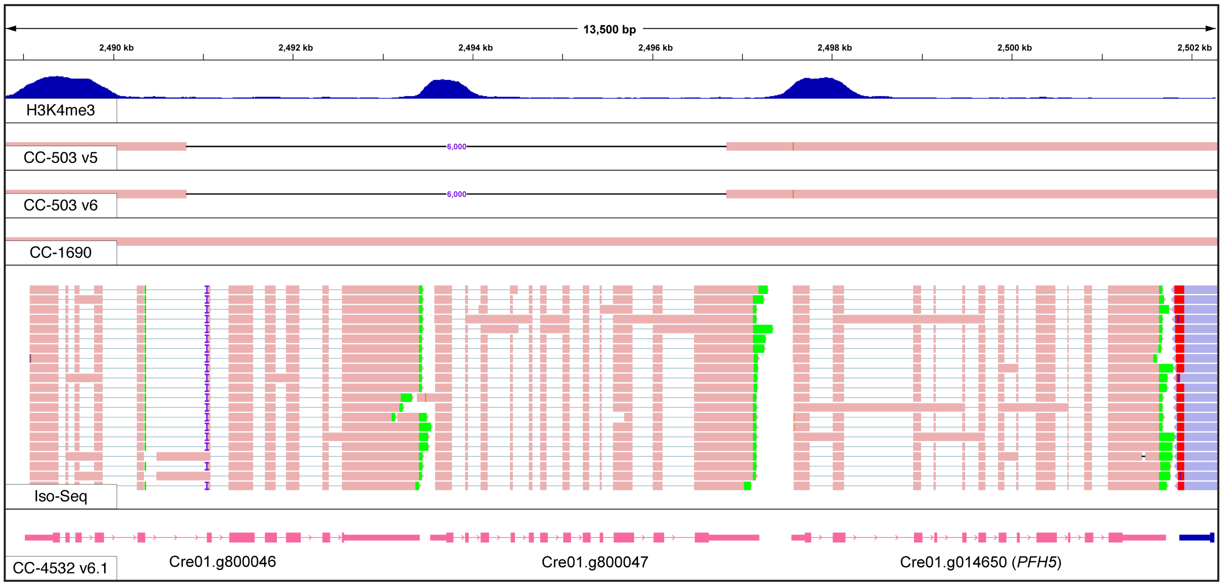

**Supplemental Figure S12. Browser view of the CC-503 specific deletion of a prolyl 4-hydroxylase gene.**
(Supports Figure 7).
Genomic regions are shown relative to the CC-4532 v6.1 assembly (i.e. the ancestral state). Data tracts from top to bottom are: i) H3K4me3 ChiP-seq data marking promoters, ii), alignment of v5 assembly, iii) alignment of CC-503 v6 assembly, iii) alignment of CC-1690 assembly, iv) Iso-Seq, v) CC-4532 v6.1 gene models. Coordinates are chromosome 1: 2,488,800 – 2,502,300 bp.

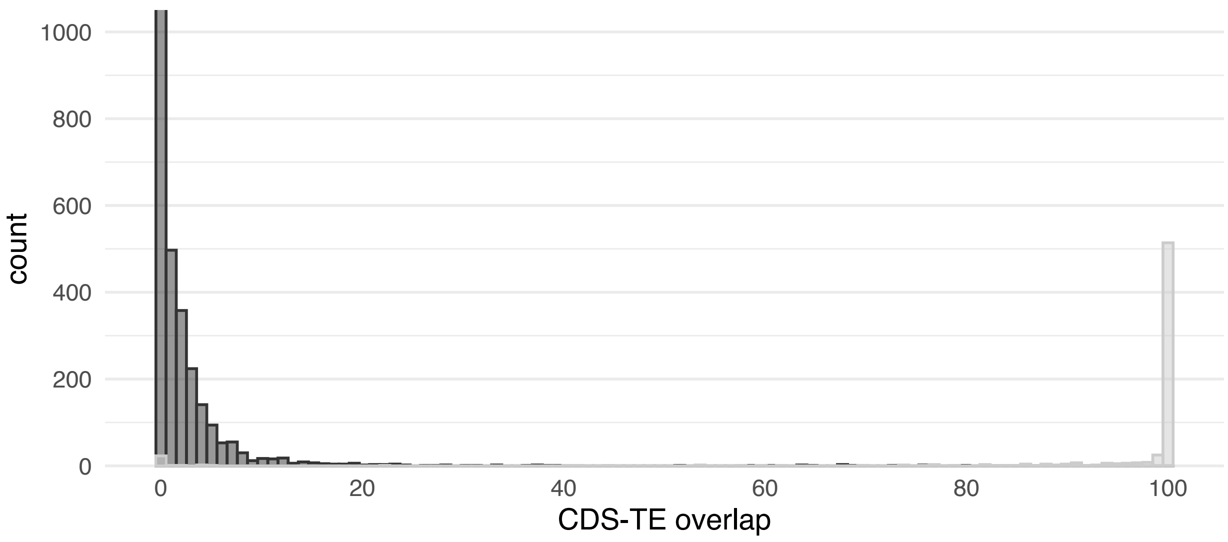

**Supplemental Figure 13. Intersect between coding sequence of CC-503 v6.1 gene models and transposable elements.**
(Supports Figure 9).
Per gene overlap between coding sequence (CDS) and TE sequence for non-TE and TE genes. Note that not all non-TE genes are shown since >>1,000 have 0% overlap with TEs.

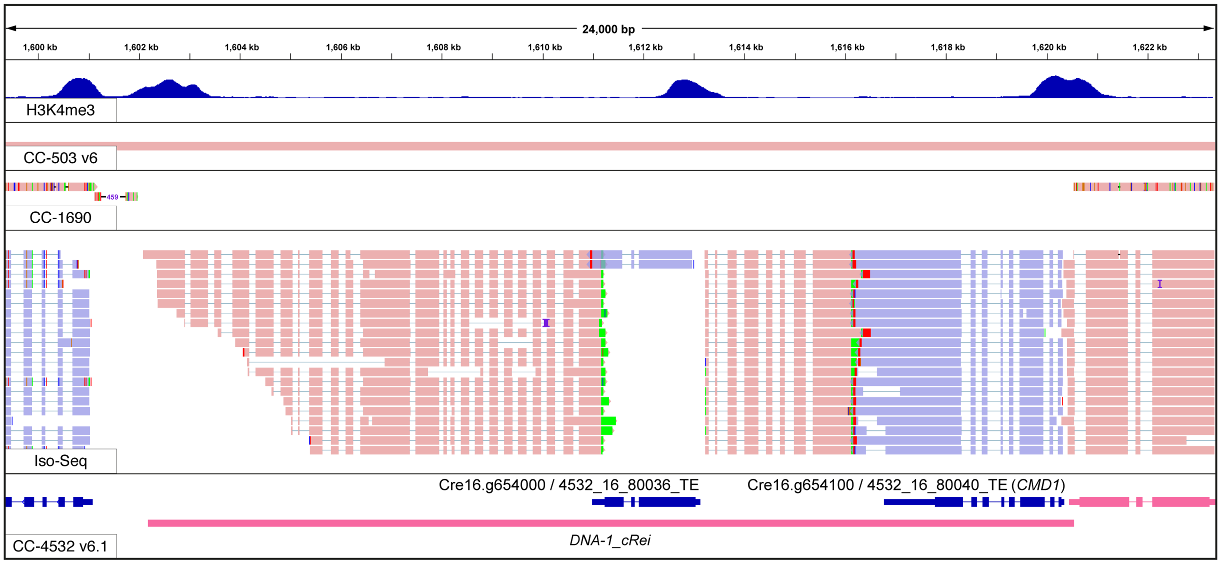

**Supplemental Figure 14. Browser view of *DNA-1_cRei* and *CMD1* on chromosome 16.**
(Supports Figure 9).
Genomic regions are shown relative to the CC-4532 v6.1 assembly. Data tracts from top to bottom are: i) alignment of CC-503 v6 assembly, ii), alignment of CC-1690 assembly, iii) Iso-Seq, iv) CC-4532 v6.1 gene models and repeat annotation. Note CC-1690 is haplotype 2 in this region and that the entire copy of *DNA-1_cRei* is absent. Also note that only two of the *DNA-1_cRei* genes are included in the CC-4532 v6.1 annotation, the predicted proteins from the excluded two genes did not have a hit to a known TE protein or contain any recognised domains. Coordinates are chromosome 16: 1,599,350-1,623,350 bp.
